# Identification of protein secretion systems and type III effectors in wood-associated bacteria of the genus *Xylophilus*

**DOI:** 10.64898/2026.02.05.703958

**Authors:** Zoe Roux, Naama Wagner, Laurent Brottier, Tal Pupko, Ralf Koebnik

**Affiliations:** PHIM, Université de Montpellier, Cirad, INRAE, Institut Agro, IRD, Montpellier, France; Shmunis School of Biomedicine and Cancer Research, Tel Aviv University, Israel

## Abstract

To shed light into the biology of bacteria belonging to the genus *Xylophilus*, including the grapevine pathogen *Xylophilus ampelinus*, we scrutinized all available genomes of the genus for the presence of type III secretion and flagellar systems. We found three different flagellar systems in the genus, one of which was present in all twelve strains with good-quality genome sequences were available. The other two flagellar systems were only detected in one or two strains. We also identified two types of type III secretion systems, likely under control of the AraC-type transcriptional activator HrpX. One system with resemblance to systems from plant-pathogenic bacteria was only found in the grapevine pathogen. The other system was found in three strains of *Xylophilus*, all isolated from plant material. We predicted genes that are co-regulated with the type III secretion systems, as supported by the presence of strongly conserved HrpX-binding promoter elements. We identified about 40 type III effectors in the grapevine pathogen with homologs in plant pathogenic bacteria. In contrast, a rhododendron flower isolate had only two type III effector gene candidates with conserved HrpX-binding promoter elements but many genes without homologs beyond the species. Finally, we predicted and confirmed three novel effector candidates from *X. ampelinus* to contain a functional type III secretion signal using an AvrBs1 reporter approach. The presence of type III effectors suggests that effector-triggered immunity may exist in grapevine or non-host plants and that strategies targeting type III effectors for resistance engineering may contribute to suitable control measures.

## INTRODUCTION

*Xylophilus* is an understudied bacterial genus within the order *Burkholderiales*, family *Comamonadaceae*. These bacteria are primarily known for their association with plants, particularly with woody tissues, which is reflected in the genus name (from Greek: *xýlon* – wood; philus – friend, loving) (https://lpsn.dsmz.de/, accessed on 08/07/2025; Freese et al. 2026). For instance, the type strain of the genus, belonging to the species *Xylophilus ampelinus* (formerly known as *Xanthomonas ampelina*), has been isolated from a diseased grapevine plant (Willems et al. 1987a, 1987b). Bacteria belonging to another proposed species, *Xylophilus rhododendri*, have been isolated from the flowers of a *Rhododendron* plant (Lee et al. 2020). Other *Xylophilus* bacteria without species designation have been identified in metagenomic samples from woody tissues, e.g., from almond and blueberry (Anguita-Maeso et al. 2022; Chacón et al. 2022).

*Xylophilus* bacteria are of particular interest to plant pathologists and viticulturists due to their potential to damage valuable crops. One of the most notable species in this genus is the pathogen*X. ampelinus*, causing bacterial necrosis in grapevines (*Vitis vinifera*). Symptoms include discoloration of young shoots, necrotic spots on leaves, reddish-brown streaks on shoots, cracks, cankers, wilting and dieback (EFSA Panel on Plant Health 2014). Disease severity depends on cultivar susceptibility and environmental conditions (Peros et al. 1995). The bacterium enters plants through wounds or natural openings and colonizes xylem vessels, forming biofilms. Later, also phloem and cambial tissues were found to be infected. The resulting disruption of water transport causes tissue decay (EPPO 2009).

Although *X. ampelinus* is relatively rare compared to other grapevine pathogens, its impact in the affected regions can be serious. It is listed as a quarantine pathogen in several countries due to its potential to spread and damage viticulture industries (EPPO 2025). Understanding and monitoring *X. ampelinus* is critical to maintaining healthy vineyards and ensuring the sustainability of grape production, particularly in Europe where it has been of most concern in the past.

Many plant-associated bacteria of the phylum Pseudomonadota, including Alphaproteobacteria (e.g., *Bradyrhizobium*, *Mesorhizobium*, *Sinorhizobium*), Betaproteobacteria (e.g., *Acidovorax*, *Burkholderia*, *Paracidovorax*, *Papaburkholderia*, *Ralstonia*) and Gammaproteobacteria (e.g., *Pantoea*, *Pseudomonas*, *Xanthomonas*) rely on a specialized protein secretion system for their interaction with the plant, be it a beneficial or a pathogenic relationship (Büttner 2016; Schreiber et al. 2021; Teulet et al. 2022). The type III secretion system (T3SS) injects effector proteins, called type III effectors (T3Es), into the host cells to the benefit of the bacteria, thereby favoring the colonization (infection or symbiotic) process. Historically, the T3SS has first been characterized in human pathogens, focusing on species of *Yersinia*, *Salmonella* and *Shigella* (Cornelis 2002).

Based on sequence similarities, T3SSs have been classified into seven major classes, reflecting the bacteria from which they were first characterized (Chlamy, Hrp1/Hrc1, Hrp2/Hrc2, SPI-1 [Inv/Mxi/Spa], SPI-2 [Esc/Ssa], Rhizo [Rhc], Ysc) (Cornelis 2006; Abby and Rocha 2012; Zboralski et al. 2022). The T3SS is a multi-component protein complex which contains 13 conserved proteins in different copies, namely the ATPase SctN, the sorting platform consisting of SctK, SctL and SctQ, the inner membrane secretion apparatus consisting of SctR, SctS, SctT, SctU and SctV, the IM ring consisting of SctD and SctJ, the inner rod protein SctI, the outer membrane secretin SctC, and the needle filament SctF (Abby and Rocha 2012), according to the unified nomenclature (Wagner and Diepold 2020). Nine of them (SctJLNQRSTUV) have clear homologs in the flagellar basal body, and two (SctP and SctF) have functional and structural counterpart analogs in the flagellum. Compelling evidence suggests that the non-flagellar T3SS evolved from the flagellum (Abby and Rocha 2012; Diepold and Armitage 2015; Worrall et al. 2023). The two gate proteins SctU (and its flagellar homolog FlhB) and SctV (and its flagellar homolog FlhA) constitute the most conserved components of the T3SS. As such, they are considered as a signature for T3SS presence (or remnants thereof) and allow classification of the non-flagellar T3SS into seven classes (Gazi et al. 2012; Zboralski et al. 2022).

Different T3SS classes exhibit distinct regulatory mechanisms governing gene expression (Büttner 2012). For instance, the Hrp1/Hrc1 system is controlled by the extracytoplasmic function (ECF) sigma factor HrpL, which is thought to direct the RNA polymerase to the ‘*hrp* box’ in the promoters of T3SS genes (Kvitko and Collmer 2023). The *hrpL* gene itself is regulated by a heterohexamer of HrpR and HrpS in strains of *Pseudomomas syringae* or by a HrpS homohexamer in the Enterobacterales (Kvitko and Collmer 2023). The other plant-associated T3SS, Hrp2/Hrc2, is under the control of the two-component response regulator HrpG. It is thought that HrpG is post-translationally activated by protein phosphorylation by a hitherto unknown component, leading to transcriptional activation of the *hrpX* gene (Wengelnik et al. 1996). The AraC-type regulator HrpX (in *Acidovorax* and *Xanthomonas*)/HrpB (in *Burkholderia* and *Ralstonia*) then binds to a conserved promoter element, the *hrpII*/PIP box, thus regulating T3SS and T3E genes (Koebnik et al. 2006). Activation of HrpX-controlled genes strictly relies on the presence of a PIP box and a properly spaced –10 promoter motif (Cunnac et al. 2004a; Tsuge et al. 2005; Furutani et al. 2006; Lipscomb and Schell 2011). Many, if not most HrpX/HrpB-regulated genes encode components of the T3SS machinery, T3Es, or secreted hydrolases, such as endoglucanases, proteases, esterases, and pectinases (Noël et al. 2001; Cunnac et al. 2004b; Occhialini et al. 2005; Tamura et al. 2005; Lipscomb and Schell 2011; Jiménez-Guerrero et al. 2020; Teper et al. 2021). The presence of HrpX-regulated promoters, identified either experimentally or in silico, is used to predict T3E genes, which can then be confirmed and characterized in more detail (Noël et al. 2001; Furutani et al. 2006; Jiménez-Guerrero et al. 2020).

In order to better understand the host-pathogen molecular interactions within the genus *Xylophilus*, we aimed at predicting and comparing T3SSs and candidate T3E proteins at a genome-wide level. Our work provides new insights into the evolution of T3SSs in the genus *Xylophilus* and the presence of T3Es in these species. For three candidate T3Es, we provide evidence that they are bona fide effectors likely translocated by the T3SS and injected into grapevine plants.

## MATERIAL AND METHODS

### Biological material

#### Bacterial strains and growth conditions

Strains of *Escherichia coli* (DH10B) and *Xanthomonas campestris* (8000 and its mutants in the T3E *avrBs1* and the type III secretion gene *hrcV*) were used in this study (Durfee et al. 2008; Xu et al. 2008; Zhao et al. 2013). The *X. campestris* strains were cultivated at 28 °C on solid PSA (Peptone-Sucrose-Agar) medium (peptone 10 g/L, sucrose 10 g/L, agar 16 g/L, glutamic acid 1 g/L).

*Escherichia coli* cells were grown at 37 °C on LB (lysogenic broth) agar plates (LB Lennox agar 35 g/L; Euromedex, Souffelweyersheim, France) or in liquid LB medium (tryptone 10 g/L, yeast extract 5 g/L, NaCl 10 g/L). Antibiotics were added, when needed, to the media at the following final concentrations: 100 mg/L for rifampicin and 5 mg/L for tetracycline.

#### Plasmids

The plasmid pBBR1MCS-3::avrBs1_59-445_ is a derivative of the plasmid pBBR1MCS-3. It encodes an N-terminally truncated variant of AvrBs1 from *X. campestris*. This variant lacks its own type III secretion signal and is used to construct translational reporter gene fusions with presumed type III secretion signals from candidate T3E genes (Kovach et al. 1995; Zhao et al. 2013). Reporter fusions were constructed by GenScript (Piscataway Township, NJ, USA) from synthetic DNA that consisted of the 100-base-pair noncoding sequence upstream of the translation initiation codon, as well as the 300-base-pair coding sequence that would include the N-terminal type III secretion signal from the candidate genes. The helper plasmid pRK2013 was used to transfer plasmids via conjugation from *E. coli* to *Xanthomonas* by triparental mating (Figurski and Helinski 1979).

#### Plant material

The pepper cultivar ECW-10R, which triggers a hypersensitive response upon perception of AvrBs1 by the corresponding *Bs1* resistance gene product, was used for plant inoculation experiments (Kousik and Ritchie 1999). Plants were grown in a greenhouse with natural daylight cycles at 27°C ± 5°C and 60% relative humidity.

### Bioinformatic analyses

#### Genomic resources

Genomic resources, such as genome sequences, genome annotations, predicted open reading frames (ORFs) and proteomes, were retrieved from the NCBI Datasets resource (https://www.ncbi.nlm.nih.gov/datasets/) (Sayers et al. 2025). Quality and correct taxonomic classifications were evaluated using the Type (Strain) Genome Server (TYGS; https://tygs.dsmz.de/) and the Microbial Genomes Atlas (MiGA; https://gateway.microbial-genomes.org/) (Rodriguez-R et al. 2018; Freese et al. 2026). Genome-wide average nucleotide identities (ANI) were calculated using FastANI version 1.3 (Jain et al. 2018), as implemented at the French Galaxy server (https://usegalaxy.fr/) (Galaxy Community 2024).

#### Sequence comparisons and phylogenetic trees

Sequence similarity searches and pairwise sequence comparisons were performed at the National Center for Biotechnology Information (NCBI) using the BLAST and Needleman-Wunsch algorithm, respectively (Needleman and Wunsch 1970; Altschul et al. 1990; Sayers et al. 2025). Multiple sequence alignments were generated with MUSCLE (Edgar 2004), as implemented at the EMBL’s European Bioinformatics Institute (EMBL-EBI) (Thakur et al. 2025). To compare T3SS gene clusters, the corresponding regions were extracted from the annotated GenBank files using the Artemis genome browser (Rutherford et al. 2000). Clinker was used for the comparison and visualization of genomic regions (Gilchrist and Chooi 2021).

Phylogenetic trees based on sequence comparisons were generated at Phylogeny.fr (https://www.phylogeny.fr/), with default parameters (Dereeper et al. 2008). Core genome-wide comparisons were established using the M1CR0B1AL1Z3R suite (https://microbializer.tau.ac.il/) (Shimony et al. 2025). For better visualization of the trees, we used the Interactive Tree of Life (iTOL) tool (Letunic and Bork 2024).

#### Prediction of protein secretion systems and type III effectors

Protein secretion systems were predicted using TXSScan, as implemented in the Galaxy platform at the Institut Pasteur (https://galaxy.pasteur.fr/root?tool_id=toolshed.pasteur.fr/repos/odoppelt/txsscan/TXSScan/1.0.2) (Abby et al. 2024). The type IV and type VI secretion system were also predicted using SeCreT4 and SeCreT6 (https://bioinfo-mml.sjtu.edu.cn/SecReT4/; https://bioinfo-mml.sjtu.edu.cn/SecReT6/), respectively (Bi et al. 2013; Zhang et al. 2023).

To predict T3Es, we employed Effectidor II, a pan-genomic AI-based algorithm for the prediction of T3Es (https://effectidor.tau.ac.il) (Wagner et al. 2025). The input consisted of predicted ORFs and GFF3 annotations, as available from NCBI, targeting the genomes of five strains that harbor a Hrp2/Hrc2-related T3SS (assembly accession numbers GCA_003217575.1, GCA_024832295.1, GCA_033546935.1, GCA_009906855.1, and GCA_026951875.1 for strains CECT7646, CFBP 1192, GOD-11R, KACC 21265, and Kf1, respectively). The predicted proteomes of strains ASV27, bin23, GW821-FHT01B05, Leaf220 and SP210_2 were used as a negative training set (assembly accession numbers GCF_016428875.1, GCA_036946225.1, GCA_038961845.1, GCA_001421705.1 and GCF_913776965.1, respectively) to represent strains that do not encode a Hrp2/Hrc2-related T3SS. Search options included prediction of the type III secretion signal and searching for the presence of a PIP-box motif in the upstream region of all ORFs. All other parameters were left at their default settings.

#### Other bioinformatic tools (Prediction of protein domains and structural modeling of predicted type III effectors)

Protein domains and sequence motifs were predicted using InterProScan (https://www.ebi.ac.uk/interpro/) (Jones et al. 2014). Three-dimensional structures of proteins were predicted by AlphaFold3 (https://alphafoldserver.com/welcome) (Abramson et al. 2024). FoldSeek was used to identify structural homologs (https://search.foldseek.com/) (van Kempen et al. 2024). To reduce complexity of datasets, groups of homologous sequences were clustered using CD-Hit at usegalaxy.fr (Galaxy Version 4.8.1) (Li et al. 2001). Signal sequences for secretion via the Sec or Tat pathway were predicted with SignalP 6.0 (https://services.healthtech.dtu.dk/services/SignalP-6.0/) (Teufel et al. 2022). GeNomad in its Galaxy version 1.11.1 was used to identify mobile genetic elements, such as prophages (Camargo et al. 2024).

### Plant inoculation experiments

Eight to twelve weeks old plants of the pepper cultivar ECW-10R were used for plant inoculation experiments (Kousik and Ritchie 1999). Bacteria were grown on PSA medium, resuspended in 10 mM MgCl_2_ and diluted to an optical density at 600 nm wavelength of 0.1. A needleless 1-mL plastic syringe was used to inoculate the bacterial solution from the lower side of the pepper leaf into the apoplast. Each bacterial solution was inoculated on at least two different leaves from at least two different plants. Experiments were performed at least two times. Plant reactions were scored over a period of seven days after the inoculation.

## RESULTS

### Clarification of taxonomic status of publicly available genome sequences

In order to obtain an overview of the presence of virulence-associated secretion systems in the genus *Xylophilus*, we first extracted all genome sequences that are assigned to this bacterial genus from NCBI (https://www.ncbi.nlm.nih.gov/datasets/genome/). NCBI GenBank lists 19 genome sequences for this genus (accessed on 29-JUL-2025), six for the species *X. ampelinus*, one for the species *X. rhododendri*, and twelve without species assignment (Table 1).

**Table 1.**
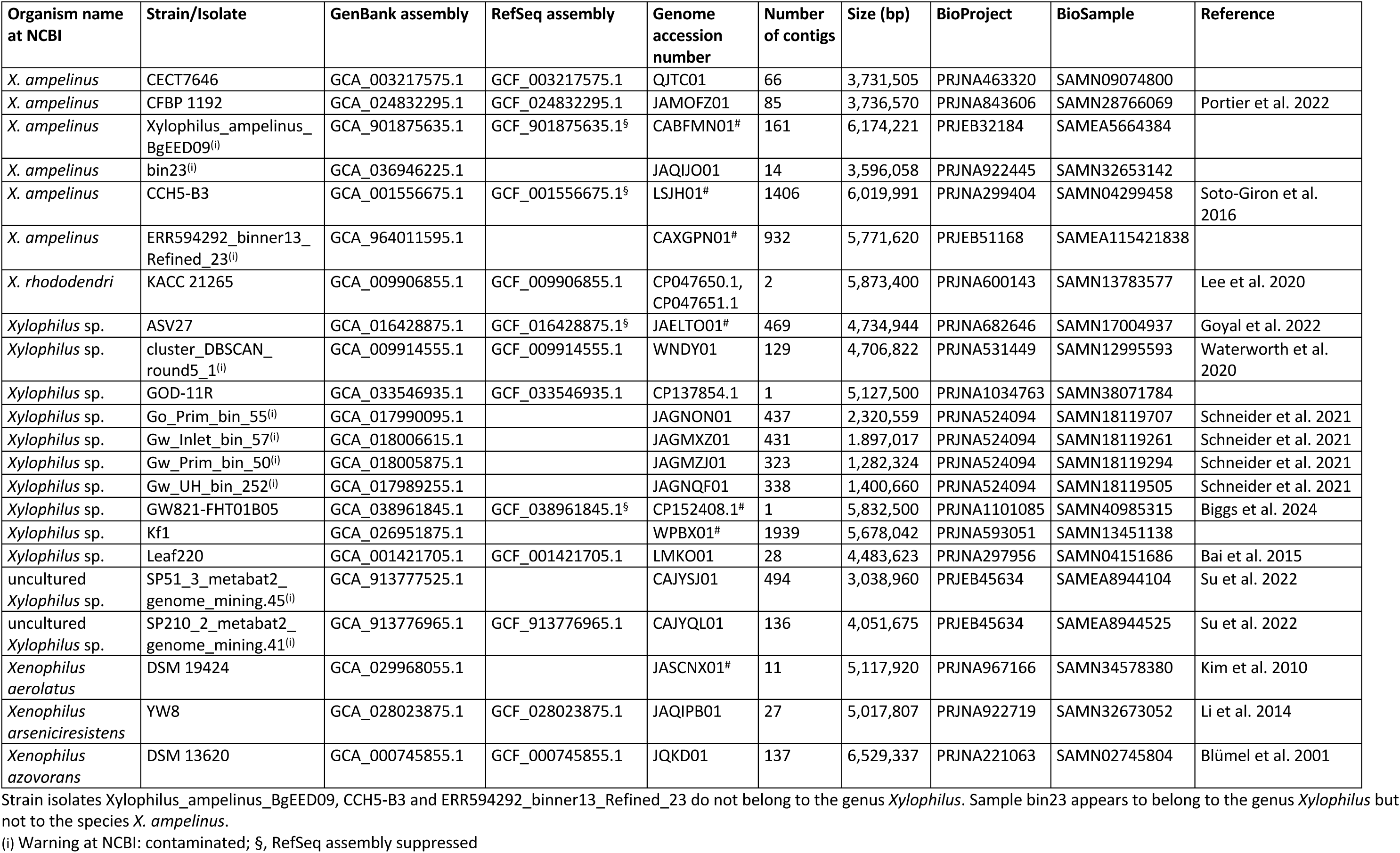
*Xylophilus* genome sequences at NCBI GenBank.

The Type strain Genome Server (TYGS; https://tygs.dsmz.de) was applied to confirm the taxonomic classification (Freese et al. 2026) (accessed on 30-JUL-2025) (Supplementary Table S1a). Only the two sequences of strains CECT7646 and CFBP 1192, both corresponding to the same type strain, which was isolated from a necrotic grapevine plant in Crete, Greece, in 1966, were reported to belong to the species *X. ampelinus*. Strain KACC 21265 was confirmed to belong to the species *X. rhododendri*. Eight genome sequences (strains/isolates ASV27, bin23, Cluster_DBSCAN, GOD-11R, GW821-FHT01B05, Leaf220, SP51_3, SP210_2) were classified as a potential new species of the genus *Xylophilus*. Three of the genome sequences (strains/isolates BgEED, CCH5-B3, ERR594292) were classified as a potential new species of the genus *Xenophilus*. Four isolates with surprisingly small genome sizes, originating from metagenome assemblies from wastewater treatment plants in Germany (Schneider et al. 2021), were classified as potential new species of the genera *Acidovorax* (Go_Prim_bin_55, Gw_Prim_bin_50, Gw_UH_bin_252) or *Macromonas* (Gw_Inlet_bin_57). Lastly, the genome sequence of strain Kf1 was flagged as potentially unreliable identification result due to potential sequence contamination, belonging to the species *Staphylococcus epidermidis*.

The TypeMat genome-assisted taxonomic classifier at the Microbial Genomes Atlas (MiGA; https://uibk.microbial-genomes.org/) was used to complement this analysis (accessed on 30-JUL-2025) (Supplementary Table S1b). Except for isolate Cluster_DBSCAN, which was classified as a new species of the genus *Paracidovorax*, all classifications with respect to *Xylophilus* and *Xenophilus* were confirmed. The four wastewater metagenome assemblies were flagged as intermediate-quality assemblies (in contrast to all the other sequences, which were of high or excellent quality) and assigned to the genera *Acidovorax* (Go_Prim_bin_55, Gw_Prim_bin_50) and *Variovorax* (Gw_UH_bin_252). Surprisingly, strain Kf1 was reported as a new species of the genus *Xylophilus*, which might be due to the fact that MiGA only considers contigs with at least 1,000 bp for classification. In the case of strain Kf1, only 116 of 1,939 contigs were considered and sequence information corresponding to 1,823 contigs (775,918 bp) was omitted, which might explain the classification as a *Staphylococcus* strain by TYGS. Hence, ten genome sequences were classified as *Xylophilus* by both tools, one was only supported by TYGS (Cluster_DBSCAN) and another was only supported by MiGA (Kf1).

To clarify the sequence relationships among the various genomes, genome-wide ANI were calculated (Jain et al. 2018) (Supplementary Table S2a). Four genome sequences, originating from the wastewater metagenome assemblies, were omitted due to poor quality and apparent incompleteness as evidenced by their small sizes and quality metrics (Supplementary Table S1b). This analysis suggests that the twelve *Xylophilus* strains, including Cluster_DBSCAN and Kf1, belong to ten different species. CFBP 1192 and CECT7646, with genome sequences sharing more than 99.99% sequence identity, are both equivalent of the *X. ampelinus* type strain of the species held in two different collections. The genome sequences of isolates SP51_3 and SP210_2 were more than 97.8% identical, suggesting that they both correspond to the same *Xylophilus* species. The three sequences that appeared to belong to the genus *Xenophilus* (strains/isolates BgEED, CCH5-B3, ERR594292) were more than 98.5% identical to each other, indicating that they belong to the same species. The most closely related of these three strains is *Xenophilus aerolatus*, whose type strain, DSM 19424, shares more than 91.9% sequence identity with their genomes.

Digital DNA-DNA hybridization (DDH) was performed to complement the ANI calculations (Meier-Kolthoff et al. 2013) (Supplementary Table S2b). This analysis confirmed the conclusion that the twelve *Xylophilus* genomes correspond to 10 species. DDH of the genome sequences of isolates SP51_3 and SP210_2 was estimated with 86%, significantly above the generally accepted species boundary of 70% (Chun et al. 2018). Likewise, strains/isolates BgEED, CCH5-B3, ERR594292 had pairwise DDH values of at least 96% to each other and were most similar to the *X. aerolatus* type strain DSM 19424 with 45-46% DDH values. Taken together, these analyses established a well-curated dataset of twelve genome sequences from eleven different strains for further analyses.

### Phylogenetic tree and presence of flagellar and type III secretion genes

Since many Gram-negative plant-associated bacteria rely on a T3SS for pathogenicity or symbiosis, and because most of the *Xylophilus* strains were found to be associated with plants, we hypothesized that these bacteria encode such a secretion system in their genomes. The genome sequences were scrutinized for the presence of T3SS-related sequences based on two of the most conserved proteins, SctV (a homolog of the flagellar FlhA) and SctU (a homolog of the flagellar FlhB) (Gazi et al. 2012; Tampakaki 2014; Zboralski et al. 2022). One representative sequence from each of the seven T3SS classes was used as the query, as well as one flagellar homologous sequence, i.e., seven SctU and FlhB sequences, and seven SctV and FlhA sequences (Figure 1). BLASTP searches against the predicted proteomes of eleven *Xylophilus* strains identified 19 hits for SctU/FlhB and 19 hits for SctV/FlhA. Each strain contained one to three paralogs of each protein. Of the 19 hits, four represent paralogous doublets derived from the two *X. ampelinus* type strain genomes. Since the genome sequence of isolate SP51_3 was not annotated, we used TBLASTN to find homologs of SctU and SctV.

**Figure 1.**
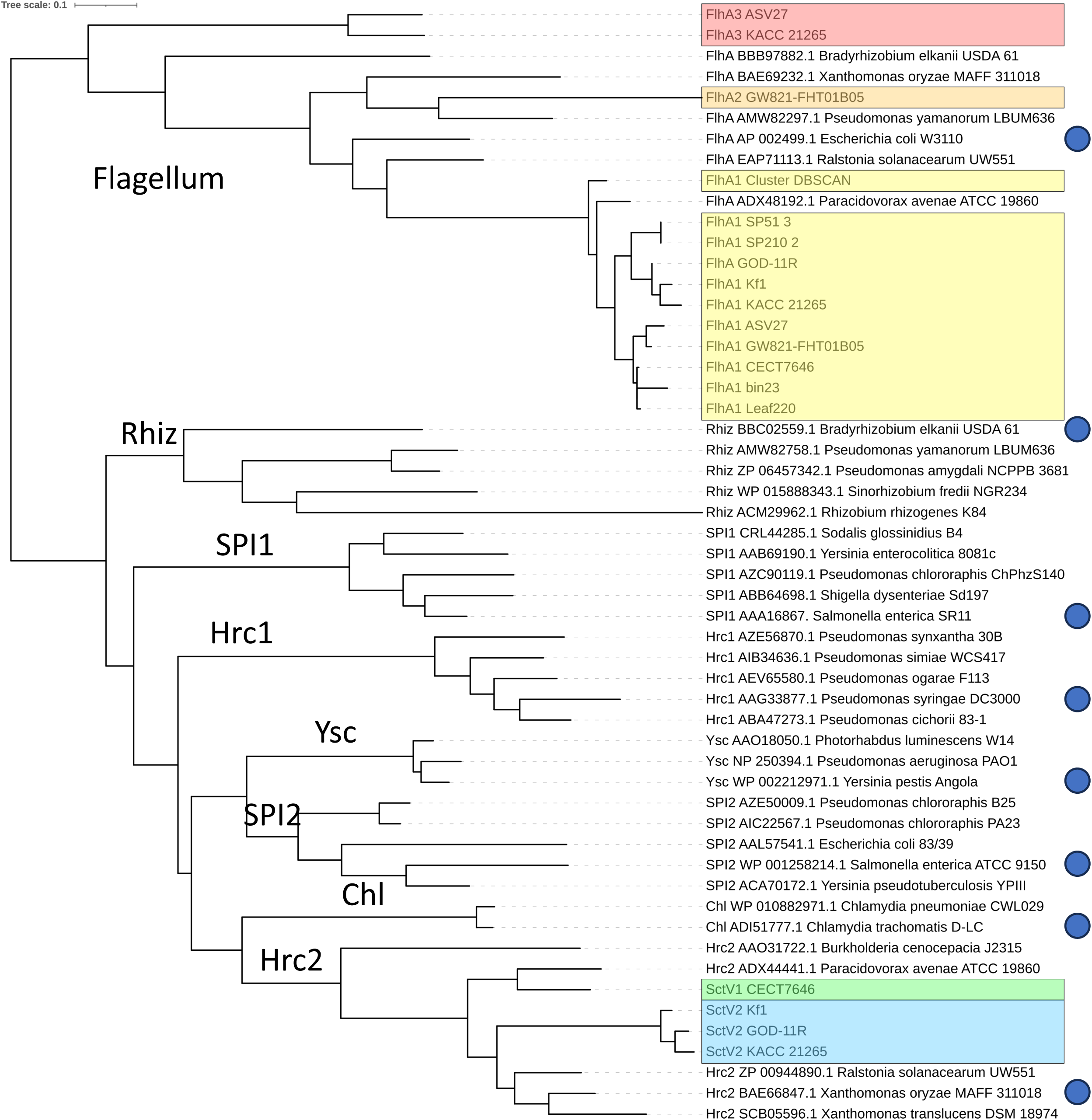
Phylogenetic tree of representative SctV/FlhA proteins and homologs from *Xylophilus*. *Xylophilus* sequences are boxed with the three flagellar proteins in yellow (type 1), orange (type 2) and red (type 3), and the Sct proteins in green (T3SS#1) and blue (T3SS#2). Strain CFBP 1192 was omitted because it is a clone of the *X. ampelinus* type strain CECT7646. Sequences that were used to identify SctV/FlhA homologs in proteomes of *Xylophilus* are indicated by a blue dot.

A multiple sequence alignment of the 20 *Xylophilus* SctV homologs revealed that they fall into five groups (Supplementary Data S1). A phylogenetic tree reconstructed based on the multiple sequence alignment of the 20 sequences as well as representatives of flagellar and T3SS sequences (Zboralski et al. 2022), further supported that these sequences form five clades, which we tentatively called SctV1, SctV2, FlhA1, FlhA2 and FlhA3 (Figure 1). Members of both SctV clades aligned within the Hrp2/Hrc2 class of SctV proteins. Members of clade FlhA1 and the FlhA2 sequence clustered within the flagellar class. Finally, the two FlhA3 sequences formed a sister clade of the flagellar class, suggesting that they belong to flagellar systems. A similar result was obtained when comparing representative SctU/FlhB sequences with the homologs from *Xylophilus* (Supplementary Figure S1).

To examine the distribution of the various flagellar and secretion systems among the *Xylophilus* strains, a core-genome based phylogenetic tree was computed using the M1CR0B1AL1Z3R suite (Shimony et al. 2025) (Figure 2). All strains were found to possess the type-1 flagellum comprising FlhA1 and FlhB1. The second flagellar type with FlhA2 and FlhB2 was only found in strain GW821-FHT01B05. The distribution of the third flagellar type with FlhA3 and FlhB3 was patchy and did not follow phylogenetic descent. The Hrp2/Hrc2 T3SS with SctU1 and SctV1 was only found in the *X. ampelinus* strain (CECT7646/CFPB 1192). The second Hrp2/Hrc2 T3SS type was present in a lineage comprising the strains Kf1, GOD-11R, and KACC 21265.

**Figure 2.**
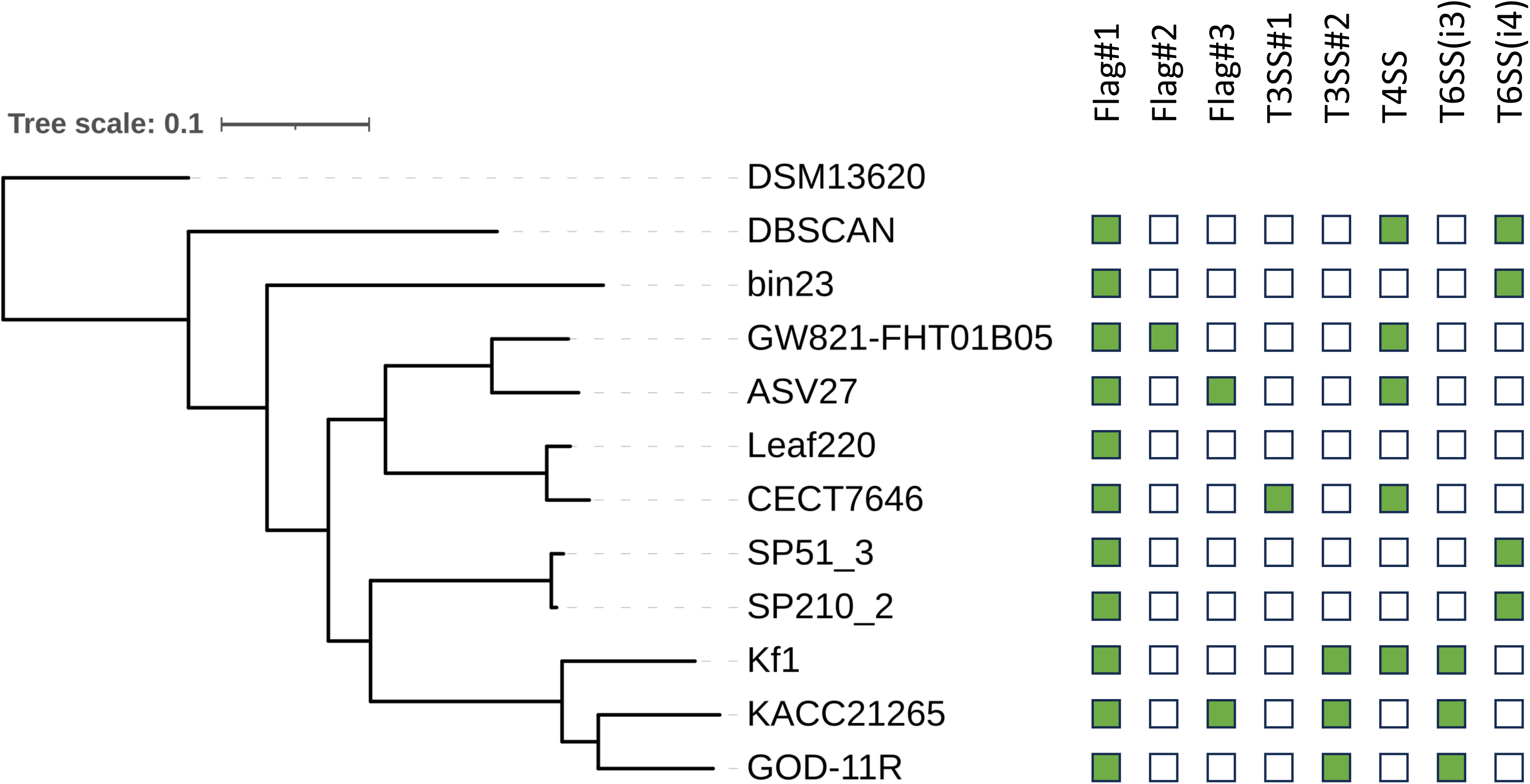
Presence of flagellar and secretion systems in strains of *Xylophilus*. The phylogenetic tree was calculated from the core proteome using M1CR0B1AL1Z3R, and is based on 644 proteins with 231,120 aligned amino acids. All nodes are supported with 100% bootstrap values. The genome of the *Xenophilus* type strain DSM 13620 was used to root the tree. Presence of flagellar and secretion systems, as deduced from homology to the conserved *sctV*/*flhA* gene and supported by TXSScan, SeCreT4 and SeCreT6 analyses, is indicated by green squares and absence by open squares.

We also evaluated the presence of two additional multi-protein secretion systems, the type 4 and type 6 secretion systems (T4SS, T6SS). The presence of these systems was predicted using SeCreT4 and SeCreT6 (Bi et al. 2013; Zhang et al. 2023). The i3-class T6SS was found in the same clade as the type-2 T3SS, whereas the i4-class T6SS was found in three genetic lineages comprising DBSCAN, bin23, and the two similar isolates SP51_3 and SP210_2. Gene clusters encoding T4SS were found in five strains, DBSCAN, Kf1, GW821-FHT01B05, ASV27 and CECT7646/CFPB 1192 and their presence-absence pattern suggests multiple gain and loss events of this system along the phylogenetic tree (Fig. 2). These results were confirmed using the TXSScan program (Supplementary Table S3) (Abby et al. 2024).

Hrp2/Hrc2-class T3SSs are under transcriptional control of HrpX, a AraC-type regulator that binds to a conserved plant-inducible promoter (PIP) box. When a matching -10 promoter element is in a proper distance of 30 to 32 base pairs, genes are transcribed, most of them encoding components of the T3SS, T3Es and type II-secreted cell wall-degrading enzymes. Indeed, perfect PIP patterns with the consensus sequence TTCGB-N_15_-TTCGB-N_30-32_-YANNNT were found in the promoters of the Hrp2/Hrc2 T3SS gene clusters, in front of *sctD* and *sctU* in strain CECT7646 and in front of *sctQ* in strain KACC 21265 (Table 2). Nine of the 16 PIP patterns in the genome of strain CECT7646 were found upstream of T3E genes (*hopQ*, *ripC*, *ripE*, *ripAC*, *ripAQ*, *xopD*, *xopZ*, *xopAI, xopBG*) and one upstream of a pectate lyase gene (*ripW*). Two genes, DFQ15_RS08770 and DFQ15_RS08785, had only a homolog in *Sphingomonas* sp. NCPPB 2930 in a region with a similar T3SS.

**Table 2.**
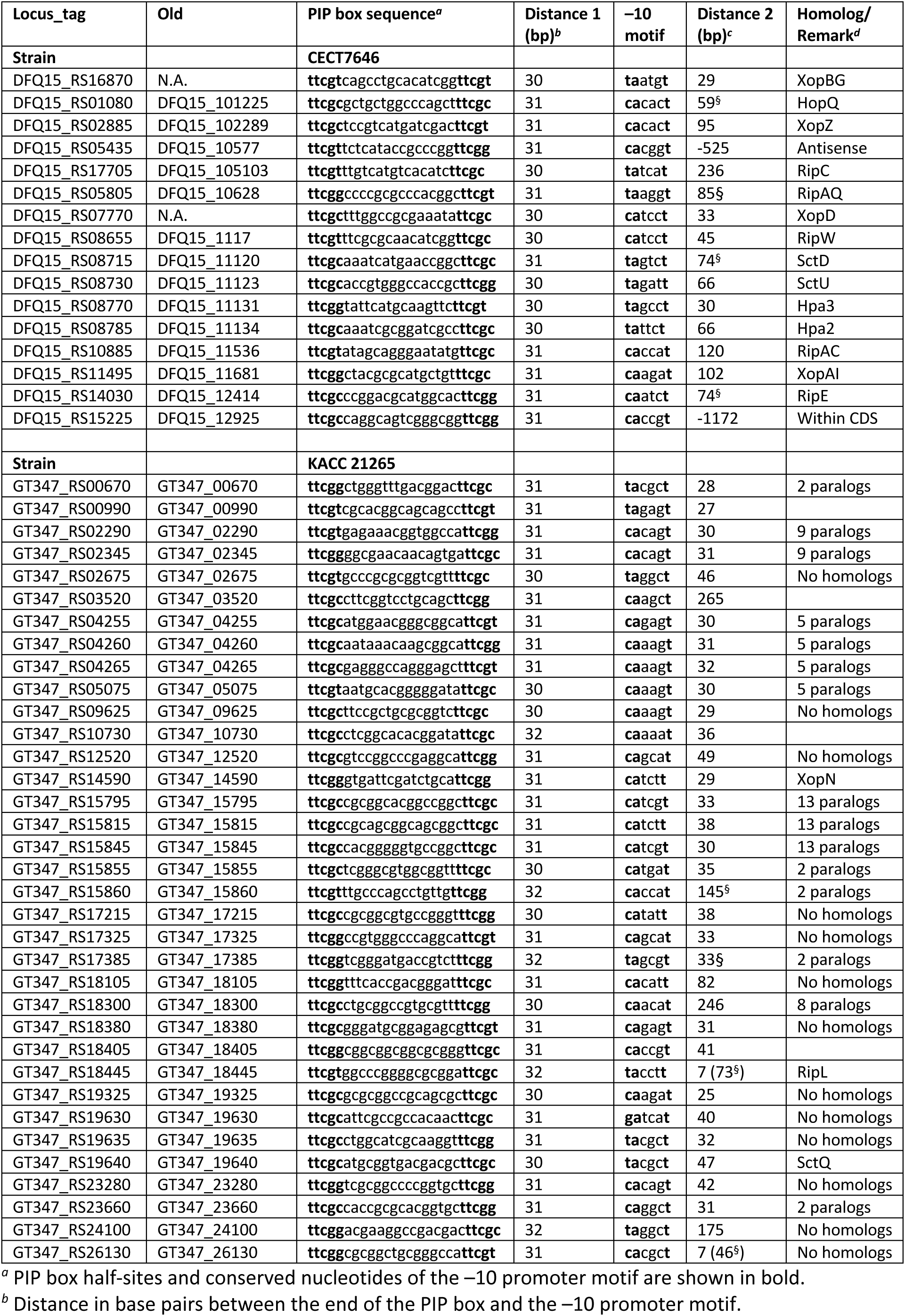
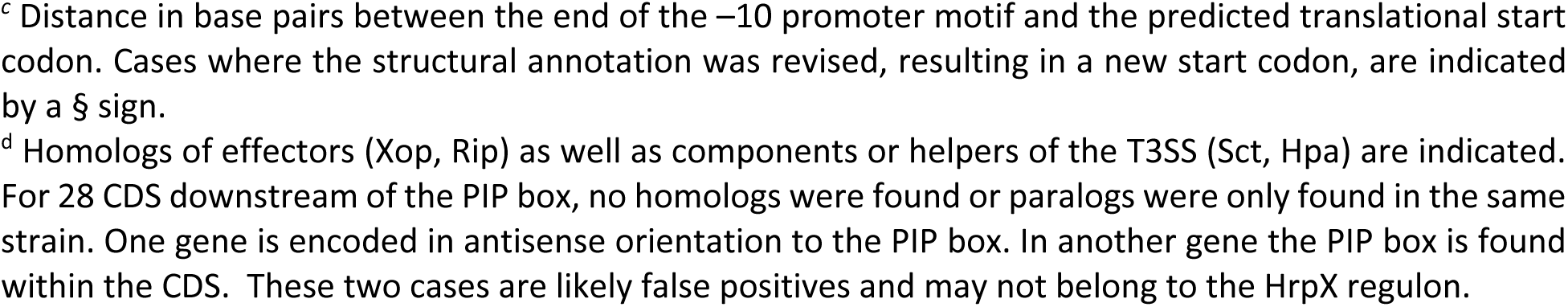
Candidate HrpX-regulated promoters of *X. ampelinus* strain CECT7646 and *X. rhododendri* strain KACC 21265.

The predicted PIP-box harboring gene repertoire in strain KACC 21265 was different from that of strain CECT7646. Among its 32 genes encoding hypothetical proteins were two candidate T3E genes. Protein QHI99106.1 (locus tag GT347_14590) was 22.90% identical to XopN (WP_011348010.1) over 69% of its sequence, and protein QHI99786.1 (locus tag GT347_18445) was 26.04% identical to RipL (WP_014630780.1) over 80% its sequence. Notably, 15 genes were found to encode members of strain-specific protein families with unknown function, with up to 13 paralogs (Supplementary Table S4; Supplementary Data S2). All of them are annotated as hypothetical proteins. We used InterProScan to shed light on their potential functions. However, no known protein domain was found for any of them. A signal sequence for secretion was predicted for two members of the 13 homologs of protein family 4. For 13 additional hypothetical proteins beyond the eight protein families, we did not find any homolog using BLASTP searches at NCBI GenBank (Table 2).

### Comparative genomic organization of flagellar and type III secretion gene clusters within *Xylophilus* strains

To better classify the flagellar systems and the T3SSs, we compared their genomic organization. To this end, we manually curated the structural and functional annotation of the genomic regions where the *sctV* and *sctU* genes were found, using BLAST and InterProScan searches.

Strain GW821-FHT01B05 appears to encode two types of flagella in large gene clusters. The gene cluster with *flhA1*, which was found in all *Xylophilus* strains, spans 55 genes in strain GW821-FHT01B05, among them two flagellin genes, three *flh* genes (ABF), 13 *flg* genes (ABCDEFGHIJKLM), 18 *fli* genes (ADEFGHIJKLMNOPQRST), two *mot* genes (AB) and seven chemotaxis genes (ABDRWYZ) (Supplementary Figure S2). The second gene cluster with *flhA2* contains 38 genes with only one flagellin gene, two *flh* genes (AB), 15 *flg* genes (ABCDEFGHIJ_2_KLMN), 18 *fli* genes (ADEFGHIJK_2_LMNPQRST), and two *mot* genes (AB). This strain contains six additional chemotaxis genes (ABDRWY), which are however encoded in another genomic location and it remains to be determined whether they correspond to the FlhA2 flagellar system.

Another candidate flagellar system, which includes *flhA3*, was found in strains ASV27 and KACC 21265. This system lacks several of the *flg*, *flh* and *fli* genes. For this system, no flagellin could be identified. Candidate genes for nine Flg proteins (ABCDEFGHI), two Flh proteins (AB) and twelve Fli proteins (AEFHIJMNOPQR) were predicted. Some of the missing genes, such as *fliG*, *fliK* and *fliL*, might correspond to the hypothetical proteins that are encoded in the gene cluster (Supplementary Figure S2).

Bona fide T3SSs were found in five strains (Figure 3). For comparison, we included the T3SSs of the Hrp2/Hrc2-representative strains that were used for classification of the SctV/FlhA and SctU/FlhB proteins (Figure 1; Supplementary Figure S1). The two *X. ampelinus* genomes were found to encode a T3SS with 13 *sct* genes (CDJKLNPQRSTUV), which have a genomic organization that is similar to the ones found in clade 1-xanthomonads (*Xanthomonas hyacinthi*, *Xanthomonas theicola*, *Xanthomonas translucens*), in the *Ralstonia solanacearum* species complex, but also in *Collimonas fungivorans*, *Paraburkholderia andropogonis* and *Uliginosibacterium gangwonense* (Pesce et al. 2017). Notably, the *Xanthomonas hrpB* (eight genes, including *sctJ*, *sctK*, *sctL*, *sctN* and *sctT*), *hrpC* (three genes, *sctU*, *sctV* and *sctP*) and *hrpD* (four genes, including *sctQ*, *sctR* and *sctS*) operons are present (Weber et al. 2007). As in all these bacteria, the AraC-type transcriptional activator HrpX is encoded within the T3SS gene cluster and likely forms an operon with *sctC*. A second HrpX paralog with 53% sequence identity is encoded at the border of the T3SS gene cluster. The gene upstream of *hrpB1* appears to encode a protein that is distantly related to the putative translocon HrpF/NolX, as supported by PSI-BLAST analysis.

**Figure 3.**
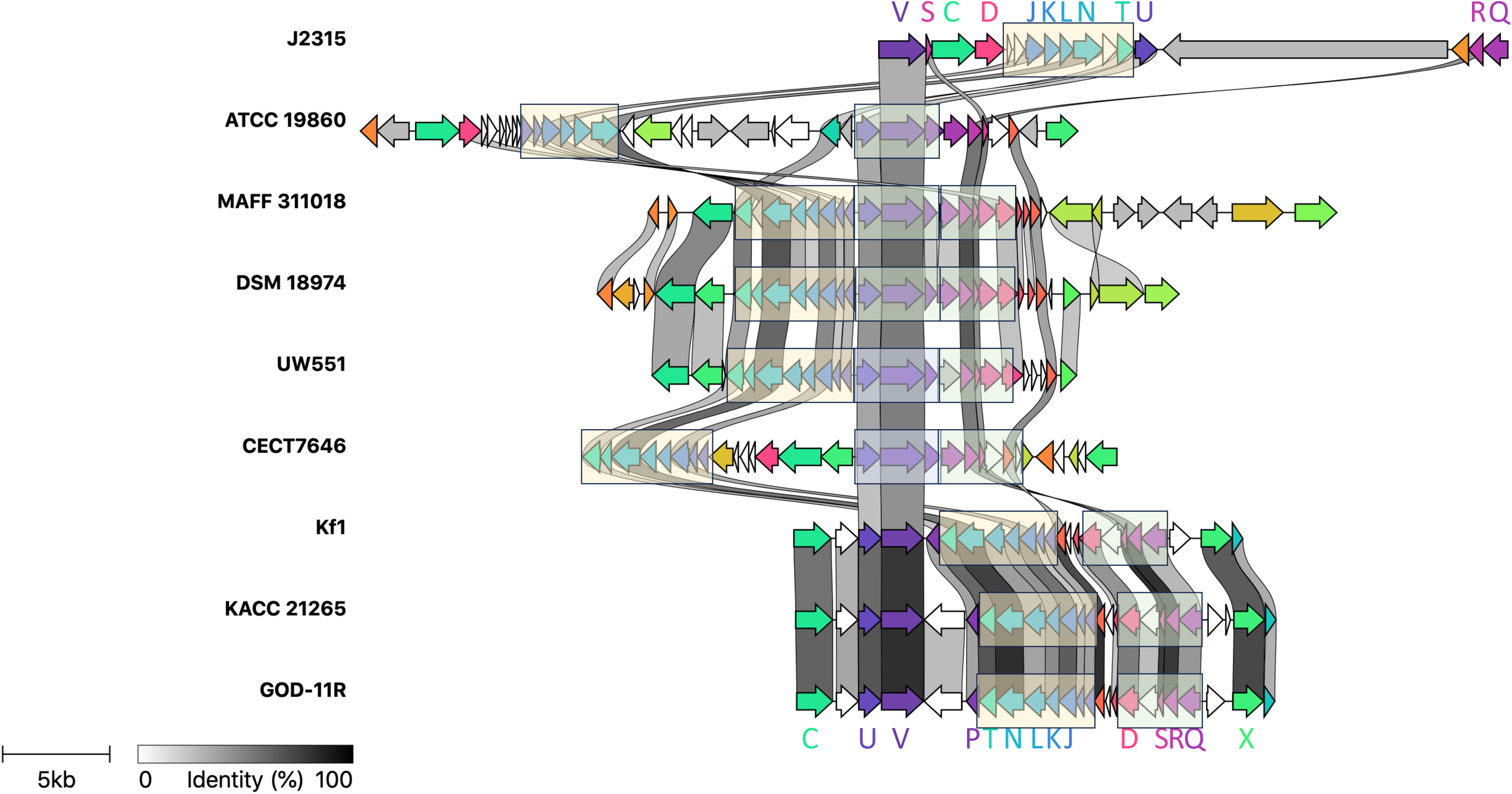
Comparison of *Xylophilus* type III secretion gene clusters. Corresponding regions, extracted from GenBank files, were compared and visualized with Clinker. Two classes of type III secretion gene clusters were observed for *Xylophilus*, as supported by synteny and sequence similarities. Conserved *sct* genes are labeled in the same color as the genes. Blocks with near-perfect synteny are boxed. The following genomic regions are depicted, from top to bottom: QU43_RS64645 (*sctV*) to QU43_RS64560 (*sctQ*) (*Burkholderia cenocepacia* J2315; NC_011001.1), Acav_0545 (*hpa2*) to ACAV_0510 (*hrpX*) (*Acidovorax avenae* ATCC 19860; CP002521.1), XOO0080 (*hpa2*) to XOO0110 (*xopAE*) (*Xanthomonas oryzae* MAFF 311018; AP008229.1), BFN94_RS13565 (*hpa2*) to BFN94_RS21845 (*xopM*) (*Xanthomonas translucens* DSM 18974; NZ_LT604072.1), RRSL_PS12385 (*sctC*) to RRSL_RS12490 (*hpaB*) (*Ralstonia solanacearum* UW551; NZ_AAKL01000026.1), DFQ15_RS08660 (*sctT*) to DFQ15_RS08795 (*hrpX2*) (*Xylophilus ampelinus* CECT7646; NZ_QJTC01000011.1), GN316_15560 (*sctC*) to GN316_15670 (*hrpB7*) (*Xylophilus* sp. Kf1; WPBX01000006.1), GT347_RS19740 (*sctC*) to GT347_RS19620 (*hrpB7*) (*Xylophilus rhododendri* KACC 21265; NP_CP047650.1), R9X41_RS04700 (*sctC*) to R9X41_RS04585 (*hrpB7*) (*Xylophilus* sp. GOD-1R; NZ_CP137854.1).

The *Xylophilus* strains KACC 21265, Kf1 and GOD11-R were found to encode another T3SS, including 13 conserved *sct* genes. Their genetic organization is different from that of systems found in other plant pathogens, such as *Acidovorax*, *Pseudomonas*, *Ralstonia* and *Xanthomonas*. For instance, *hrpB7*, which is encoded between *sctN* and *sctT* in *Ralstonia* and *Xanthomonas* and upstream of *sctT* in *Acidovorax*, is encoded downstream of *hrpX*. Also, *sctP*, which is usually encoded downstream on *sctV*, is here encoded downstream of *sctT*. No homolog of the putative translocons HrpF/NolX or HpaT was identified. And none of the *Xylophilus* T3SS gene clusters encoded an ortholog of the HrpG response regulator. In addition, we found no evidence for the presence of an HrpG ortholog elsewhere in the genome.

### Prediction of candidate type III effectors in *Xylophilus*

Candidate effectors were predicted using Effectidor II, a pan-genomic AI-based web server (Wagner et al. 2025). Prediction was targeted for the five genome sequences in which T3SSs were found, i.e., strains CECT7646, CFBP 1192, GOD11R, KACC 21265, and Kf1. Effectidor II optionally utilizes information from related strains without T3SSs (ORFs shared between strains with and without T3SSs are less likely to encode effectors). The proteomes of strains ASV27, bin23, GW821-FHT01B05, Leaf220 and SP210_2 were used for this purpose.

For *X. ampelinus* with the SctV1-type T3SS, Effectidor II predicted 36 orthology groups with scores higher than 0.25 (Table 3). Two groups, OG_3401 (NBM50_08240) and OG_1546 (DFQ15_10953), are bona fide orthologs which were separated due to differences in the assembly and structural annotation of the genome sequences. The NBM50_08240 protein variant contains a 52-aa tandem repeat, which is only present once in the DFQ15_1095 variant. The NBM50_08240 protein variant has an N-terminal extension of 378 amino acids compared to the DFQ15_1095 variant. Three orthology groups, OG_3358 (NBM50_04355), OG_3350 (NBM50_03010) and OG_3346 (NBM50_02550), were only structurally annotated in the genome sequence of strain CFBP 1192. Among the 35 bona fide orthology groups, 30 had homologs in a T3E database of plant-pathogenic bacteria, assembled from three individual databases for *Pseudomonas* (Hop’s), *Ralstonia* (Rip’s) and *Xanthomonas* (Xop’s) (Lindeberg et al. 2005; Peeters et al. 2013; Costa et al. 2024) (Table 3). Altogether, the 31 effectors fall into 28 classes, with AvrB, RipTPS and XopD being represented twice.

**Table 3.**
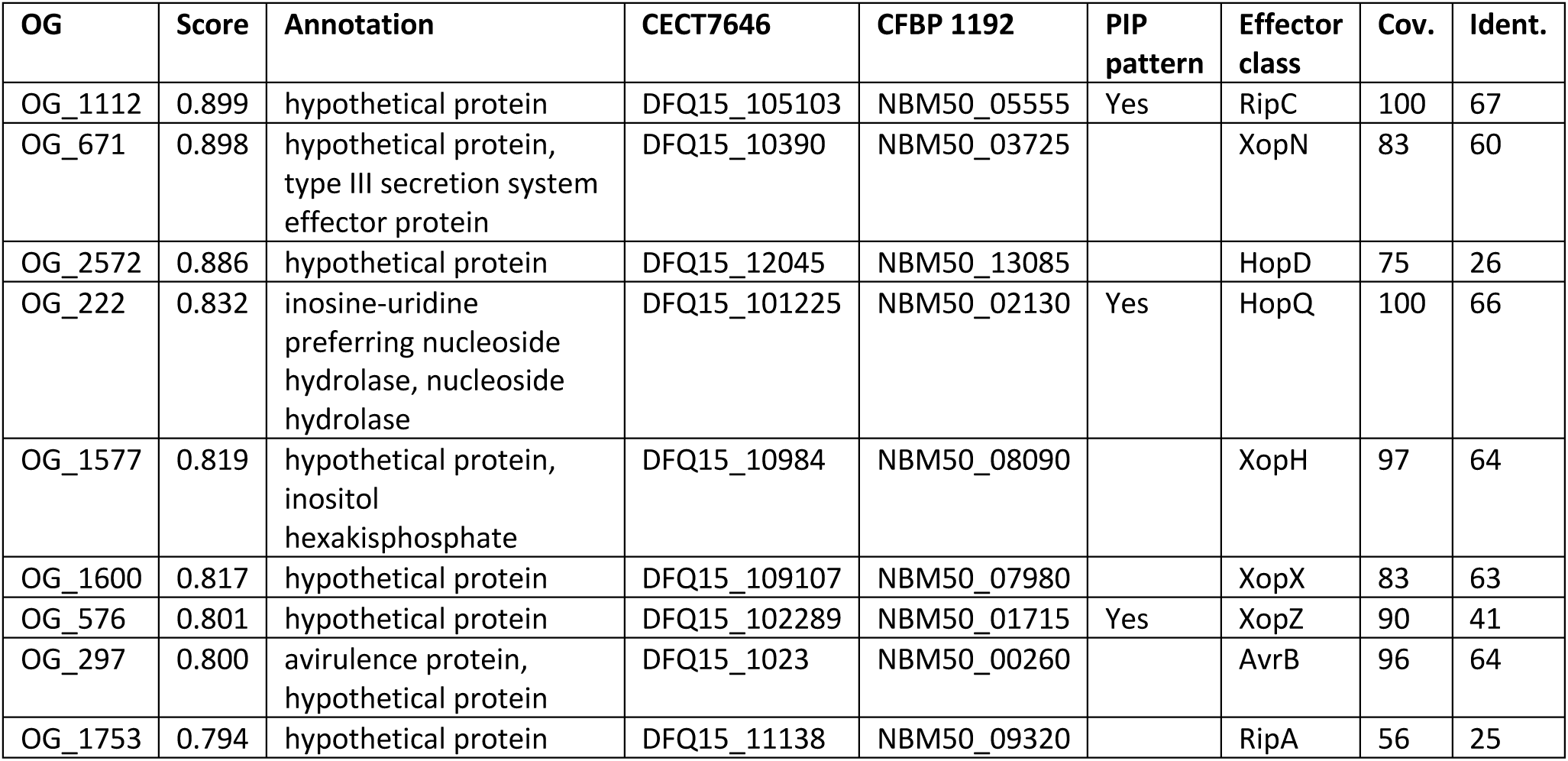
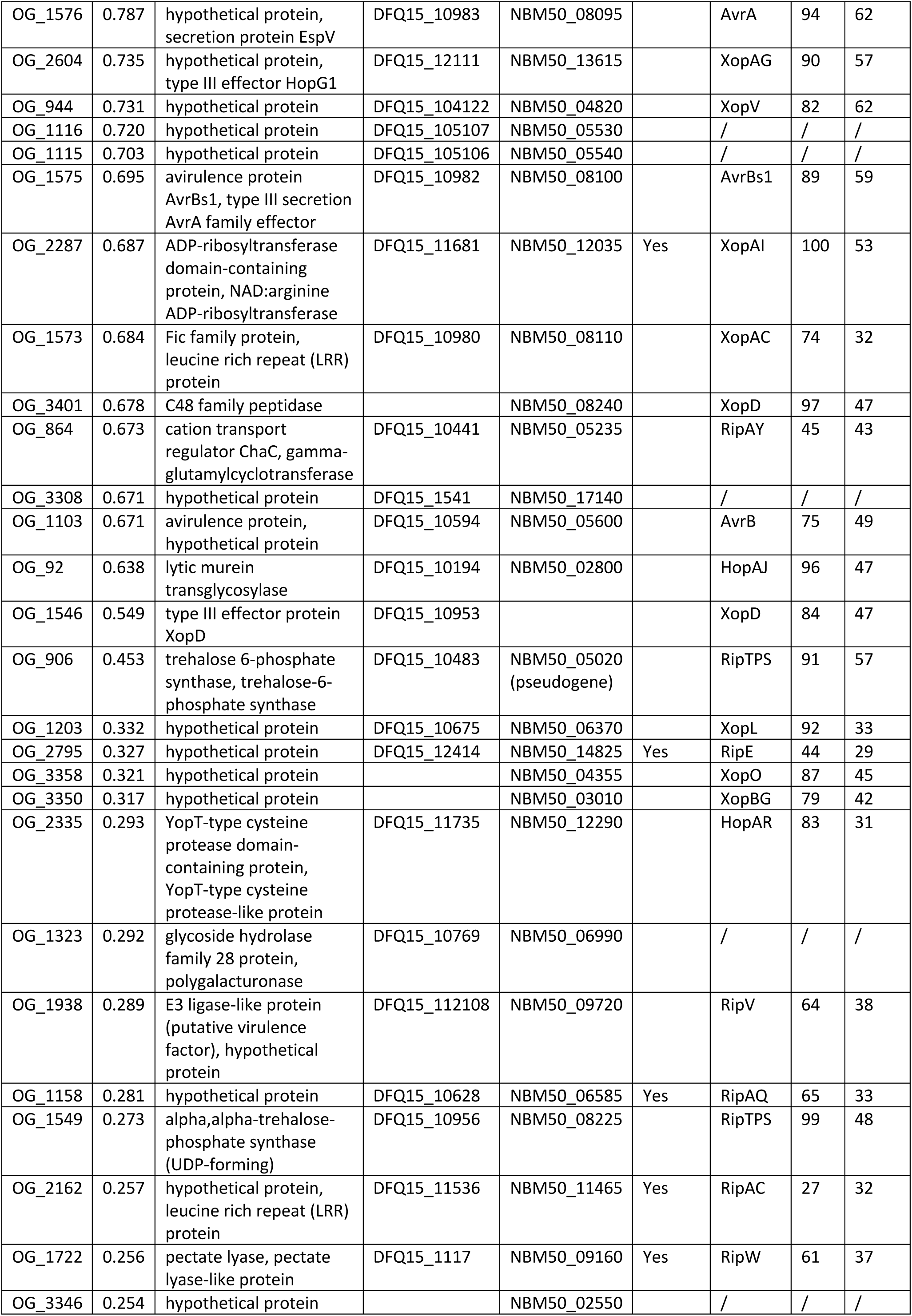
Prediction of type III-secreted proteins for SctV1-type T3SS of *X. ampelinus* using Effectidor II. Best hits with at least 25% coverage are shown. Candidate effectors with a corresponding PIP pattern (Table 2) are indicated. Corresponding effector classes, as revealed by BLASTP searches against a plant-pathogen T3E database, are listed along with the coverage and sequence identity.

| OG | Score | Annotation | CECT7646 | CFBP 1192 | PIP pattern | Effector class | Cov. | Ident. |
| --- | --- | --- | --- | --- | --- | --- | --- | --- |
| OG_1112 | 0.899 | hypothetical protein | DFQ15_105103 | NBM50_05555 | Yes | RipC | 100 | 67 |
| OG_671 | 0.898 | hypothetical protein, type III secretion system effector protein | DFQ15_10390 | NBM50_03725 |  | XopN | 83 | 60 |
| OG_2572 | 0.886 | hypothetical protein | DFQ15_12045 | NBM50_13085 |  | HopD | 75 | 26 |
| OG_222 | 0.832 | inosine-uridine preferring nucleoside hydrolase, nucleoside hydrolase | DFQ15_101225 | NBM50_02130 | Yes | HopQ | 100 | 66 |
| OG_1577 | 0.819 | hypothetical protein, inositol hexakisphosphate | DFQ15_10984 | NBM50_08090 |  | XopH | 97 | 64 |
| OG_1600 | 0.817 | hypothetical protein | DFQ15_109107 | NBM50_07980 |  | XopX | 83 | 63 |
| OG_576 | 0.801 | hypothetical protein | DFQ15_102289 | NBM50_01715 | Yes | XopZ | 90 | 41 |
| OG_297 | 0.800 | avirulence protein, hypothetical protein | DFQ15_1023 | NBM50_00260 |  | AvrB | 96 | 64 |
| OG_1753 | 0.794 | hypothetical protein | DFQ15_11138 | NBM50_09320 |  | RipA | 56 | 25 |
| OG_1576 | 0.787 | hypothetical protein, secretion protein EspV | DFQ15_10983 | NBM50_08095 |  | AvrA | 94 | 62 |
| OG_2604 | 0.735 | hypothetical protein, type III effector HopG1 | DFQ15_12111 | NBM50_13615 |  | XopAG | 90 | 57 |
| OG_944 | 0.731 | hypothetical protein | DFQ15_104122 | NBM50_04820 |  | XopV | 82 | 62 |
| OG_1116 | 0.720 | hypothetical protein | DFQ15_105107 | NBM50_05530 |  | / | / | / |
| OG_1115 | 0.703 | hypothetical protein | DFQ15_105106 | NBM50_05540 |  | / | / | / |
| OG_1575 | 0.695 | avirulence protein AvrBs1, type III secretion AvrA family effector | DFQ15_10982 | NBM50_08100 |  | AvrBs1 | 89 | 59 |
| OG_2287 | 0.687 | ADP-ribosyltransferase domain-containing protein, NAD:arginine ADP-ribosyltransferase | DFQ15_11681 | NBM50_12035 | Yes | XopAI | 100 | 53 |
| OG_1573 | 0.684 | Fic family protein, leucine rich repeat (LRR) protein | DFQ15_10980 | NBM50_08110 |  | XopAC | 74 | 32 |
| OG_3401 | 0.678 | C48 family peptidase |  | NBM50_08240 |  | XopD | 97 | 47 |
| OG_864 | 0.673 | cation transport regulator ChaC, gamma-glutamylcyclotransferase | DFQ15_10441 | NBM50_05235 |  | RipAY | 45 | 43 |
| OG_3308 | 0.671 | hypothetical protein | DFQ15_1541 | NBM50_17140 |  | / | / | / |
| OG_1103 | 0.671 | avirulence protein, hypothetical protein | DFQ15_10594 | NBM50_05600 |  | AvrB | 75 | 49 |
| OG_92 | 0.638 | lytic murein transglycosylase | DFQ15_10194 | NBM50_02800 |  | HopAJ | 96 | 47 |
| OG_1546 | 0.549 | type III effector protein XopD | DFQ15_10953 |  |  | XopD | 84 | 47 |
| OG_906 | 0.453 | trehalose 6-phosphate synthase, trehalose-6-phosphate synthase | DFQ15_10483 | NBM50_05020 (pseudogene) |  | RipTPS | 91 | 57 |
| OG_1203 | 0.332 | hypothetical protein | DFQ15_10675 | NBM50_06370 |  | XopL | 92 | 33 |
| OG_2795 | 0.327 | hypothetical protein | DFQ15_12414 | NBM50_14825 | Yes | RipE | 44 | 29 |
| OG_3358 | 0.321 | hypothetical protein |  | NBM50_04355 |  | XopO | 87 | 45 |
| OG_3350 | 0.317 | hypothetical protein |  | NBM50_03010 |  | XopBG | 79 | 42 |
| OG_2335 | 0.293 | YopT-type cysteine protease domain-containing protein, YopT-type cysteine protease-like protein | DFQ15_11735 | NBM50_12290 |  | HopAR | 83 | 31 |
| OG_1323 | 0.292 | glycoside hydrolase family 28 protein, polygalacturonase | DFQ15_10769 | NBM50_06990 |  | / | / | / |
| OG_1938 | 0.289 | E3 ligase-like protein (putative virulence factor), hypothetical protein | DFQ15_112108 | NBM50_09720 |  | RipV | 64 | 38 |
| OG_1158 | 0.281 | hypothetical protein | DFQ15_10628 | NBM50_06585 | Yes | RipAQ | 65 | 33 |
| OG_1549 | 0.273 | alpha,alpha-trehalose-phosphate synthase (UDP-forming) | DFQ15_10956 | NBM50_08225 |  | RipTPS | 99 | 48 |
| OG_2162 | 0.257 | hypothetical protein, leucine rich repeat (LRR) protein | DFQ15_11536 | NBM50_11465 | Yes | RipAC | 27 | 32 |
| OG_1722 | 0.256 | pectate lyase, pectate lyase-like protein | DFQ15_1117 | NBM50_09160 | Yes | RipW | 61 | 37 |
| OG_3346 | 0.254 | hypothetical protein |  | NBM50_02550 |  | / | / | / |

The situation was much different for the SctV2-type T3SS, which was found in strains GOD-11R, KACC 21265 and Kf1. Effectidor II predicted only six orthology groups with scores higher than 0.25 (Table 4). Four orthology groups had homologs in the T3E database (HopAJ, RipA, RipL, RipTPS), and each strain had two or three of these T3E candidates.

**Table 4.** Prediction of type III-secreted proteins for SctV2-type T3SS of *Xylophilus* using Effectidor II. Best hits with at least 25% coverage are shown. Corresponding effector classes, as revealed by BLASTP searches against a plant-pathogen T3E database, are listed along with the coverage and sequence identity.

| OG | Score | Annotation | GOD-11R | KACC 21265 | Kf1 | Effector class | Cov. | Ident. |
| --- | --- | --- | --- | --- | --- | --- | --- | --- |
| OG_92 | 0.638 | lytic murein transglycosylase | R9X41_13525 | GT347_03245 | GN316_20430 | HopAJ | 98 | 49 |
| OG_9901 | 0.285 | hypothetical protein |  |  | GN316_16015 | RipA | 83 | 30 |
| OG_1549 | 0.273 | alpha,alpha-trehalose-phosphate synthase (UDP-forming) | R9X41_19915 | GT347_17110 | GN316_10830 | RipTPS | 99 | 48 |
| OG_5313 | 0.259 | hypothetical protein | R9X41_16315 |  |  | / | / | / |
| OG_7634 | 0.255 | hypothetical protein |  | GT347_22755 |  | / | / | / |
| OG_7234 | 0.251 | hypothetical protein |  | GT347_17980 |  | RipL | 67 | 32 |

### Confirmation and structural modeling of predicted type III effectors

We next aimed to experimentally verify the secretion signal of novel effector candidates. Based on the high Effectidor II scores, BLASTP and InterProScan analyses, we focused on three orthology groups, OG_1116, OG_1115 and OG_3308, with Effectidor II scores of 0.720, 0.703 and 0.671, respectively (Table 3). Members of the other orthology groups had a substantially lower Effectidor II score (below 0.3) and had either no homologs beyond *Xylophilus* (OG_5313, OG_7634) or well-defined homologs that are not considered to be T3Es (OG_1323, OG_3346) (Tables 3 and 4). Members of OG_1323 are annotated as a glycoside hydrolase family 28 protein or a polygalacturonase, proteins that are typically secreted by the type II secretion pathway. Indeed, SignalP predicts a Sec/SPI-type Signal Peptide (Sec/SPI) with a likelihood of 0.999 for the 629-aa protein PYE78419 with the locus ID DFQ15_10769. OG_3346 has close homologs with more than 60% sequence identity that are annotated as cytosine-specific methyltransferases and that are encoded on *Acidovorax* bacteriophages (UYL85549.1, UYL85448.1). This observation prompted us to consider that the locus NBM50_02550 may belong to a prophage. To test this, we used geNomad to assess whether this locus is part of a mobile genetic element (Camargo et al. 2024), which was indeed the case. NBM50_02550 corresponds to one out of 58 ORFs that are encoded on a predicted 46-kb prophage (region 142964 to 188749 of contig QJTC01000001.1). For these reasons, we did not further study the orthology groups OG_5313, OG_7634, OG_1323, and OG_3346.

Three orthology groups, OG_1116 (DFQ15_105107, NBM50_05530), OG_1115 (DFQ15_105106, NBM50_05540) and OG_3308 (DFQ15_1541, NBM50_17140), corresponded to proteins that are similar to each other. However, a multiple sequence alignment revealed that the proteins belonging to OG_3308 lack the C terminus, which can be explained by the fact that the gene is located at the end of the contigs in both strains, CFBP 1192 (contig ID: JAMOFZ010000052.1) and CECT7646 (contig ID: QJTC01000054.1). The full-length versions of OG_1116 and OG_1115 have amino acids sequences that are 58% identical to each other. The C-terminally truncated version of OG_3308 is 54% identical to OG_1116 and 80% identical to OG_1115.

In total, there are twelve homologous protein variants deposited at NCBI, corresponding to the two strains, CECT7646 and CFBP 1192, each with three paralogs, and each paralog either annotated by the submitters and re-annotated by NCBI (RefSeq versions predicted with GeneMarkS-2+). Their multiple sequence alignment revealed that three different translation start codons were used (Supplementary Data S3). Based on sequence comparisons and the presence of a putative Shine-Dalgarno sequence, GGAG, with a distance of six base pairs to the ATG translation initiation codon, we considered the protein versions starting with MKDLYPF (Met-Lys-Asp-Leu-Tyr-Pro-Phe) as the most likely ones. The G+C content of the coding sequences is 56.7% for OG_1115, 56.9% for OG_1116 and 53.8 for OG_3308, compared to an overall G+C content of the genome of 67.8%. Such atypical G+C content can be taken as evidence for horizontal gene transfer, a process that is known to be involved in shaping the T3E repertoires of bacterial pathogens (Dillon et al. 2019).

We employed a reporter fusion system where the candidate effector’s type III secretion signal is translationally fused the defense-triggering domain of AvrBs1, a T3E from *Xanthomonas campestris* (Zhao et al. 2013). Our chimeric constructs consisted of the 100 base pairs upstream of the most likely translation initiation codon and the first 100 codons of the CDS, fused to the *avrBs1* CDS covering codons 59 to 445. When introduced into the *X. campestris* reporter strain 8004β*avrBs1*, bacteria expressing AvrBs1 reporter fusions triggered a *Bs1*-dependent hypersensitive response in pepper plants of the cultivar ECW-10R (Figure 4). In contrast, plants showed no reaction upon infiltration with bacteria that expressed the chimeric proteins but were mutated in the essential T3SS gene *hrcV*, thus demonstrating that the N-terminal 100 amino acids functioned as a type III secretion/translocation signal.

**Figure 4.**
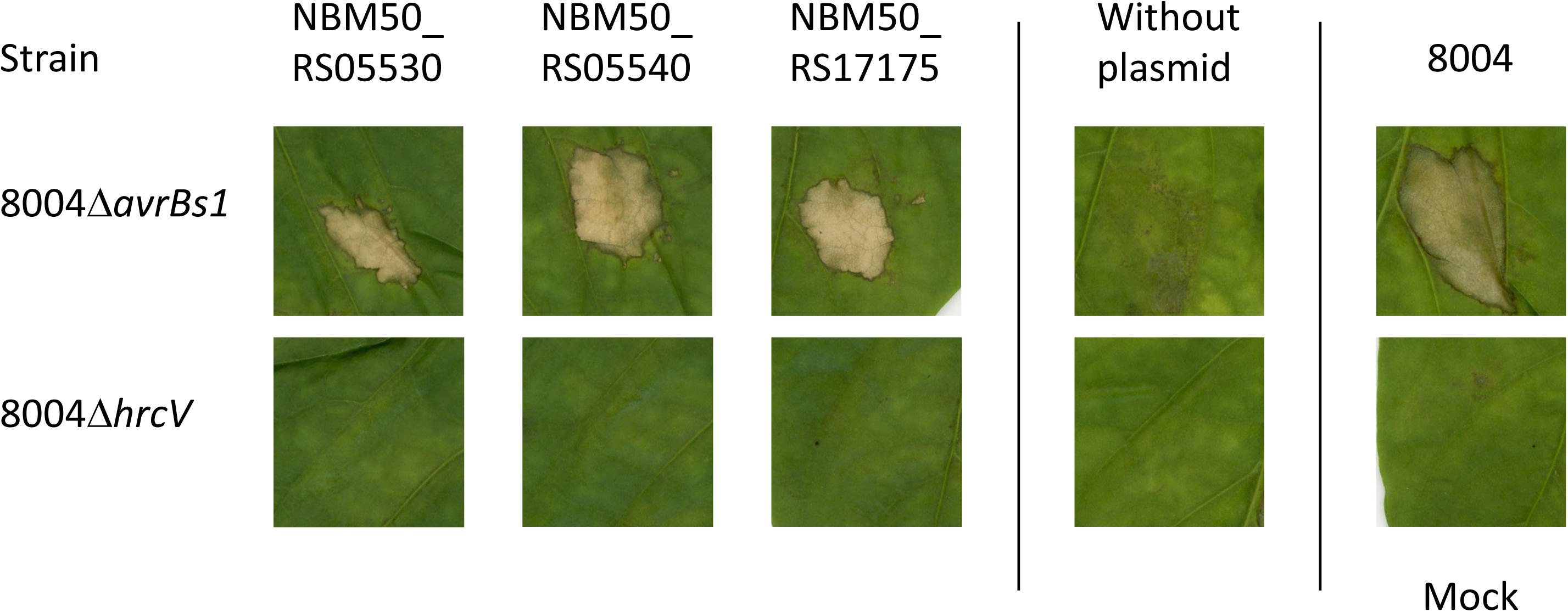
Translocation assays of novel T3E AvrBs1 reporter protein fusions. Bacterial *Xanthomonas* cells were inoculated into leaves of pepper cv. ECW-10R using needleless 1-mL plastic syringe. Strain 8004, which carries a copy of the avrBs1 gene and can elicit HR on non-host pepper cv. ECW-10R (Xu et al. 2008), was the positive control. Strains 8004Δ*avrBs1*, 8004Δ*hrcV* and mock inoculation with the buffer served as negative controls. The results presented are from a representative experiment, with similar results obtained in all other experiments.

To gain insight into the function and conservation of the novel T3Es, we used BLASTP to identify homologous sequences in the ClusteredNR database at NCBI GenBank. Sequences corresponding to OG_1115 and OG_1116 had 70 unique hits with an e-value lower than 10^−10^ (Supplementary Table S5). Most of these sequences were annotated as hypothetical proteins and they originated from *Pseudomonas savastanoi*, *Pseudomonas amygdali*, *Acidovorax* sp., *Pseudomonas syringae*, *Xanthomonas fragariae*, *Xanthomonas hydrangeae*, *Paraburkholderia aspalathi*, *Ralstonia* sp. and *Bradyrhizobium* sp., thus including both pathogenic as well as symbiotic bacteria. Notably, a few of the hits were annotated as C48 family peptidase or Ulp1 family isopeptidase. We then applied CD-Hit with a sequence identity threshold of 0.5 and a word size of 2 to cluster these sequences and established a set of 21 representative sequences. For functional predictions, these sequences were scanned at InterProScan (Supplementary Table S6). Five of the sequences had hits with InterPro entries corresponding to the Ulp1 protease family. However, the portions of OG_1115 and OG_1116 that aligned with these sequences did not correspond to the Ulp1 protease motifs.

To obtain further insight into the molecular function of the novel T3Es, we modeled the 3D structure of the two full-length versions with AlphaFold3 (Abramson et al. 2024) (Supplementary Figure S3). The predicted template modeling (pTM) scores were 0.39 for OG_1115 and 0.36 for OG_1116. Notably, the N-termini were largely predicted to be disordered (135 aa for OG_1115 and 109 aa for OG_1116), as was the case for the InterProScan predictions (regions 24-67 and 88-124 for OG_1115, region 16-116 for OG_1116), or they would include alpha-helices with very low confidence (pIDDT scores < 50). Structural predictions focusing on the apparently folded C-terminal domains, OG_1115_149-636_ and OG_1116_117-598_, improved the overall confidence of the predictions slightly to 0.42 and 0.40, respectively.

The predicted models were then used to find structural homologs, using FoldSeek (van Kempen et al. 2024). This way, a few T3Es (e.g., HopAV1 from *Pseudomonas amygdali*, HsvG and HsvB from *Pantoea agglomerans*), but also several ubiquitin-like protease domain-containing proteins from rhizobia (*Mesorhizobium*, *Bradyrhizobium*, *Rhizobium* and *Sinorhizobium*) were found (Supplementary Table S7). Most hits in other plants pathogens beyond *Xylophilus* had lower e-values (between 2.3 × 10^−59^ and 1.7 × 10^−11^) than hits in symbiotic bacteria (e-values above 3.6 × 10^−12^). Thus, additional experimental work is needed to better elucidate the function(s) of these newly identified effectors.

## DISCUSSION

Based on genomic data, we aimed to perform a genus-wide prediction of type III secretion and flagellar systems in *Xylophilus*, including the grapevine pathogen *Xylophilus ampelinus*. First, we curated the data available in NCBI to resolve taxonomic uncertainties. We found 19 genome sequences at NCBI GenBank, six of which were annotated as *X. ampelinus* and one of which was annotated as *X. rhododendri*. Based on average nucleotide identities, however, only two strains (CECT7646, CFBP 1192) corresponding to the same original isolate belong to the species *X. ampelinus*, while the genomes of BgEED09, bin23, CCH5-B3, and ERR594292 do not belong to this species. Based on ANI and DDH, isolates BgEED09, CCH5-B3 and ERR594292 do not even belong to the genus *Xylophilus*, since they are genetically closer to three type strains of the genus *Xenophilus* (Blümel et al. 2001; Kim et al. 2010; Li et al. 2014). Only the isolate bin23 appears to belong to the genus *Xylophilus* and may represent a new, yet uncharacterized species within this genus. Altogether, the remaining 16 genome sequences would correspond to ten different species, among which only two, *X. ampelinus* and *X. rhododendri*, have been formally described (Willems et al. 1987b; Lee et al. 2020).

Since many Gram-negative pathogenic and symbiotic bacteria rely on a T3SS and its set of secreted effectors for their interaction with eukaryotic organism, we scrutinized all twelve good-quality genome sequences of the genus *Xylophilus* for the presence of such a protein secretion system and the evolutionary related flagellar system. Firstly, we found three different flagellar systems in the genus, one of which was present in all twelve strains and is thus vertically inherited. The other two flagellar systems were only detected in one or two strains, and it remains to be analyzed whether the systems were lost in the other strains of the genus or if they were acquired by horizontal gene transfer in this subset of strains.

Based on sequence similarities and supported by synteny, we identified two types of T3SSs, both of which are classified as Hrp2/Hrc2-type T3SSs. Genes of this secretion system are under transcriptional control of the AraC-type transcriptional activator HrpX in *Acidovorax* and *Xanthomonas* and its ortholog HrpB in *Burkholderia* and *Ralstonia* (Genin et al. 1992; Wengelnik et al. 1996). We identified homologs of HrpX/HrpB in the T3SS gene clusters of *Xylophilus*, and we also identified the HrpX/HrpB-binding motif, the *hrpII*/PIP box (Cunnac et al. 2004a; Koebnik et al. 2006), in the promoters of several T3SS operons in *Xylophilus*, suggesting a similar regulatory mechanism to that in the well-studied plant pathogens mentioned above. However, we were unable to identify a bona fide ortholog of the upstream regulatory component HrpG (Teper et al. 2021), although each of the various *Xylophilus* genomes encodes dozens of response regulators. Notably, we found two paralogs of HrpB/HrpX genetically linked to the T3SS gene cluster in *X. ampelinus*, one of which is encoded immediately upstream of *sctC* and the other, which has 53% sequence identity, is encoded seven genes downstream of *hpaB*. It remains to be determined how transcription of the *hrpB*/*hrpX* genes is controlled in *Xylophilus*.

T3Es, which are secreted by the T3SS, need assistance in entering the eukaryotic target cell. For Hrp2/Hrc2-type T3SS, two types of putative translocons have been described, HrpF/NolX and HpaT (Büttner et al. 2002; Pesce et al. 2017). The gene upstream of the *X. ampelinus hrpB1* gene appears to encode a protein that is distantly related to the putative translocon HrpF/NolX, as supported by PSI-BLAST analysis and structural modeling (data not shown). However, we were unable to identify a putative translocon protein for the other T3SS, which is present in GOD11-R, Kf1, and KACC 21265, and it remains to be determined whether this T3SS injects effector proteins into eukaryotic target cells.

The presence of HrpB/HrpX homologs prompted us to predict, using a very conservative approach, genes that are very likely under transcriptional control of HrpB/HrpX and thus co-regulated with the T3SS genes. The power of this approach is underlined by the fact that four out of 16 predicted promoters in *X. ampelinus* control the expression of components or helpers of the T3SS (Hpa2, Hpa3, SctD, SctU) and another ten promoters control candidate T3E genes (*hopQ*, *ripC*, *ripE*, *ripW*, *ripAC*, *ripAQ*, *xopD*, *xopZ*, *xopAI*, *xopBG*). Two additional patterns were found within a CDS or in the antisense direction of a gene and likely represent false positives. For *X. rhododendri*, 35 HrpB/HrpX-regulated promoters were predicted, among which three are found in a region of 1,040 bp upstream of the *sctQ* CDS and two belong to genes with remote sequence homology to T3Es (XopN, RipL). Surprisingly, the remaining thirty promoters would all control the expression of hypothetical proteins, most of them without homologs beyond *X. rhododendri* and many of them belonging to gene families with several, sometimes distantly related, paralogs. It is unknown whether they correspond to T3Es or if they are involved in the interaction with other organisms.

Using a pan-genomic AI-based algorithm, we predicted candidate T3Es for *X. ampelinus* and *X. rhododendri*. In this way, we identified 31 homologs of Hop, Rip and Xop proteins from *Pseudomonas*, *Ralstonia* and *Xanthomonas* in *X. ampelinus*, which collectively form a large repertoire of candidate T3Es. This situation resembles other highly virulent plant-pathogenic bacteria, which use their effector repertoires, also called the effectome or effectorome, to successfully infect their host plants (Landry et al. 2020; Bundalovic-Torma et al. 2022). By contrast, we only predicted four Hop or Rip homologs in *X. rhododendri*, which raises questions about its pathogenic potential. Indeed, strain KACC 21265 was isolated from a rhododendron flower and there is no evidence that it would cause symptoms on rhododendron or other plants.

Three of the predicted T3Es of *X. ampelinus*, for which no homolog was found in T3E databases of *Pseudomonas*, *Ralstonia* and *Xanthomonas*, were found to possess a functional type III secretion signal that enabled the secretion and translocation of an AvrBs1 reporter protein from *Xanthomonas* into pepper plants. This shows that heterologous expression in a gamma proteobacterium can be used to validate candidate effectors from a beta proteobacterium. There are few examples for such trans-class secretion experiments. For instance, *Xanthomonas euvesicatoria* pv. *euvesicatoria* (syn. *X. campestris* pv. *vesicatoria*) was able to secrete PopA, a type III-secreted protein from *R. solanacearum* (Rossier et al. 1999). While these secretion experiments were tested for secretion using the same class of T3SS, here the Hrp2/Hrc2 class, other studies have shown that T3Es can even be secreted by a different T3SS class, at least when overexpressed. For instance, *X. euvesicatoria* pv. *euvesicatoria* could secrete AvrB, a Hrc1-class type III-secreted avirulence protein from *P. syringae* pv. *glycinea*, and YopE, a Ysc-class type III-secreted cytotoxin of the mammalian pathogen *Yersinia pseudotuberculosis* (Rossier et al. 1999). Similarly, Olaf Schneewind and colleagues reported that the Hrc1-class T3SS of *Erwinia chrysanthemi* recognized the secretion signals of the Ysc-class secreted proteins YopE and YopQ from *Yersinia*, and that the Hrc1-class secreted avirulence proteins AvrB and AvrPto from *P. syringae* could be secreted by the Ysc-class T3SS of *Yersinia* (Anderson et al. 1999). Such successful reciprocal secretion of proteins by different classes of T3SSs may suggest universal recognition of secretion signals. However, due to the lack of systematic studies and the fact that unsuccessful examples of heterologous secretion attempts are unlikely to be published, it remains unclear whether these examples are the exception or the rule. The discovery of effector-or effector-class-specific chaperones and translocation motifs suggests that effectors may not be universally recognized or translocated (Scheibner et al. 2018; Drehkopf et al. 2022; Pintor et al. 2025). Nevertheless, even if reciprocal secretion is not the rule, it provides a useful and straightforward strategy to test candidate effectors, as shown in this study.

The three candidate effectors from *X. ampelinus*, which were experimentally confirmed to contain a functional N-terminal type III secretion/translocation signal, share structural similarity with proteins of other plant-associated bacteria, including both pathogens (*Acidovorax* sp., *Pantoea agglomerans*, *P. amygdali*, *P. syringae*, *Ralstonia* sp., *Xanthomonas fragariae*) as well as symbionts (*Bradyrhizobium* sp.). Some of the proteins with similar structure are annotated as Ulp1 family isopeptidases or C48 family peptidases, known to be T3Es. Yet, there is no evidence that the *X. ampelinus* effectors have such a peptidase activity because the peptidase domains of these effectors do not align with the *X. ampelinus* amino acid sequence. The other homologs, which are annotated as hypothetical proteins, might also correspond to T3Es. Further studies need to determine how these effectors contribute to the infection of grapevine plants. Even more work is needed to understand the function of the other Hrp2/Hrc2-class T3SS, its candidate effectors and co-regulated gene families, as exemplified by the *X. rhododendri* strain KACC 21265.

## ACKNOWLEDGEMENTS

We thank Sebastien Santini (CNRS/AMU IGS UMR7256, France) and the PACA Bioinfo platform for the availability and management of the phylogeny.fr website used to generate phylogenetic trees (Figure 1, Supplementary Figure S1).

This work was supported by the Chateaubriand fellowships program of the French government to NW. NW was supported in part by a fellowship from the Edmond J. Safra Center for Bioinformatics at Tel Aviv University.

---

**Supplementary Figure S1.** Phylogenetic tree of representative SctU/FlhB proteins and homologs from *Xylophilus*. *Xylophilus* sequences are boxed with the three flagellar proteins in yellow (type 1), orange (type 2) and red (type 3), and the Sct proteins in green (T3SS#1) and blue (T3SS#2). Strain CFBP 1192 was omitted because it’s a clone of strain CECT7646. Sequences that were used to identify SctV/FlhA homologs in proteomes of *Xylophilus* are indicated by a blue dot.

**Supplementary Figure S2.** Comparison of *Xylophilus* flagellar gene clusters. Corresponding regions, extracted from GenBank files, were compared and visualized with Clinker. Three types of flagellar gene clusters were observed, as supported by synteny and sequence similarities. Flagellar genes are indicated as follows: flagellin genes in yellow, *flg* genes in red, *flh* genes in purple, *fli* genes in blue, *mot* genes in green and *che* genes in orange.

**Supplementary Figure S3.** Three-dimensional models of the tertiary structure of members of orthology groups OG_1115 and OG_1116. Snapshots of the predicted structure, coloured according to the pIDDT confidence scores, and the expected position errors are shown.

**Supplementary Table S1.** (a) Taxonomic classification of *Xylophilus* genome sequences using TYGS. (b) Taxonomic classification of *Xylophilus* genome sequences using MiGA.

**Supplementary Table S2.** (a) Genetic relationships between *Xylophilus* genomes based on genome-wide average nucleotide identities. (b) Genetic relationships between *Xylophilus* genomes based on digital DNA-DNA hybridization.

**Supplementary Table S3.** Predicted protein secretion systems in *Xylophilus* genomes using TXSScan.

**Supplementary Table S4.** HrpX-regulated protein families in *Xylophilus rhododendri*.

**Supplementary Table S5.** BLASTP hits of novel T3Es (OG_1115 and OG_1116).

**Supplementary Table S6.** InterPro hits of representation homologs of novel T3Es (OG_1115 and OG_1116).

**Supplementary Table S7.** FoldSeek hits of AlphaFold structural models of novel T3Es (OG_1115 and OG_1116).

**Supplementary Data S1.** Multiple sequence alignment of SctV/FlhA homologs in *Xylophilus*.

**Supplementary Data S2.** HrpX-regulated protein families in *Xylophilus rhododendri*.

**Supplementary Data S3.** Multiple sequence alignments of novel T3Es (OG_1115, OG_1116, OG_3308).

## Notes

### Competing Interest Statement

The authors have declared no competing interest.

### Summary of Updates

This version of the manuscript has been revised to include supplemental material.

