## Supplementary figures and images for "Identification of protein secretion systems and type III effectors in wood-associated bacteria of the genus *Xylophilus*"

### Supplementary Figure S1

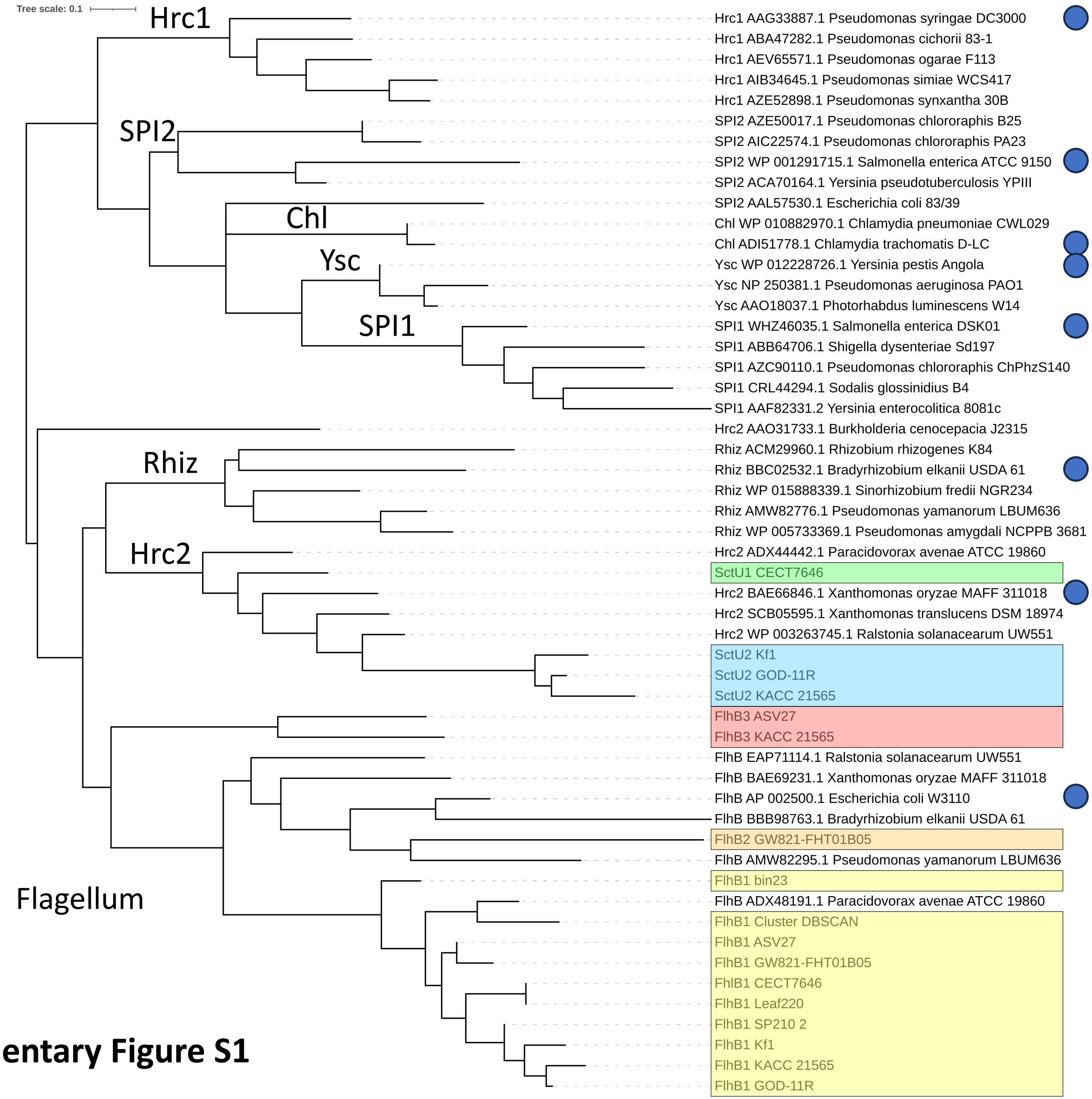


**Supplementary Figure S1**

### Supplementary Figure S2

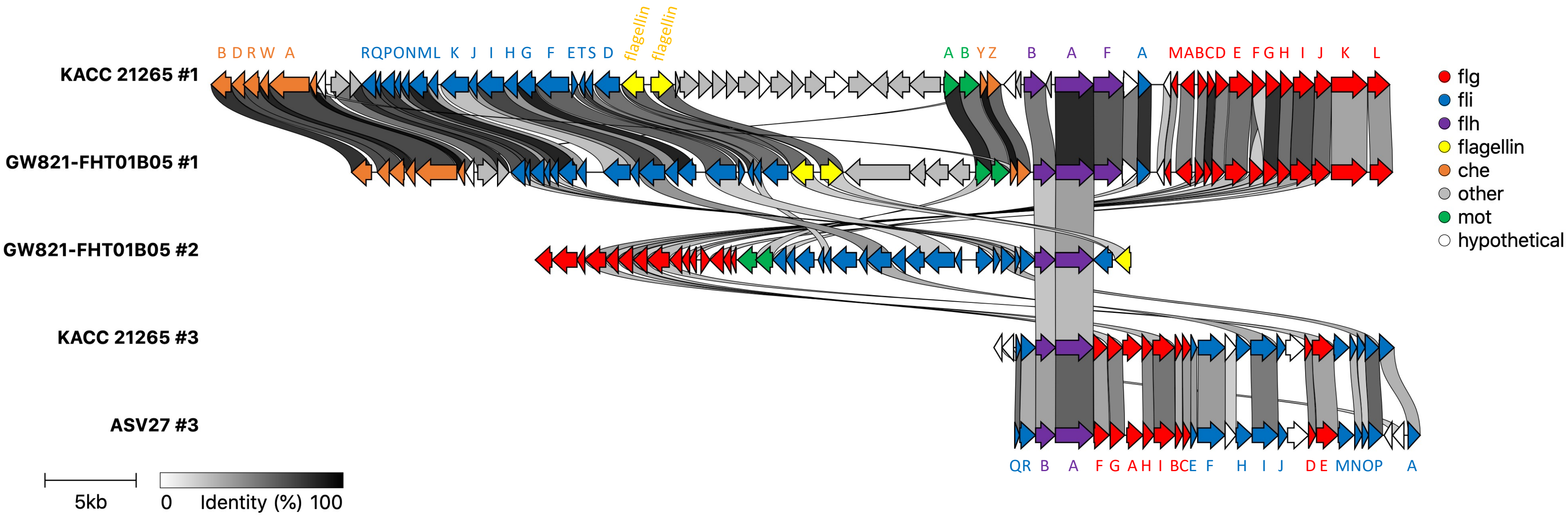


#### Supplementary Figure S2

### Supplementary Figure S3

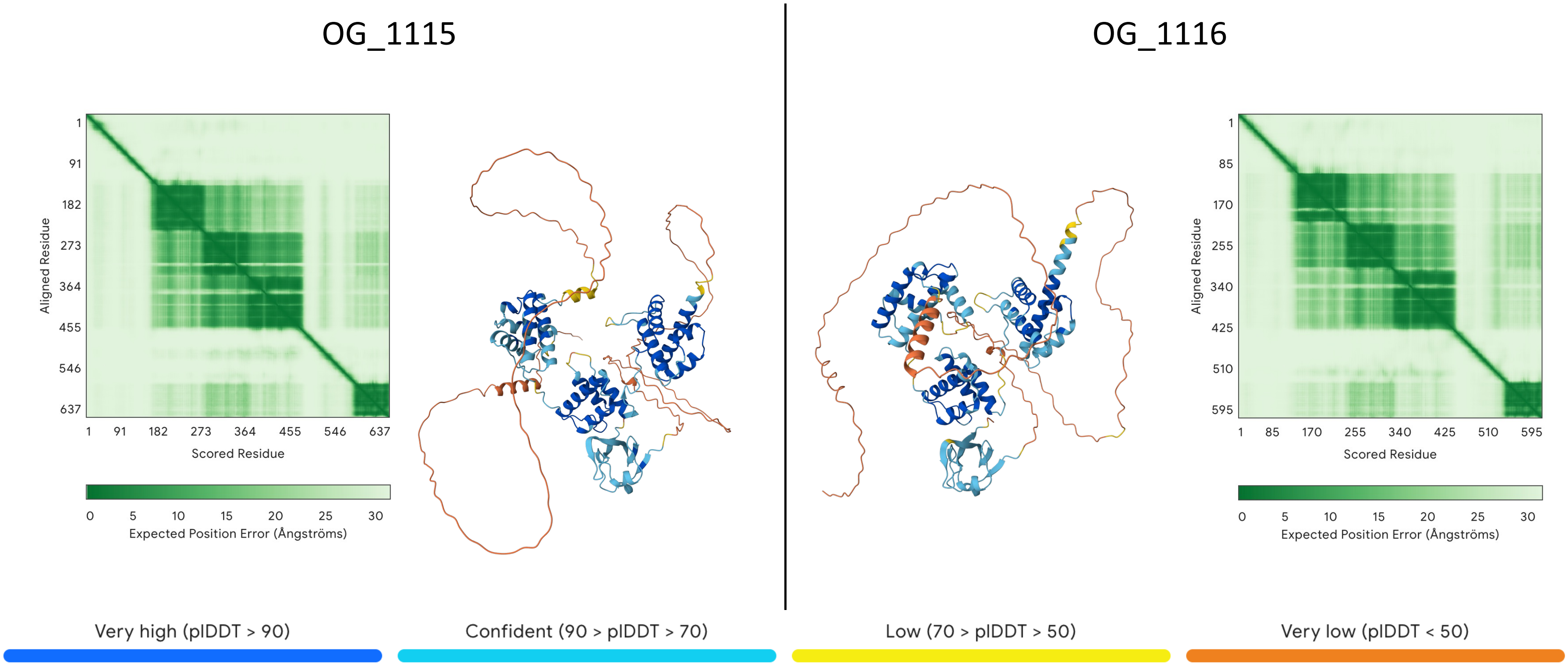


**Supplementary Figure S3**
