## Supplementary Data S1 for "Identification of protein secretion systems and type III effectors in wood-associated bacteria of the genus *Xylophilus*"

**Xylophilus**

```
CLUSTAL multiple sequence alignment by MUSCLE (3.8)

CECT7646_SctV1      -------------MAG---------------LVVAIVSLMILPLPTFAIDSLLAVNISVS
CFBP1192_SctV1      -------------MAG---------------LVVAIVSLMILPLPTFAIDSLLAVNISVS
Kf1_SctV2           MTPRKPPARARKPFGGELATA----G-----LVIAIIGLMILPLPPPVIDALLAVNITLS
GOD-11E_SctV2       MKA-------KRPFGGELATA----G-----LVIAIIGLMILPLPPPIIDALLAVNITLS
KACC21264_SctV2     MKA-------RRPFGGELATA----G-----LVIAIVGLMILPLPPPVIDALLAVNITLS
ASV27_FlhA3         MKI-------LRETFGRHS------DVSIVLLVLGVLVVLFAPIPSPLLDFLILTNFSFA
KACC21264_FlhA3     ----------MKQLFSRHGDVLL---IA---LVLGVLIVLFAPVPPALLDFLILTNFSFA
GW821_FlhA2         -----------MAYLTRLLAELRRHRFATPLFVLAILAMIILPLPPAVLDILFTFNIVLA
DBSCAN_FlhA1        MNARIAS---ARSWLGANTAQLQ--GIAAPILVVAILALMVLPMPSWMLDTFFTLNIAVA
SP210_2_FlhA1       MNTQVVR---ARAWMGRNAASLQ--GLAVPLFVVMILAMMVLPMPAWLLDVFFTLNIAIA
SP51_3_FlhA1        MNTQVVR---ARAWMGRNAASLQ--GLAVPLFVVMILAMMVLPMPAWLLDVFFTLNIAIA
KACC21264_FlhA1     MKARVTS---ARNWMGANGAAIK--GIAAPLFVVMILAMMVLPMPAWLLDTFFTINIAVA
Kf1_FlhA1           MKARVMT---ARNWMGANGEAIK--GIAAPLFVVMILAMMVLPMPAWLLDTFFTINIAVA
GOD-11E_FhlA1       MKARVMT---ARSWMGANGAAIK--GIAAPLFVVMILAMMVLPMPAWLLDVFFTLNIAIA
bin23_FlhA1         MNAQLLQ---ARNWLGRNAGLIQ--GVAAPLFVVMILAMMVLPMASWLLDVFFTINIAVA
CECT7646_FlhA1      MNATLGN---ARNWLGKNAAMIQ--GIAAPLFVVMILAMMVLPMPTWLLDVFFTINIAVA
CFBP1192_FlhA1      MNATLGN---ARNWLGKNAAMIQ--GIAAPLFVVMILAMMVLPMPTWLLDVFFTINIAVA
Leaf220_FlhA1       MNASLGN---ARNWLGKNAAMIQ--GIAAPLFVVMILAMMVLPMPTWLLDVFFTVNIAVA
ASV27_FlhA1         MNASLGQ---ARLWLGKNATMLQ--GVAAPLFVVMILAMMVLPMPPWLLDTFFTLNIAVA
GW821_FlhA1         MNASLGQLGQARLWLGKHATMLQ--GIAAPLFVVMILAMMVLPMPPWLLDTFFTINIAVA
                                                   :*: :: ::. *:..  :* ::  *: .:

CECT7646_SctV1      VLLLMTTLFIPNAIALSTFPSLLLFTTLFRLSLNIASTKSILLHADA-----GHVIETFG
CFBP1192_SctV1      VLLLMTTLFIPNAIALSTFPSLLLFTTLFRLSLNIASTKSILLHADA-----GHVIETFG
Kf1_SctV2           VLLLMSALYAPTAVSLSSFPSLLLFTTLLRLSLNIAATKAILLHGDA-----GHIIQSFG
GOD-11E_SctV2       VLLLMSALYAPTAVSLSSFPSLLLFTTLLRLSLNIAATKAILLHGDA-----GHIIESFG
KACC21264_SctV2     VLLLMSALYAPTAVSMSSFPSLLLFTTLLRLSLNIAATKAILLRGDA-----GHIIESFG
ASV27_FlhA3         LLILLLTFYMARPVEFSTFPSLLLVATLFRLSLNVAATRLILSEGDA-----GRVIAAIG
KACC21264_FlhA3     FLILLLTFYVARPVEFSTFPSLLLIATLFRLSLNVSATRLILSDASA-----GRVISAIG
GW821_FlhA2         LIVILVSVNSRRPLDFSVFPIVILATTLMRLTLNVASTRVVLLHGHEGTHAAGRVIEAFG
DBSCAN_FlhA1        LMVMMVAAYMLKPLDFAAFPAVLLLTTLMRLSLNVASTRVVLLEGHTGPSAAGAVIEAFG
SP210_2_FlhA1       LMVMMVAAYMVRPLDFAAFPAVLLFTTLMRLSLNVASTRVVLLEGHTGAGAAGAVIEAFG
SP51_3_FlhA1        LMVMMVAAYMVRPLDFAAFPAVLLFTTLMRLSLNVASTRVVLLEGHTGAGAAGAVIEAFG
KACC21264_FlhA1     LMVMMVAAYMVKPLDFAAFPLVLLLTTLMRLSLNVASTRVVLLEGHSGAGAAGAVIEAFG
Kf1_FlhA1           LMVMMVAAYMVKPLDFAAFPLVLLLTTLMRLSLNVASTRVVLLEGHTGAGAAGAVIEAFG
GOD-11E_FhlA1       LMVMMVAAYMVKPLDFAAFPLVLLLTTLMRLSLNVASTRVVLLEGHTGAGAAGAVIEAFG
bin23_FlhA1         LIVMMVAAYMVKPLDFVAFPSVLLLTTLMRLSLNVASTRVVLLEGHTGAGAAGAVIEAFG
CECT7646_FlhA1      LMVMMVAAYMVKPLDFIAFPSVLLLTTLMRLSLNVASTRVVLLEGHTGAGAAGAVIEAFG
CFBP1192_FlhA1      LMVMMVAAYMVKPLDFIAFPSVLLLTTLMRLSLNVASTRVVLLEGHTGAGAAGAVIEAFG
Leaf220_FlhA1       LMVMMVAAYMVKPLDFIAFPSVLLLTTLMRLSLNVASTRVVLLEGHTGAGAAGAVIEAFG
ASV27_FlhA1         LMVMMVAAYMVKPLDFVAFPAVLLLTTLMRLSLNVASTRVVLLEGHTGPGAAGAVIEAFG
GW821_FlhA1         LMVMMVAAYMIKPLDFIAFPSVLLLTTLMRLSLNVASTRVVLLEGHTGAGAAGAVIEAFG
                    .:::: :     .: :  ** ::* :**:**:**:::*. :*  .       * :* ::*

CECT7646_SctV1      ELVVGGNLVVGIVIFLIITIVQFIVISKGSERVAEVGARFTLDAMPGKQMSIDADLRAGL
CFBP1192_SctV1      ELVVGGNLVVGIVIFLIITIVQFIVISKGSERVAEVGARFTLDAMPGKQMSIDADLRAGL
Kf1_SctV2           ELVVGGNLLVGMTVFVIIAAVQFIVIAKGSERASEVSARFALDGLPGQQMSIDAELRAGS
GOD-11E_SctV2       NLVVGGNLLVGLTVFVIIAAVQFIVIAKGSERASEVSARFALDGLPGQQMSIDAELRAGS
KACC21264_SctV2     NLVVGGNLLVGMTVFVIIAAVQFIVIAKGSERASEVSARFALDGLPGQQMSIDAELRAGS
ASV27_FlhA3         SYVVGGNYVIGLIVFLILIVVQYVVVTSGAQRVSEVAARFTLDSMPGQQMSIDADLNMGF
KACC21264_FlhA3     SFVVGGNYVIGLIVFLILIVVQYVVVTSGAQRVSEVAARFTLDSMPGQQMSIDADLNMGF
GW821_FlhA2         NVVIGGNFVVGLVVFVILMIINFVVVTKGAERISEVSARFTLDALPGKQMAIDADLNAGL
DBSCAN_FlhA1        HFLIGGNFAVGLIVFAILVVINFVVVTKGAERIAEVGARFTLDAMPGKQMAVDADLNAGL
SP210_2_FlhA1       HFLIGGNFAVGLIVFAILVVINFVVVTKGSERIAEVSARFSLDAMPGKQMAVDADLNAGV
SP51_3_FlhA1        HFLIGGNFAVGLIVFAILVVINFVVVTKGSERIAEVSARFSLDAMPGKQMAVDADLNAGV
KACC21264_FlhA1     HFLIGGNFAVGLIVFAIVVVINFVVVTKGAERIAEVSARFTLDAMPGKQMAVDADLNAGV
Kf1_FlhA1           HFLIGGNFAVGLIVFAIVVVINFVVVTKGSERIAEVSARFTLDAMPGKQMAVDADLNAGV
GOD-11E_FhlA1       HFLIGGNFAVGLIVFAIVVVINFVVVTKGSERIAEVSARFTLDAMPGKQMAVDADLNAGV
bin23_FlhA1         HFLIGGNFAVGLIVFAILVVINFVVVTKGAERIAEVSARFTLDAMPGKQMAVDADLNAGL
CECT7646_FlhA1      HFLIGGNFAVGLIVFAILVVINFVVVTKGAERIAEVSARFTLDAMPGKQMAVDADLNAGL
CFBP1192_FlhA1      HFLIGGNFAVGLIVFAILVVINFVVVTKGAERIAEVSARFTLDAMPGKQMAVDADLNAGL
Leaf220_FlhA1       HFLIGGNFAVGLIVFAILVVINFVVVTKGAERIAEVSARFTLDAMPGKQMAVDADLNAGL
ASV27_FlhA1         HFLIGGNFAVGLIVFSILVVINFIVVTKGAERIAEVSARFTLDAMPGKQMAVDADLNAGL
GW821_FlhA1         HFLIGGNFAVGLIVFAILVVINFVVVTKGAERIAEVSARFTLDAMPGKQMAVDADLNAGL
                      ::***  :*: :* *:  ::::*::.*::* :**.***:**.:**:**::**:*. *
```

#### Supplementary Data S1

```
CECT7646_SctV1      LTGEEARKKRALLAMESQMHGGMDGAMKFVKGDTIAGLIITLVNLVAGIIVGVLYHNMTA
CFBP1192_SctV1      LTGEEARKKRALLAMESQMHGGMDGAMKFVKGDTIAGLIITLVNLVAGIIVGVLYHNMTA
Kf1_SctV2           MTADEARRKRAMLAMESKLHGSMDGAMKFVKGDAIAALVITIINICAGITIGVTYHDLAM
GOD-11E_SctV2       MTAEEARKKRAMLAMESKLHGSMDGAMKFVKGDAIAALVITIINICAGITIGVSYHDMSM
KACC21264_SctV2     MTADEARRKRAMLAMESKLHGSMDGAMKFVKGDAIAALVITIINICAGITIGVSYHEMSM
ASV27_FlhA3         IDQAEAQRRRKNVDKEAAFYGAMDGASKFVKGDAIAGIIILLIDIIGGLVIGVMQHKMAW
KACC21264_FlhA3     IDQEQAQARRRNLEKEAGFYGAMDGASKFVKGDAVAGIVIMLINIVGGLVIGVMQHGYGW
GW821_FlhA2         INQEKAQARRHEVASEADFYGAMDGASKFVRGDAIASILILIVNLVGGVAIGALMHDLSF
DBSCAN_FlhA1        IDEKEAKRRRAEVQEEANFFGSMDGASKFVRGDAIAGILILLINIVGGFVIGMLQHGLSA
SP210_2_FlhA1       IDEKEARKRRLEIGEEANFFGSMDGASKFVRGDAMAGILILLINIVGGFAIGVLQHGLSA
SP51_3_FlhA1        IDEKEARKRRLEIGEEANFFGSMDGASKFVRGDAMAGILILLINIVGGFAIGVLQHGLSA
KACC21264_FlhA1     IDEKEARKRRAEVGEEAAFFGSMDGASKFVRGDAMAGIIILLINIIGGFTIGVLQHDLTA
Kf1_FlhA1           IDEKEARKRRAEVGEEANFFGSMDGASKFVRGDAVAGIIILLINIIGGFAIGVLQHDLSA
GOD-11E_FhlA1       IDEKEARKRRAEVGEEANFFGSMDGASKFVRGDAMAGIIILLINIIGGFAIGVLQHDLTA
bin23_FlhA1         IDEATARKRRAEVGEEANFFGAMDGASKFVRGDAVAGILILLINIVGGFAIGMIQHGLSA
CECT7646_FlhA1      IDEATARKRRAEVGEEANFFGAMDGASKFVRGDAIAGILILLINIVGGFAIGMIQHNLSA
CFBP1192_FlhA1      IDEATARKRRAEVGEEANFFGAMDGASKFVRGDAIAGILILLINIVGGFAIGMIQHNLSA
Leaf220_FlhA1       IDEATARKRRAEVGEEANFFGAMDGASKFVRGDAIAGILILLINIVGGFAIGMIQHSLSA
ASV27_FlhA1         IDEATARKRRAEVGEEANFFGAMDGASKFVRGDAIAGILILLINIVGGFAIGVLQHDLSA
GW821_FlhA1         IDEATARKRRVEVGEEANFFGAMDGASKFVRGDAIAGILILLINIVGGFAVGVLQHNLSA
                    :    *. .*  :  *: :.*.**** ***.**::*.::* :::: .*. :*   *

CECT7646_SctV1      GEAANRFAVLSIGDAMVSQIASLFICVAAGVLITRVADDGEKKPRSLGQEVGAQITGNAR
CFBP1192_SctV1      GEAANRFAVLSIGDAMVSQIASLFICVAAGVLITRVADDGEKKPRSLGQEVGAQITGNAR
Kf1_SctV2           GAAAQKYSVLSIGDAMVSQIPSLLISVAAGVLTTRVLDTRTTKRDSLGTEVVRQLGGNAP
GOD-11E_SctV2       GLAAQKYSVLSIGDAMVSQIPSLLISVAAGVLTTRVLDTRTTKRDSLGQEVVRQLGGNAP
KACC21264_SctV2     AVAAQKYSVLSIGDAMVSQIPSLLISVAAGVLTTRVLDTRTAKRDSLGQEVVRQLGSSTP
ASV27_FlhA3         DQALQRYTLLTIGDGIVTQVPALVIAVGTGIIVTRS-----ASDGNLSQEVLRQITSFPK
KACC21264_FlhA3     SQALQTYTLMTIGDGIVTQVPALVIAVGTGIIVTRS-----SSDRNLSAEVLRQITSFPK
GW821_FlhA2         GDAFRQYALLTIGDGLVAQIPALLLSAAAAIIVTRI-----SDSGQFEQQVGSQLLASPT
DBSCAN_FlhA1        GQAADSYILLAVGDALVAQIPGLLISVAAAMVVSRV-----GKDTDMGGQIVGQLFTSPR
SP210_2_FlhA1       GKAADTYILLAVGDALVAQIPGLLISVAAAMVISRV-----GKESDVGGQIVDQLFMSPR
SP51_3_FlhA1        GKAADTYILLAVGDALVAQIPGLLISVAAAMVISRV-----GKESDVGGQIVDQLFMSPR
KACC21264_FlhA1     AKAADTYILLAVGDALVAQIPGLLISVAAAMVISRV-----GKENDVGQQVIAQLFNSPR
Kf1_FlhA1           SKAADTYILLAVGDALVAQIPGLLISVAAAMVISRV-----GKEKDVGGQVIGQLFNSPR
GOD-11E_FhlA1       GKAADTYILLAVGDALVAQIPGLLISVAAAMVISRV-----GKEKDVGGQVIGQLFNSPR
bin23_FlhA1         GQAADSYILLAVGDALVAQIPGLLISVAAAMVISRV-----GKEQDVGGQIIGQLFGSAK
CECT7646_FlhA1      GQAADSYILLAVGDALVAQIPGLLISVAAAMVISRV-----GKEQDVGGQIIGQLFNSPK
CFBP1192_FlhA1      GQAADSYILLAVGDALVAQIPGLLISVAAAMVISRV-----GKEQDVGGQIIGQLFNSPK
Leaf220_FlhA1       GQAADSYILLAVGDALVAQIPGLLISVAAAMVISRV-----GKEQDVGGQIIGQLFNSPK
ASV27_FlhA1         SQAASSYILLAIGDALVAQIPGLLISVAAAMVISRV-----GKEQDVGGQILGQLFNSPK
GW821_FlhA1         SQAADSYILLAVGDALVAQIPGLLISVAAAMVISRV-----GKEQDVGGQILGQLFNSPK
                      *   : ::::**.:*:*:..*.:...:.:: :*       .  ..  ::  *:   .

CECT7646_SctV1      AMFLAGALVLAFAAVPGFPV---LQFLAISTGLGAAG--WVLD---KRRRKQEQ------
CFBP1192_SctV1      AMFLAGALVLAFAAVPGFPV---LQFLAISTGLGAAG--WVLD---KRRRKQEQ------
Kf1_SctV2           ALYMSSLLLLGFAAVPGFPY---ALFLAAAAGLSFCA--WRLQ---HRAVAETE---GEE
GOD-11E_SctV2       AMYMSSLLLLGFAAVPGFPY---PLFLIASGGLAFAA--RRLQ---RQSAAGGPSTGEEG
KACC21264_SctV2     ALYMSSLLLVGFAAVPGFPY---PLFLTASAGLALCA--WRLQ---RGAAPGGE----GA
ASV27_FlhA3         TLLLVAAALLGLLILPGIPAWPALFLICMVGVAAWFA--YRAA---AERTADES--INAP
KACC21264_FlhA3     TLVLVAAVLGGLLLLPGLPLL--PSSLLIACLLGAAALSMRVR---ANAALQTQ------
GW821_FlhA2         VLYGAAGLMLALGLIPGMPW---PTFLCFAGALGYVA--WRMGQHLKRPAEADT------
DBSCAN_FlhA1        VLGVTAGILVLLGLIPGMPH---AVFLLIGALVGYAA--WVLA---RRNAAAAK------
SP210_2_FlhA1       VIGIAALILVLVGLIPGMPH---LVFLGIGGVLGYLA--WTLA---KRPPPADL------
SP51_3_FlhA1        VIGIAALILVLVGLIPGMPH---LVFLGIGGVLGYLA--WTLA---KRPPPADL------
KACC21264_FlhA1     VLGITGGILLSLGLIPGMPH---LVFLAIGGLVSYGA--WRMS---KRPPPPTP-EQLAA
Kf1_FlhA1           VLGITGAILLTLGIIPGMPH---VVFLAIGSGCAYMA--WSLS---KRPPPPDPKEVAAL
GOD-11E_FhlA1       VLGITGGILLTLGLIPGMPH---LVFLGIGGAVAYGA--WKLS---KRPPPPDPKQVEAA
bin23_FlhA1         VLSLTGGILILLGLIPGMPH---LVFLGVGGAISYFA--WRLH---KRPPPPDP----AA
CECT7646_FlhA1      VLGLTAGILILLGLIPGMPH---LVFLGVGGVVAWMT--WRLH---KRPPPPDP------
CFBP1192_FlhA1      VLGLTAGILILLGLIPGMPH---LVFLGVGGVVAWMT--WRLH---KRPPPPDP------
Leaf220_FlhA1       VLGLTAGILILLGLIPGMPH---LVFLGVGGGMAWMA--WRLH---KRPPPPDP------
ASV27_FlhA1         VLGLTAGILILLGLIPGMPH---AVFLVVGSVFGYLA--WRLY---KRPPPADP------
GW821_FlhA1         VLGLTAGILILLGLIPGMPH---AVFLGVGSAIAYMA--WRLH---KRPPPADP------
                    .:   .  :  .  :**:*       :      .
```

**Supplementary Data S1**

```
CECT7646_SctV1      ---VAISKPIGALQRDGAKGEPPSIRQEPPLFAKPLAVRMSRQLGMLLEADALDEAITTE
CFBP1192_SctV1      ---VAISKPIGALQRDGAKGEPPSIRQEPPLFAKPLAVRMSRQLGMLLEADALDEAITTE
Kf1_SctV2           PGEASDGRRMPSLRAYGAKGEAPSITDIAPSFASPLGLRLSEALSRRIAPATLDNAFRDA
GOD-11E_SctV2       GGEASEGGRMPSLRGYGAKGDAPSITENAPAFASPLGLRLSSAMVPRIEPAALDEAFKRE
KACC21264_SctV2     DAETSDGRRMPSLRGYGAKGEAPSITDAAPVFASPLGLRLSEAMVPRLQPAALDDAFRVA
ASV27_FlhA3         EDDIGEAAAAPAQDPYALL----PV----EPIEVHVGSHWIGLV--NQPGNVFMERITTF
KACC21264_FlhA3     DTPPQEENSAPDSAAAQAYESL-RV----PPVEVAIGSRWTGLAGDA--GP-LLPRIQEF
GW821_FlhA2         --AAVEAALLAAPPPDLDWRSLPFV----EQVTVTVGYKLASLLDTAHGAP-LAKRLKGI
DBSCAN_FlhA1        -AAAAAPPPAPAGDGEASWDDLQPV----DLLGLELGYRLITLVDKHRQGD-LLTRIKGV
SP210_2_FlhA1       ----QAEAPPAPSDGEASWDDLQPV----DLLGLELGYRLITLVDKNRQGD-LLNRIKGV
SP51_3_FlhA1        ----QAEAPPAPSDGEASWDDLQPV----DLLGLELGYRLITLVDKNRQGD-LLNRIKGV
KACC21264_FlhA1     AEAAASAQAAPGNDGEASWDDLQPV----DLLGLELGYRLIILVDKNRQGD-LLTRIKGV
Kf1_FlhA1           EAAAAAQAAAANSDGEASWDDLQPV----DLLGLELGYRLIILVDKNRQGD-LLTRIKGV
GOD-11E_FhlA1       EAAAAAQQAAASNDGEASWDDLQPV----DLLGLELGYRLIILVDKNRQGD-LLTRIKGV
bin23_FlhA1         AAAAAAAAAAPANDGEATWDDLQPV----DLLGLELGYRLITLVDKNRQGD-LLTRIKGV
CECT7646_FlhA1      ---AAAAAAAPPPDGEASWDDLQPV----DLLGLELGYRLITLVDKNRQGD-LLTRIKGV
CFBP1192_FlhA1      ---AAAAAAAPPPDGEASWDDLQPV----DLLGLELGYRLITLVDKNRQGD-LLTRIKGV
Leaf220_FlhA1       --AAAAAAAAPPPDGEASWDDLQPV----DLLGLELGYRLITLVDKNRQGD-LLTRIKGV
ASV27_FlhA1         ---AAAAAATALPDGEATWDDLQPV----DLLGLELGYRLIALVDKTRQGD-LLTRIKGV
GW821_FlhA1         AAAAAAAATTAQPDGEASWDDLQPV----DLLGLELGYRLIALVDKNRQGD-LLTRIKGV
                                            :      .   :. .             :   :

CECT7646_SctV1      RQRLQEELGLPFPGIAMWVTHNLEALEFDVLLNDVPTREVKLPGRMMLLMDP---DSPLA
CFBP1192_SctV1      RQRLQEELGLPFPGIAMWVTHNLEALEFDVLLNDVPTREVKLPGRMMLLMDP---DSPLA
Kf1_SctV2           RAALEEEYGIPFPGMAVWSSPLAGQRGYDVLVQDVPVAGGPLPPD---------------
GOD-11E_SctV2       RAALEEEYGVPFPGMAVWSSPSMTASGYDVLVQDVPM-----------------------
KACC21264_SctV2     RAALEEEYGVPFPGMAVWSSQMAEGRGYDVLVQDVPVE-----------------TRPL-
ASV27_FlhA3         RKQHAQDLGLVLPRVRFKDSARLGADRYEIHLDGAVCGRGEARSDRLLAIHPTGKTDILA
KACC21264_FlhA3     RRGFALELGMVLPPIRLRESSRLAPDRYEILIDGVACARAEVRQDRLLAIHPAGDVKSVP
GW821_FlhA2         RQNLSEAMGLLLPPIGLRDDLALKPSQYAVLLGGAAIARGEVFQDRLMAIPSPNVYGQLD
DBSCAN_FlhA1        RRKFAQEVGFLPPAVHVRDNLELKPSGYRITLRGVVVGEGEAFPGQFLAINPGGITTPLI
SP210_2_FlhA1       RRKFAQEVGFLPPAVHVRDNLELKPSGYRITLRGVMVGEGEAFPGMFLAINPGGISTPLI
SP51_3_FlhA1        RRKFAQEVGFLPPAVHVRDNLELKPSGYRITLRGVMVGEGEAFPGMFLAINPGGISTPLI
KACC21264_FlhA1     RRKFAQEVGFLPPAVHVRDNLELKPSGYRITLRGVVVGEGEAFPGMFLAINPGGISTPLI
Kf1_FlhA1           RRKFAQEVGFLPPAVHVRDNLELKPSGYRITLRGVVVGDGEAFPGMYLAINPGGITTPLI
GOD-11E_FhlA1       RRKFAQEVGFLPPAVHVRDNLELKPSGYRITLRGVVVGEGEAFPGMFLAINPGGISTPLI
bin23_FlhA1         RRKFAQEVGFLPPPVHVRDNLELKPSGYRITLRGVVVGEGEAFPGMYLAINPGGIATPLI
CECT7646_FlhA1      RRKFAQEVGFLPPAVHVRDNLELKPSGYRITLRGVVMGDGEAFPGMFLAINPGGITTPLI
CFBP1192_FlhA1      RRKFAQEVGFLPPAVHVRDNLELKPSGYRITLRGVVMGDGEAFPGMFLAINPGGITTPLI
Leaf220_FlhA1       RRKFAQEVGFLPPAVHVRDNLELKPSGYRITLRGVVMGDGEAFPGMFLAINPGGITTPLI
ASV27_FlhA1         RRKFAQEVGFLPPPVHVRDNLELKPSAYRITLRGVVVGEGEAFPGMYLAINPGGITTPLI
GW821_FlhA1         RRKFAQEVGFLPPAVHVRDNLELKPSGYRITLRGVVVGEGEAFPGMFLAINPGGITTPLI
                    *       *.  * : ..         : : : ..

CECT7646_SctV1      QRAERHGPVLDQPESLWIKDGALPEADRR--TVLSIENVIARHVTGALRQHAHMFMGVQE
CFBP1192_SctV1      QRAERHGPVLDQPESLWIKDGALPEADRR--TVLSIENVIARHVTGALRQHAHMFMGVQE
Kf1_SctV2           -------GATGLE-----------------------VALATETVAVLGRNAHLFVGIQE
GOD-11E_SctV2       -------QSRPLP--------PPDAKGLEGALAADAVAVLARN--------AHLFVGIQE
KACC21264_SctV2     -------PPPGLA---------------------QLETAVATDVVAVLARNAHLFVGIQE
ASV27_FlhA3         GEPTKD-PTYGLP-ALWIEEQQRDAAIAAKFTIVDAPTVFMTHLTEVLRRESATLLTRAE
KACC21264_FlhA3     GELTRD-PTYHLP-ALWIEESQRAAAQAARYTLVDGPTVFLTHLGEVLRREAATLLSRAE
GW821_FlhA2         GIPGTE-PAYALP-VTWIEAEHKAHALGLGYQVVDGPSVIATHLFKLLQDHLSELIRHED
DBSCAN_FlhA1        GTATTD-PAFGLP-AHWIDARQKEAAQMAGFTVVDSETVLATHLSHLMQVQAARLLSRTE
SP210_2_FlhA1       GTPTTD-PAFGLP-AHWIDERQKEAAQMAGFTVVDSETVMATHLSHLMQVQAARLLSRTE
SP51_3_FlhA1        GTPTTD-PAFGLP-AHWIDERQKEAAQMAGFTVVDSETVMATHLSHLMQVQAARLLSRTE
KACC21264_FlhA1     GTPTTD-PAFGLP-AHWIDERQKEAAQMAGFTVVDSETVMATHLSHLMQVQAAKLLSRTE
Kf1_FlhA1           GTPTTD-PAFGLP-AHWIDERQKEAAQMAGFTVVDSETVMATHLSHLMQVQAAKLLSRTE
GOD-11E_FhlA1       GTPTTD-PAFGLP-AHWIDERQKEAAQMAGFTVVDSETVMATHLSHLMQVQAAKLLSRTE
bin23_FlhA1         GTATTD-PAFGLP-AHWIDERQKEAAQMAGFTVVDSETVMATHLSHLMQVQAAKLLSRTE
CECT7646_FlhA1      GTATTD-PAFGLP-AHWIDERQKEAAQMAGFTVVDSETVMATHLSHLMQVQAAKLLSRTE
CFBP1192_FlhA1      GTATTD-PAFGLP-AHWIDERQKEAAQMAGFTVVDSETVMATHLSHLMQVQAAKLLSRTE
Leaf220_FlhA1       GTATTD-PAFGLP-AHWIDERQKEAAQMAGFTVVDSETVMATHLSHLMQVQAAKLLSRTE
ASV27_FlhA1         GTATTD-PAFGLP-AHWIDERQKEAAQMAGFTVVDSETVMATHLSHLMQVQAAKLLSRTE
GW821_FlhA1         GTPTTD-PAFGLP-AHWIDERQKEAAQMAGFTVVDSETVMATHLSHLMQVQAAKLLSRTE
                                                          ..              ::   :
```

### Supplementary Data S1

```
CECT7646_SctV1      VQWVMERVSAEYPGLVTEV-QKILPAQRIAEVLRRLLEEQVPVRNVRTILEGLIAWGPKE
CFBP1192_SctV1      VQWVMERVSAEYPGLVTEV-QKILPAQRIAEVLRRLLEEQVPVRNVRTILEGLIAWGPKE
Kf1_SctV2           TQWLFDRLNVDYPGLVTEV-QKVLPLQRVADVLRRLLEEQVPIRNLRSILESLVYWGPKE
GOD-11E_SctV2       TQWMFDKLNADYPGLITEV-QKVLPLQRVADVLRRLLEEQIPIRNLRSILESLVYWGPKE
KACC21264_SctV2     TQWMFDKLNADYPGLITEV-QKVLPLQRVADVLRRLLEEQIPIRNLRSILESLVYWGPKE
ASV27_FlhA3         TDRLLARVRQTQASLVEELIPTVLSVSDVQRVLQNLLREKVSIRHMEAILETLADAGRQS
KACC21264_FlhA3     VEKLLARVRGSQPGLVEDIVPTVLSLTDVQKVLQALLREKVSIRNLEAILETLADAGRHT
GW821_FlhA2         VPALLERLAAQAPKLSAAL-EKALTHSQLLKVFRVLLAELVPLRDIVPIATTLLEASETT
DBSCAN_FlhA1        TQQLVEHITRLAPKLIEEVVPKMVPIATFQKVLQLLLEESVHIRDIRTIVETLAEHAAAT
SP210_2_FlhA1       TQQLVEHVAKLAPKLIEEVVPKMVSIAVFQKVLQLLLDESVHIRDIRTIVETLAEHAGAT
SP51_3_FlhA1        TQQLVEHVAKLAPKLIEEVVPKMVSIAVFQKVLQLLLDESVHIRDIRTIVETLAEHAGAT
KACC21264_FlhA1     TQQLVEHVAKLAPKLIEEVVPKMVTIAVFQKVLQLLLDESVHIRDIRTIIETLAEHAAAT
Kf1_FlhA1           TQQLVEHVAKLAPKLIEEVVPKMVSIAVFQKVLQLLLDESVHIRDIRTIIETLAEHAGSV
GOD-11E_FhlA1       TQQLVEHVAKLAPKLIEEVVPKMVSIAVFQKVLQLLLDESVHIRDIRTIIETLAEHAGSV
bin23_FlhA1         TQQLVEHVAKLAPKLIEEVVPKMVSIAGFQKVLQLLLEESVHIRDIRTIIESLAEHAASV
CECT7646_FlhA1      TQQLVEHVAKLAPKLIEEVVPKMVSIAVFQKVLQLLLEESVHIRDIRTIVETLAEHAGGV
CFBP1192_FlhA1      TQQLVEHVAKLAPKLIEEVVPKMVSIAVFQKVLQLLLEESVHIRDIRTIVETLAEHAGGV
Leaf220_FlhA1       TQQLVEHVAKLAPKLIEEVVPKMVSIAVFQKVLQLLLEESVHIRDIRTIVETLAEHAGSV
ASV27_FlhA1         TQQLVEHVAKLAPKLIEEVVPKMVPIAVFQKVLQLLLDESVHIRDIRTIVETLAEYAGSI
GW821_FlhA1         TQQLVEHVAKLAPKLIEEVVPKMVPIAVFQKVLQLLLDESVHIRDIRTIVETLAEHAGSI
                    .  :. .:    . *   :  . :.   .  *:. ** * : :* : .*   *   .

CECT7646_SctV1      KDMLLLTEYIRADLGRYLAFEAA-GGGSV---VQAILMEPDAEQLIRQSIK----STPAG
CFBP1192_SctV1      KDMLLLTEYIRADLGRYLAFEAA-GGGSV---VQAILMEPDAEQLIRQSIK----STPAG
Kf1_SctV2           KDVLVLTEYVRGDLGRMIAWRAG-RGGNT---LGALFLQPELEQLIRQAIK----PTPTG
GOD-11E_SctV2       KDVLVLTEYVRADLGRMIAWRAG-RGGRQ---LRAVFLQVDLEQLIRQGIK----PTPTG
KACC21264_SctV2     KDVLVLTEYVRGDLGRMIAWRAG-GSARK---LHAVFLQVDLEQLIRQGIK----PTPTG
ASV27_FlhA3         KDASHLTEMVRNRLGHAICQSLL-GDANA---LHVMTLDPVVESQFMQSMQQARISRPEE
KACC21264_FlhA3     RDAVLLAEQVRQRLGHAICQSLLPGDAADPGALQVMTLDPAAESRLLGSFTA---PRPEA
GW821_FlhA2         KDPILLAAEVRCTLRRQIVAGLF-GQTQE---LKAFNLSGDLENLLLGALNQ---ARQTG
DBSCAN_FlhA1        TDPAELARRVRVALSPAIVQQIY-GPVRE---LSVIAIEPGLERLLVQAL-----GNPNG
SP210_2_FlhA1       QDPAELARRVRVALSPAIVQQIY-GPTRE---LSVIAIEPGLERLLVQAL-----GNPNG
SP51_3_FlhA1        QDPAELARRVRVALSPAIVQQIY-GPTRE---LSVIAIEPGLERLLVQAL-----GNPNG
KACC21264_FlhA1     QDPVELARRIRIALAPSIVQQIY-GQARE---LSVIAIEPQLERLLVQAL-----GSNNG
Kf1_FlhA1           QDPAELARRVRVALAPSIVQQIY-GQARE---LSVIAIEPNLERLLVQAL-----GSNNG
GOD-11E_FhlA1       QDPAELARRVRVALSPSIVQQIY-GQARE---LSVIAIEPQLERLLVQAL-----GSNNG
bin23_FlhA1         ADPAELARRVRIALAPAIVQQIY-GPVRE---LSVIAIEPGLERLLVQAL-----GHPNG
CECT7646_FlhA1      QDPAELARRVRVALAPAIVQQIY-GPIRE---LAVIAIEPGLERLLVQAL-----GNAAG
CFBP1192_FlhA1      QDPAELARRVRVALAPAIVQQIY-GPIRE---LAVIAIEPGLERLLVQAL-----GNAAG
Leaf220_FlhA1       QDPAELARRVRVALAPAIVQQIY-GPIRE---LAVIAIEPGLERLLVQAL-----GNAAG
ASV27_FlhA1         TDPAELARRVRVALAPAIVQQIY-GPTRE---LAVIAIEPGLERLLVQAL-----GGASG
GW821_FlhA1         SDPAELARRVRVALAPAIVQQIY-GPTRE---LAVIAIEPGLERLLVQAL-----GGAGG
                     *   *:  :*  *   :              :  .. :.   *  :  .:

CECT7646_SctV1      ----NYLAIAPEASAMLSDQIAQLAGDQARKGV--AVVTSMDIRRYVRKMIEVRLSWLRV
CFBP1192_SctV1      ----NYLAIAPEASAMLSDQIAQLAGDQARKGV--AVVTSMDIRRYVRKMIEVRLSWLRV
Kf1_SctV2           ----NFLTLPPEQVASTVDAILQISGAEPRTDL--AIVTAMDIRRYVRRMIQQRLPWLQV
GOD-11E_SctV2       ----NFLTLPPEQVTAAIEAIERIAGTTPCDDL--AVVTAMDVRRYVRRMIQQRLPWLAV
KACC21264_SctV2     ----NFLTLPPEQVAGAIEAIQRVSGEQPRDDL--AIVTAMDIRRYVRRMIQQRLPWLNV
ASV27_FlhA3         GSQLPAFVLEPRLAEQFITRLLQTAERMMKSNLLPVLLCSPELRRHVRALSERVLPHLRV
KACC21264_FlhA3     ---QAQFAIEPQLAEQLLLRLVQQAERMMKNNLLPVLLCAPELRRHLRAMSERVMPHLRV
GW821_FlhA2         KVALDNYPIDPHLLAQMQLNMPAARTQMKQQGAVPLLLVMPQIRPLLARYARLFAPGLHV
DBSCAN_FlhA1        ------AALDPGVADILSRTAAEVALQQEEQGLPACLLVPDAIRGAIARLVRRVAPRLQV
SP210_2_FlhA1       ------PALDPGVADILTRTAAEVAMKQEEMGLPACLLVPDQIRTSIARLVRRVAPRLQV
SP51_3_FlhA1        ------PALDPGVADILTRTAAEVAMKQEEMGLPACLLVPDQIRTSIARLVRRVAPRLQV
KACC21264_FlhA1     ------PSLDPGVADMLTRSAAEVALKQEEIGLPACLLVPDQIRSAMARLVRRVAPRLQV
Kf1_FlhA1           ------PSLDPGVAEMLTKSAAEVALRQEEIGLPACLLVPDQIRSAIARLVRRVAPRLQV
GOD-11E_FhlA1       ------PSLDPGVADMLTRSAAEVALKQEEIGLPACLLVPDQIRSAMARLVRRVAPRLQV
bin23_FlhA1         ------PSLDPGVADILTRTAAEVALKQEEMGLPACLLVPDAIRGAVARLVRRVAPRLQV
CECT7646_FlhA1      ------PALDPGVADVLTRSAAEVALKQEELGLPACLLVPDVIRSAIARLVRRVAPRLQV
CFBP1192_FlhA1      ------PALDPGVADVLTRSAAEVALKQEELGLPACLLVPDVIRSAIARLVRRVAPRLQV
Leaf220_FlhA1       ------PALDPGVADVLTRSAAEVALKQEELGLPACLLVPDVIRSAIARLVRRVAPRLQV
ASV27_FlhA1         ------PALDPGVADVLTRSAAEVALKQEEMGLPACLLVPDVIRSAVARLVRRVAPRLQV
GW821_FlhA1         ------PALDPGVADVLTRSAAEVALKQEELGLPACLLVPDIIRSAIARLVRRVAPRLQV
                            : *                    .    ::    :*  :        . * *
```

**Supplementary Data S1**

```
CECT7646_SctV1      YSFQELGNTIELRPIGRVV-----V-----------
CFBP1192_SctV1      YSFQELGNTIELRPIGRVV-----V-----------
Kf1_SctV2           YSFHELGDYLQLVSVGEAR-----LG----------
GOD-11E_SctV2       YSFHELADHLELVSAGEAR-----TG----------
KACC21264_SctV2     FSFHELGEHLELVSAGEARLAEPPRGGGGMGAPGMR
ASV27_FlhA3         LSMTEIPHSVELK-SYAVV--------SL-------
KACC21264_FlhA3     LSMAEVPQNVELK-SWAVV------GL---------
GW821_FlhA2         LSYNEIPEQRDIN-I---------VGSLG-------
DBSCAN_FlhA1        LAHSEIPETHTIR-IGPIL-----RGAAA-------
SP210_2_FlhA1       LAHSEIPETHSIR-IGPIL-----RGASA-------
SP51_3_FlhA1        LAHSEIPETHSIR-IGPIL-----RGASA-------
KACC21264_FlhA1     LAHSEIPETHSIR-IGPIL-----RGANA-------
Kf1_FlhA1           LAHSEIPETHSIR-IGPIL-----RGATA-------
GOD-11E_FhlA1       LAHSEIPETHSIR-IGPIL-----RGATA-------
bin23_FlhA1         LAHSEIPETHTIR-IGPIL-----QGASA-------
CECT7646_FlhA1      LAHSEIPETHTIR-IGPIL-----RGA---------
CFBP1192_FlhA1      LAHSEIPETHTIR-IGPIL-----RGA---------
Leaf220_FlhA1       LAHSEIPETHTIR-IGPIL-----RGA---------
ASV27_FlhA1         LAHSEIPETHTIR-IGPIL-----RGASA-------
GW821_FlhA1         LAHSEIPETHTIR-IGPIL-----RGASA-------
                     :  *:     :
```

[https://www.ebi.ac.uk/jdispatcher/msa/muscle?stype=protein](https://www.ebi.ac.uk/jdispatcher/msa/muscle?stype=protein)
