## Supplementary Data S2 for "Identification of protein secretion systems and type III effectors in wood-associated bacteria of the genus *Xylophilus*"

**HrpX-regulated protein family 1**

```
>QHI96638.1 hypothetical protein GT347_00670 [Xylophilus rhododendri]
MNLPSTSATAPRLHHPYEFDPLAPAGTASQAAGTVATQPPLIDVPTASGEACFADKRQQFARLVDDIDTL
TQQDAPVLRRRMLDFLRFGGHDIAPVMLQPTQMHLDAACRGYALPDPLLVESVGMRALVCWLPASTGWEA
GKRGYFKQALMAAFDALEAAQAVRRQPPGLDIADRKAVEGHLAELVADPCADKRGHKRAALLSILGRYGY
EGGVRPNAWTWKRLQDGGPDAQDEALGMPQLLAAIESMPAHPELHASLQQALLQAVVEQGRQWKAAQALG
VQAQPEPEPEPDAARVDRPVNESAAPDAARTPLARFEASWRWIEAEVPPSTGIPSRHVFRPAGSKASAGD
VLLYRGGSGTRMVVQSVRQSRFLSQDINTQGLALLRNADGTAAVTLDRQPVTDPRTLDLLLRAVETFRGD
PASGPALLLQESLELLLGAAEPDIAQKARAAQFALAGLMSIDLPAGEPMLADGALAWRDPEALLVVLTLA
GSSADLRCWVAAFQVGLRNPQDVLAQARVDIAIRLALAGQPDPAQLRFYRRAGVDERHAIVAELLAFTRE
KSLFCLGTLLWRTAPQDARDGRDDSVGTLEYASPDGFARVTDMLCELRQAFEEDPQFIAICEAFVAEFAM
GCTDRAAIGLAGLGLLLNWHKVRDDQTRALQALRSLLNFHRMAEAAVSMLTASGSSSEPAQAALRLMIAA
RSRYLLLPDHGPRYLRHLDEEGTHAATMQRMDAALRFIMEASPGPVLSTPLIQASPYGADYIQRNLGHVE
PAQLEQRGDEIERLLTPYAYLRLLPGLACTPLTVPVLKRLLLEVVAPDVEPPTPMQELLRRMGAALTQQD
GWPAEAQAEQMRRGLIGELIDAIARQEPAFAALVARRLTVGDSLQAQCAALAEELSG

>QHI96636.1 hypothetical protein GT347_00660 [Xylophilus rhododendri]
MTYPSVSVTTSSHAGYLPDVLHSQATPAADPPSRTAADQPLPDIAATGGAYFSGAREQLDTLMSGLDAAQ
VEDAPALRRQLLDFLRHRQHTVQESFLGSSEVWLDAACFGEPLPHPREVEAMAMGSLLCLLPRCGDWEVR
KRAYLKQALGRAFEALEEAGGVAAAAEPSPVEPAQMQGPKAEVVEASMPARFDAVWHSIEGWLQAGMGAL
SPLAGQLPGEVPAAGAVVCHRDASGIRQTIQAHGDHAGTDPAAASRGIALCRNPDGSTAVWLDGQAVADP
QVLDLLLLAAQTFLGHHLPADSQGRSTPATLVERACGLLRNPGPDIGLRTRAAQFALAGLLNVEVPAVPP
EGAGGAFWPGFPRELLALLPGVSQEKRLASWLSTWIDGHYRPEDVLAQARLDMVMGTLRSQAPEADLLRH
HRRWSDEERDTVIPRLCDQAKAHGLFGLWQLWARCQPPDEPDAFHALPEYRTPDGRERLTAMLCELHAAF
EADEEFARACNRFAAECATGSSDRALIGLNGLAPMLNWWLARGNPAKELLALRSLLHFFRLMQEGERIGD
GESFGGVLHTRLELMRLSRTKHLPIPDSGPCVPRSELPAAEQQAEIERLNPALQKAVADSARPVLLSPAI
QASYCGRAFIERYVGLEPAETLDERCAALETMMAPYAYVGVLPGLERVALTAPVLKRLLQQVLASPEQGV
SQSPVLRRLYRLAAALGRQSGLRPSECADRTKRVVAGELLDAIFRLEPAFAAAVAPRLKLQDPLKVQAAA
LTELLEQDVEAQARSAASRAAAS
```

```
QHI96638.1      MNLPSTSATAPRLHHPYEFDPLAPAGT-ASQAAGTVATQPPLIDVPTASGEACFADKRQQ
QHI96636.1      MTYPSVSVTTSS-HAGYLPDVLHSQATPAADPPSRTAADQPLPDI-AATGGAYFSGAREQ
                *. **.*.*:.  *  *  * *  . .* *:::... .*:: ** *: :*:* * *:. *:*

QHI96638.1      FARLVDDIDTLTQQDAPVLRRRMLDFLRFGGHDIAPVMLQPTQMHLDAACRGYALPDPLL
QHI96636.1      LDTLMSGLDAAQVEDAPALRRQLLDFLRHRQHTVQESFLGSSEVWLDAACFGEPLPHPRE
                :  *:..:*:   :***.***.:*****.  * :   :* .:::.***** * .** *

QHI96638.1      VESVGMRALVCWLPASTGWEAGKRGYFKQALMAAFDALEAAQAVRRQPPGLDIADRKAVE
QHI96636.1      VEAMAMGSLLCLLPRCGDWEVRKRAYLKQALGRAFEALEEA-------------------
                **::.* :*:* **  . .**. **.*:****  **:*** *

QHI96638.1      GHLAELVADPCADKRGHKRAALLSILGRYGYEGGVRPNAWTWKRLQDGGPDAQDEALGMP
QHI96636.1      --------------------------------GGVA------------------------
                                                ***

QHI96638.1      QLLAAIESMPAHPELHASLQQALLQAVVEQGRQWKAAQALGVQAQPEPEPEPDAARVDRP
QHI96636.1      ---AAAEPSPVEP-----------------------AQMQGPKAE---------------
                   ** *. *. *                       **  * :*:

QHI96638.1      VNESAAPDAARTPLARFEASWRWIEAEVPPSTGIPSRHVFRPAGSKASAGDVLLYRGGSG
QHI96636.1      VVEASMP-------ARFDAVWHSIEGWLQAGMGALSPLAGQLPGEVPAAGAVVCHRDASG
                * *:: *       ***:* *. **. : .. *  *   . . .*. .:** *: :*..**

QHI96638.1      TRMVVQSVRQSRFLSQDINTQGLALLRNADGTAAVTLDRQPVTDPRTLDLLLRAVETFRG
QHI96636.1      IRQTIQAHGDHAGTDPAAASRGIALCRNPDGSTAVWLDGQAVADPQVLDLLLLAAQTFLG
                 * .:*:  :     .    :.*:** **.**::** ** *.*:**..***** *.:** *

QHI96638.1      D--PA-----SGPALLLQESLELLLGAAEPDIAQKARAAQFALAGLMSIDLPAGEPMLAD
QHI96636.1      HHLPADSQGRSTPATLVERACGLLRNPG-PDIGLRTRAAQFALAGLLNVEVPAVPPEGAG
                   **     * ** *:: :  ** ... ***. .:*********:.:::**  *  *.
```

### Supplementary Data S2

```
QHI96638.1      GALAWRD-PEALLVVLTLAGSSADLRCWVAAFQVGLRNPQDVLAQARVDIAIRLALAGQP
QHI96636.1      GAF-WPGFPRELLALLPGVSQEKRLASWLSTWIDGHYRPEDVLAQARLDMVMGTLRSQAP
                **: * . *  **.:*. ....  * .*::::  *  .*:*******:*:.:     :  *

QHI96638.1      DPAQLRFYRRAGVDERHAIVAELLAFTREKSLFCLGTLLWRTAPQDARDGRDDSVGTLEY
QHI96636.1      EADLLRHHRRWSDEERDTVIPRLCDQAKAHGLFGLWQLWARCQPPDEPDAFH---ALPEY
                :.  **.:** . :** :::. *   :. :.** *  *  *  * *  *.     .  **

QHI96638.1      ASPDGFARVTDMLCELRQAFEEDPQFIAICEAFVAEFAMGCTDRAAIGLAGLGLLLNWHK
QHI96636.1      RTPDGRERLTAMLCELHAAFEADEEFARACNRFAAECATGSSDRALIGLNGLAPMLNWWL
                 :***  *:* *****. *** * :*   *: *.** * *.:*** *** **. :***.

QHI96638.1      VRDDQTRALQALRSLLNFHRMAEAAVSMLTASGSSSEPAQAALRLMIAARSRYLLLPDHG
QHI96636.1      ARGNPAKELLALRSLLHFFRLMQEG-ERIGDGESFGGVLHTRLELMRLSRTKHLPIPDSG
                .*.: :. * ******:*.*: : . . :  . * .   :: * **  :*:.:* :** *

QHI96638.1      PRYLRHLDEEGTHAATMQRMDAALRFIMEASPGPVLSTPLIQASPYGADYIQRNLGHVEP
QHI96636.1      PCVPRSELPAAEQQAEIERLNPALQKAVADSARPVLLSPAIQASYCGRAFIERYVG-LEP
                *   *     . : * ::*::.**.  :  *. *** :* ****  *  :*:* :* :**

QHI96638.1      AQ-LEQRGDEIERLLTPYAYLRLLPGLACTPLTVPVLKRLLLEVVAPDVE--PPTPMQEL
QHI96636.1      AETLDERCAALETMMAPYAYVGVLPGLERVALTAPVLKRLLQQVLASPEQGVSQSPVLRR
                *: *::*   :* :::****: :****  ..**.******* :*:*.  :  . :*:

QHI96638.1      LRRMGAALTQQDGWPAEAQAEQMRRGLIGELIDAIARQEPAFAALVARRLTVGDSLQAQC
QHI96636.1      LYRLAAALGRQSGLRPSECADRTKRVVAGELLDAIFRLEPAFAAAVAPRLKLQDPLKVQA
                * *:.*** .*.*  ..  *:. .* : ***:*** * ****** ** **.: *.*:.*.

QHI96638.1      AALAEELSG----------------
QHI96636.1      AALTELLEQDVEAQARSAASRAAAS
                ***:* *.
```

### Supplementary Data S2

**HrpX-regulated protein family 2**

>QHI96917.1 hypothetical protein GT347_02290 [Xylophilus rhododendri]
MRFDPAAGTAFPNTSTTSTVSTISTMANVALAVLQRRQRLGHATRLDHVPIEGVNQAQLAQFQPMSQEMK
DLLDLEPEASANASAPEQDTVFTEPVSRRRTRRAGPELQDRLDASIRSMKTAFRRVLGLPGTFFVPGGTS
GNDAFETRATQLATSYAKAIQPHLTTLDWEAIHSNARSSLQLALEGWIEFYAHAYATMDHPYEWNGAYIL
DWARTEAAQIDAEADSPAFANRIRASFLAGKASPIFNSPDAGLLTPLDSFASVSDAQAFAACGRPARLAE
ITRDGQVRSRTITPSAPDCQGDGLTHHAYVVGRQLAHQEPGSIFFSLSSLAVTPRNEQLPITATLNIRQC
GDETTRREIQSAQQVDREPPPCPPYVPPLQAWRKSAQAELVQRYQAQASQRPSYARHRYAP

>QHI96926.1 hypothetical protein GT347_02335 [Xylophilus rhododendri]
MVFDLASNSSHGATPTAATFASLAAGLLWRSASQTSMETASRPFSDARGPKQELSDEMTALLGLKSSAAI
KDDAIFGAGQPDQKPRISQNDIKPEARRTSRTKRASPYLQNRLDTALRYALAGFDQQFGRHNPNLPAAPD
SFSQLAFDKGLEYGGTIRGTLNSAEYTLMHSNENSQLRQVLEDWVTVYAYAHATAGDLYRWDYYKIIGWA
RREVAQIDAQADDDVGEKRRLEAYHAGKNSEIAISRTRGIIGALWAYPNEADMNRFAPCGKPVYVFDWID
KNALKYQFIVRLPDPDKCLGDASTDEAYMVSRRLWRGSQNEIFRKEDAVHMPRGVILRVSSNEVAQNCGP
DNILIRKVSMFNPFYKIPLCPPYVPPYKHWRDSMKSSLVTEYERKHKRGLSEQDVFSANQRQPTVQPQQR
VDRLPRPSA

>QHI96928.1 hypothetical protein GT347_02345 [Xylophilus rhododendri]
MGFNIVSNTTNGNGSNIGAVANLALGLMGRYPRRVTLQNNSRPSETRVDEYRELNAEMATLLGLTADDVG
RDGAIQHSAIFSQDADIEPEKPLARKKRNAPLDFKRRLDVTLHNALAGFRTACRIPGETASVPQDPFEKR
QFQNGESYGGGTCSFLNTDEINSLRNDKQSPLRQILEGWVRVQAHSDATGSGVHTWKSANILAWARDEVK
QVISQADKPDREERLRDAFQRGRSSPLAFERLYSATSPLDCYPDLRDLHEFAECGVATRLIEWIGHGGGE
NHPIFSISLVYPDPAQCKGDATTHMAYITGRRIGYLPDWIFDNPKVIVNVPRGVTAWLADNVKARNCGPD
NILRKVISRWNPFAGDVPACPPFVTRHAQWRAAIIKGILDNIDYWQRKRQLEEDQVPKVQWQQVSPPART
NLQRQFDRQSQEFREQQWRLQQQAATARQPG

>QHI96931.1 hypothetical protein GT347_02360 [Xylophilus rhododendri]
MGFPLTAGNAMATKLVGMPRFPFRGTQDSAATAEKRILMSVADSAHVNNLENIGAISSEMVDLLQLRPGA
IAGAGEVHYDDSFKMKRALEKQPPSIARTRRLTDDMQILLDNTINNAVGAYLVEMGQPGFFSDNIKNPAT
KAAIDLGQNIARAIKPHLNEYERQMLRSSSAFSAAFKNWVEVYAYRLVFGGVGYEEDGKNIVEWVRQEAA
QLDSQADYPAIEESLRRGYKGGLQSKLVGKYLHQIPLSVYPNVEDMENYAPCNTDVLLYAGNKRSGQHSV
IHARVGPERCKGNVYTDASYRVARFWQHTAKDHVRFSLNEVQIVPRGVVLPTGEDTWSRNCGPDNILTRR
GSFWDWFNHGAPACPPYVSLFEKWRETLKNGIVNNFKATLPNGMIPQSPPRPAPPKLPNRG

>QHI96929.1 hypothetical protein GT347_02350 [Xylophilus rhododendri]
MGFDTTSFVGGSMPAASNISALAAVAQGAAAHFLNKSKSTELKASHTSFSKSELSPQEISHDLQELLGLP
TPATPENPQAVFHGAFSASKMPATLNTSPNESIKPASIPARHSIGKRAASTTLQQRLKIAIPVTIASFRH
TMGLPNSIIHPNNNLLQVTSILNGRRIGLAIRPLLTHIEMQTLQSQTTSAFQNSLEAWVEYLAYRMAAIN
QRFEDNQEEILKWVREETAQVISQADNFIREEKIRTAYLAGKNSLLFGNSDPAQLHPAYAYATMQDMNNY
APCDFIAFVWYKKNDKQYGTMLARTLHECSGNTETHHAYSVGRQQKNLGQEAVFSSDYLNDIPRGERLRL
EGNLDARNCGPGNFLYKQVPTFNWFKTTVPPLCPPYVSPHTRRRTELIAQLDQDFDNASNKRSLDPRILS
TMTLGVPAQTEPNQQVFGPLQRPFIDQPSFDKQRQQFDGQAINQPGGHGRNG

>QHI96925.1 hypothetical protein GT347_02330 [Xylophilus rhododendri]
MRFSLANAIPSTTARVPERFMLADMQHSNPPETRQPLAARVFDDSLSHIGAVSQQMKDLLDFLPGEIAGS
ADVHYGQGFTTATADPVRREDVLLDDPFRDVPEEQSEVVARDKRLEDDIQIKLDNTVNNAVGAYLLEIGQ
VGFISDNPEGPVAKPAIALGRQIARAIQPHLSEIERKLPRLSPFSQALKGWVEVYAYRLALGGLEFKGGG
EDIVEWARKEAAQTDAHADHPERERSLQAGYRGGLKSKRGGTPRRQAPLPAYPDREDMQNYAPCGQDVLL
HEFNRQMKKYATISVRPHPDNCRGNQYTHIAYGIGRIFAHTPWQLTKFTVDTVRNVPRDVQLPTGRDTWR
RNCGPDNIVTKNVSFWDWSSHGVPVCPPYVSRLAKWREQLKEAMETNFQTMV

>QHI96932.1 hypothetical protein GT347_02365 [Xylophilus rhododendri]
MLSTAKATSGAPNPTGSLFNKPLDPIGTISPDLRSFGGLGPVAIEILELMRAKADAPQPLKHRPVPVLDQ
NRLPPQEIGWETAELLGMHSRSGVEDAGFDQPRAMPRAKRVATASKEDLETAARYSVQGFRHEVGLPDLS
VPAPETPLSTQALRNGRDFAQQIKPLLNDFEAHIVRSQPASTFVRALEGWAEVSGYKMVASGFPFTQTPA
SILNMVRSEAAQMDAEADNPEEEGWRRSGFRGGMSSVFLHFHSKPSMSGVYSYPSNEDVMRYAPCGATVV
LFGPVQNRANFGPKMVQPKPGQCHGNWFTHKAYMVGRRAKHVIDPIFSLDELQAAPFNFQGLLSPNVKGR
NCGSGNVLTKEFQWWHLRKSAPECPPYESPHKKQRNQLKAAMRESLEKKHATRQFDDMFEMPPEMSQNSF
QPRRPGF

>QHI96930.1 hypothetical protein GT347_02355 [Xylophilus rhododendri]
MSFPISATNILTDNLDRARRFLFPGMQDSSAAAKDKIPTSVADTVYKDSLKDIGEISPKMIDLLGLKRDD
VAKSGEVHYGEPFRAKRAALSDPNAPVPKSVFRASAGLQPAAAPDIRADQAEVPTASEGLTEEMQIRLDG
TVSNAVAAFLEETGQPGVDLDDSDEPAAKAAAALGRQIARAVKPHLTEYELKLLGSSSPFTDAFKGWVEV
YAYRLTFGNLDYTEDGKKLVAWARQEAAQVEAQSDNPRRQNSLAYGYEGGLESERGGTRRRPAPLDAYPN
KQDMESYAPCGKFVRLYQFNPHIGDYATVYTRVDPEDCRGNKDTDIAYQLGRFFTHTSHLNVKFHITDME
NLPYGVEVPTGDNAWSRNCGPDNILTADNNRWEWFGKGAPECPPYVSNLAKWRDHLKSFMSTDFWAALPD
GNLPKALPKRLLTSSALRTPSRPSNSTRA

### Supplementary Data S2

```
>QHI96924.1 hypothetical protein GT347_02325 [Xylophilus rhododendri]
MHAYAQATGGDLLTWEPQKILSWVRKELAQVRAQADDDQRETRIREAFDAGKNSQIARSRTHGIVSPIYA
YPNMEDVNKFAPCGIPVLRFQWVDPDVPSYSLIFGPPDPALCRGNAETDIAYKLARQIAHGPRNQLFKPD
NAKYLPRNMAVQVSRTEVSQNCGPDNLVIKKIPALSPLIRFDLCPPYVSPYHRWRESMKNSLVTDYEAKR
KREFFDPQPQQQFSPPQRPFVGQPGFDNQRKQFDGQAFNQAGRQQQDG
```

```
QHI96932.1      MLSTAKATSGAPNPTGSLFNKPLDPIGTISPDLRSFGGLGPVAIEILELMRAKAD-----
QHI96917.1      ----------------MRFDPAAGTAFPNTSTTSTVSTISTMANVALAVLQRRQRLGHAT
QHI96928.1      ----------------MGFNIVSNTTNGNGSNIGAVANLAL------GLMGRYPR-----
QHI96930.1      ----------------MSFPISA----TNILTDNLDRARRFLFPGMQDSSAAAKD-----
QHI96925.1      ----------------MRFSLAN-----AIPSTTARVPERFMLADMQHSNPPETR-----
QHI96931.1      ----------------MGFPLTA----GNAMATKLVGMPRFPFRGTQDSAATAEK-----
QHI96929.1      ----------------MGFD-TTSFVGGSMPAASNISALAAVAQGAAAHFLNKSK-----
QHI96926.1      ----------------MVFDLASNSSHGATPTAATFASLAA------GLLWRSAS-----
QHI96924.1      ------------------------------------------------------------

QHI96932.1      -APQPLKHRPVPVLDQNRLPPQEIGWETAELLGMHSRSGVEDAGFD--------------
QHI96917.1      RLDHVPIEGV---NQAQLAQFQPMSQEMKDLLDLEPEASANASAPEQDTVFT--------
QHI96928.1      RVTLQNNSRP---SETRVDEYRELNAEMATLLGLTADDVGRDGAIQHSAIFSQD------
QHI96930.1      KIPTSVADTV---YKDSLKDIGEISPKMIDLLGLKRDDVAKSGEVHYGEPFRAKRAALSD
QHI96925.1      ---QPLAARV---FDDSLSHIGAVSQQMKDLLDFLPGEIAGSADVHYGQGFTTATADPVR
QHI96931.1      RILMSVADSA---HVNNLENIGAISSEMVDLLQLRPGAIAGAGEVHYDDSFKMK------
QHI96929.1      STELKASHTS---FSKSELSPQEISHDLQELLGLPTPATPENPQAVFHGAFSASKMPATL
QHI96926.1      QTSMETASRP---FSDARGPKQELSDEMTALLGLKSSAAIKDDAIFGAGQPDQKPRISQ-
QHI96924.1      ------------------------------------------------------------

QHI96932.1      ----------------QPRAMPRAKR-----------VATASKEDLETAARYSVQGFRHE
QHI96917.1      ----------------EPVSRRRTRR-----------AGPELQDRLDASIRSMKTAFRRV
QHI96928.1      -----------ADIEPEKPLARKKRN-----------APLDFKRRLDVTLHNALAGFRTA
QHI96930.1      PNAPVPKSVFRASAGLQPAAAPDIRADQAEVPTASEGLTEEMQIRLDGTVSNAVAAFLEE
QHI96925.1      REDVLLDDPFRDVPEEQSEVVARDKR-----------LEDDIQIKLDNTVNNAVGAYLLE
QHI96931.1      -----------RALEKQPPSIARTRR-----------LTDDMQILLDNTINNAVGAYLVE
QHI96929.1      NTSPNESI---KPASIPARHSIGKRA-----------ASTTLQQRLKIAIPVTIASFRHT
QHI96926.1      -----------NDIKPEARRTSRTKR-----------ASPYLQNRLDTALRYALAGFDQQ
QHI96924.1      ------------------------------------------------------------

QHI96932.1      VGLPDLSV----PAPETPLSTQALRNGRDFAQQIKPLLNDFEAHIVRSQPASTFVRALEG
QHI96917.1      LGLPGTFFVPGGTSGNDAFETRATQLATSYAKAIQPHLTTLDWEAIHSNARSSLQLALEG
QHI96928.1      CRIPGETA----SVPQDPFEKRQFQNGESYGGGTCSFLNTDEINSLRNDKQSPLRQILEG
QHI96930.1      TGQPGVDL----DDSDEPAAKAAAALGRQIARAVKPHLTEYELKLLGS--SSPFTDAFKG
QHI96925.1      IGQVGFIS----DNPEGPVAKPAIALGRQIARAIQPHLSEIERKLPR---LSPFSQALKG
QHI96931.1      MGQPGFFS----DNIKNPATKAAIDLGQNIARAIKPHLNEYERQMLRS--SSAFSAAFKN
QHI96929.1      MGLPNSII----HPNNNLLQVTSILNGRRIGLAIRPLLTHIEMQTLQSQTTSAFQNSLEA
QHI96926.1      FGRHNPNL----PAAPDSFSQLAFDKGLEYGGTIRGTLNSAEYTLMHSNENSQLRQVLED
QHI96924.1      ------------------------------------------------------------

QHI96932.1      WAEVSGYKMVASGFPFTQTPASILNMVRSEAAQMDAEADNPEEEGWRRSGFRGGMSSVFL
QHI96917.1      WIEFYAHAYATMDHPYEWNGAYILDWARTEAAQIDAEADSPAFANRIRASFLAGKASPIF
QHI96928.1      WVRVQAHSDATGSGVHTWKSANILAWARDEVKQVISQADKPDREERLRDAFQRGRSSPLA
QHI96930.1      WVEVYAYRLTFGNLDYTEDGKKLVAWARQEAAQVEAQSDNPRRQNSLAYGYEGGLESERG
QHI96925.1      WVEVYAYRLALGGLEFKGGGEDIVEWARKEAAQTDAHADHPERERSLQAGYRGGLKSKRG
QHI96931.1      WVEVYAYRLVFGGVGYEEDGKNIVEWVRQEAAQLDSQADYPAIEESLRRGYKGGLQSKLV
QHI96929.1      WVEYLAYRMAAINQRFEDNQEEILKWVREETAQVISQADNFIREEKIRTAYLAGKNSLLF
QHI96926.1      WVTVYAYAHATAGDLYRWDYYKIIGWARREVAQIDAQADDDVGEKRRLEAYHAGKNSEIA
QHI96924.1      ---MHAYAQATGGDLLTWEPQKILSWVRKELAQVRAQADDDQRETRIREAFDAGKNSQIA
                     .:  .  .         ::  .* *  *  : :*          .:  *  *
```

### Supplementary Data S2

```
QHI96932.1      HFHSKPSMSGVYSYPSNEDVMRYAPCGATVVLFGPVQNRAN-----FGPKMVQPKPGQCH
QHI96917.1      NSPDAGLLTPLDSFASVSDAQAFAACGRPARLAEI--TRDGQ----VRSRTITPSAPDCQ
QHI96928.1      FERLYSATSPLDCYPDLRDLHEFAECGVATRLIEWIGHGGGENHPIFSISLVYPDPAQCK
QHI96930.1      GTRRRPA--PLDAYPNKQDMESYAPCGKFVRLYQF-NPHIGD----YATVYTRVDPEDCR
QHI96925.1      GTPRRQA--PLPAYPDREDMQNYAPCGQDVLLHEF-NRQMKK----YATISVRPHPDNCR
QHI96931.1      GKYLHQI--PLSVYPNVEDMENYAPCNTDVLLYAG-NKRSGQ----HSVIHARVGPERCK
QHI96929.1      GNSDPAQLHPAYAYATMQDMNNYAPC--DFIAFVWYKKNDKQ----YGTMLAR-TLHECS
QHI96926.1      ISRTRGIIGALWAYPNEADMNRFAPCGKPVYVFDWIDKNALK----YQFIVRLPDPDKCL
QHI96924.1      RSRTHGIVSPIYAYPNMEDVNKFAPCGIPVLRFQWVDPDVPS----YSLIFGPPDPALCR
                             :.   *   :* *                                *

QHI96932.1      GNWFTHKAYMVGRRAKHVIDPI--FSLDELQAAPFNFQGLLSPNVKGRNCGSGNVLTKEF
QHI96917.1      GDGLTHHAYVVGRQLAHQEPGSIFFSLSSLAVTPRNEQLPITATLNIRQCG-DETTRREI
QHI96928.1      GDATTHMAYITGRRIGYLPDWIF-DNPKVIVNVPRGVTAWLADNVKARNCGPDNILRKVI
QHI96930.1      GNKDTDIAYQLGRFFTHTSHLNVKFHITDMENLPYGVEVPTGDNAWSRNCGPDNILTADN
QHI96925.1      GNQYTHIAYGIGRIFAHTPWQLTKFTVDTVRNVPRDVQLPTGRDTWRRNCGPDNIVTKNV
QHI96931.1      GNVYTDASYRVARFWQHTAKDHVRFSLNEVQIVPRGVVLPTGEDTWSRNCGPDNILTRRG
QHI96929.1      GNTETHHAYSVGRQQKNLGQEAV-FSSDYLNDIPRGERLRLEGNLDARNCGPGNFLYKQV
QHI96926.1      GDASTDEAYMVSRRLWRGSQNEI-FRKEDAVHMPRGVILRVSSNEVAQNCGPDNILIRKV
QHI96924.1      GNAETDIAYKLARQIAHGPRNQL-FKPDNAKYLPRNMAVQVSRTEVSQNCGPDNLVIKKI
                *:  *  :*  .*                    * .           .:** .:

QHI96932.1      QWWH-LRKSAPE-CPPYESPHKKQRNQLKAAMRESLEKKHATRQFD--------------
QHI96917.1      QSAQQVDREPPP-CPPYVPPLQAWRKSAQAELVQRYQAQAS-------------------
QHI96928.1      SRWNPFAGDVPA-CPPFVTRHAQWRAAIIKGILDNIDYWQRKRQLE--------------
QHI96930.1      NRWEWFGKGAPE-CPPYVSNLAKWRDHLKSFMSTDFWAALPDGNLP--------------
QHI96925.1      SFWDWSSHGVPV-CPPYVSRLAKWREQLKEAMETNFQTMV--------------------
QHI96931.1      SFWDWFNHGAPA-CPPYVSLFEKWRETLKNGIVNNFKATLPNGMIP--------------
QHI96929.1      PTFNWFKTTVPPLCPPYVSPHTRRRTELIAQLDQDFDNASNKRSLDPRILSTMTLGVPAQ
QHI96926.1      SMFNPFYK-IPL-CPPYVPPYKHWRDSMKSSLVTEYERKHKRGLSE--------------
QHI96924.1      PALSPLIR-FDL-CPPYVSPYHRWRESMKNSLVTDYEAKRKREFFDPQ------------
                             ***: .    .*      :

QHI96932.1      -----DMFEMP--PEMSQNSFQPRRPGF-------------------------
QHI96917.1      ------------QRPSYARHRYAP-----------------------------
QHI96928.1      ----EDQVPKVQWQQVSPPARTNLQRQFDRQSQEFREQQWRLQQQAATARQPG
QHI96930.1      -------KALPKRLLTSSALRTPSRPSNSTRA---------------------
QHI96925.1      -----------------------------------------------------
QHI96931.1      ------------QSPPRPAPPKLPNR---------------------------G
QHI96929.1      TEPNQQVFGPLQRPFIDQPSFDKQRQQFDGQAIN----------QPGGHGRNG
QHI96926.1      ----QDVFSANQRQPTVQPQQRVDR----------------------LPRPSA
QHI96924.1      ---PQQQFSPPQRPFVGQPGFDNQRKQFDGQAFN----------QAGRQQQDG
```

### Supplementary Data S2

**HrpX-regulated protein family 3**

>QHI97260.1 hypothetical protein GT347_04255 [Xylophilus rhododendri]
MHLYPISTPSNNSSAPSVVSTLALLAQGVIQRYLPSHVNPTPRVVENDQLPSQGISHEMAGLLALDETHA
KTQSRSKRESSASDYRRKETTEPDYDANQGGRAFDKPSSIPRTNALPSDPVPRHAYYQRALNKAPQKPEQ
SDTVTDYYGRPVNPPPGAYGFNSETHPSGRGRKQFLGYDMTGKPIYRADSRYLVGWDEQRKPIYSDDVRP
QPSIDPSLIGNIPSTPAYGKGFEFADTNWWDYTTTTRQPRKKAPAKKPSHLNQYDADGNILPSKATRREG
KNQYIDGGTGTVLASSEPMLHNFIHDPAPSDSPKKQHRFFI

>QHI97411.1 hypothetical protein GT347_05075 [Xylophilus rhododendri]
MTAEFNPSSSANNSAAHSSISALTLLTQSLLRRNQKQPQDTLQTDSIIKEPNLPPREISEEMNKLLNLKS
TEASHESLHPPAEVPANSESDKPRHGRSRRELRAMQDRNGRHHYVTAQVEAQHGGQFVEWATGLYHYYHN
GVLQGQTSYIVGLDGSIYQIFEPQHAGPAPASAYSQGQTGIRTTQAPPQITTSRNPKLPYWGPNQSDLPG
VSTRKPGQAGNNNFDGNAATRAGVFQKSSLAARHFSTKRPAPLYHSKYDENGRPVDASQEYTGVTDYYGR
TVKTHVVGYDMNGKEIFAHLGRRKLIGMLPNKTAVYGDYERVLIGWDENRRPMYMGDNRPQGTIDPQYLG
RTASTPAYGKGFEFADTNWWEHNYTTTTRGPRINIPTSRPSYLNQYDKYGHVTRDPETEKTRGRGRSRAG
QHQPAGPGNPAPIQGQQPGHRPPDNRAPHYGENSDPGHKYRNPAGGHLPQRHGSSNQLARWQVARPKYPQ
YFVFEDIIYKGVDQMFASVRSKRQIGVRFTELAAPPIPTGNQSIDIQAWTDYAAQNNIYQMQQRQFSSTI
ESEKPLMDTIKTLEREMFEAEQARMENAQQKRESVDETAVAKLPVAYFTSETMDALLEEVLPRYRDAYGN
ALTEVQAEESFHNKIAEDIKTRIFQYFRNPESELEKKIIRFIKACAGFKTIFGKNASFVFTPTLQKGLVG
EYPRNISLELRYALADKLLNIQYNPIYIAAGPPDHPYIHLSNAFVQKIETELYDTQTIVNIYYDPAHADA
LTRASEGLEKTAKTRIADEMIARRTPGFPLVRTSKAFFTPESGKYSPVSINTLEEYEQVVIAYDLEPGVS
IRDERAPLEVLARMPFQQAGQELRKIFLPLKSRLDRVFVRFAGKDSAGNATGKIVRWKPERSVGEEVANY
YIDPLYGFNKIVTAIMGKDRGDLHMLMQFAEREFNSISGIHALCEALEDCETRMLETIYHSKIAWHVVGV
AAESEEEFRRMPAYRIWVLCKYAISALDDGKGGVTLAMARFLTDIAEGNERGDLVRIALKEINRFCLAQD
PEQWRIAEQADEKLVELRNSVREIKLQDHIETRPQIAELLSRIGQLNFESLVETDDIVKARQISQEVGSA
LIDLIITVVSRGSLFQSSAQNAVEIIDNHTLGVMNYVIAGCVEHFDKVLGAIPGADKYITTDAIEKVMAE
IEANFDANDNLLGKTARIFRGHFVQNIKQMFCIPGSFSPAGLISLLPQTIFSTLNDLVYLDMITHNIKGY
RNISREHVNALQRKYFNSLADKGREELKANFKWVYYSLCKIGAIEDSEFLTKSQPLFFGENFFSLLQEYV
ASTTGEAPLPEHHPLEAITAMAESGAGAEPPAADADPNAVDDKKRTLVDPADIVPIQSSNVMLQGQEAYE
KKHFLVAQIYIRQIAEIKHLRPTGMEFPDADYINMLGPLDPEVAQLKATFNMRGGTGGYPAYDEYIRKFT
ITTTNAAGQPREVSFEVTLKSNNVESAQEVIAEQMQKNQEFMDFLADKNRKAAEFCVIADVRKSGGGRTQ
TVARQTVNHGIRFFDVQVDYQAVHKVKQDGYRAVIGVLPRYSPVNPTNPEALDYCRLMNDGQVLQEDRFK
VITEVPDRVVYIKGRMKGRQRELFYDEKNNLITTREGGTQQFSSVDSDFTREARIDHARKRGFFTVDTET
HVVPVATEFGKFKVFSKTPFIFRALAPIVSPLMTHTDVETLKRAIWAGQWVADRARTAPFMGLGVVHVDM
TKTASMGLGWPYVEAGVVGSAVGQTTLYVDFETISLMRTSMDEAAAASAASRALRNRVAANPFSERPTAI
MQFFWDHLHAQFNILATSLTDAVRFATPDIRFNEIIKAVFAPLTPYADVGQLKSLLLPSTQQQLRAAAQA
ASAQPNNAAAQAAAVSATRTVLRQNLYAAALNSPVAVMDAIARFDEMVSGISAAFAHDTLDLAVKHEADH
ASNNHAFTMSASDKESFYQEKIFYSFVKSLQIAAYGIEMSTAVGLNQQDAVATFYGHFDSAQHTAQRWGY
DEFNEVTAEQTSMVGEVWDKLKVLSRANPQDYERRIDQITRSLIKNPTPQTLKELGGIFAVANRHLTTGD
RHQLADTAYDFLSQKTQRALQNLQDFTHAVYTQIKMYGQDQNRYIQLLRSNNIIKQDYDVEWRSVTTFIK
EFDSSLTAIRNLDLTRCFRDIRMVFDPEDLQVKKQLTRIGGRAFATIYSRETQQPLYTVAHQLAISMDRQ
SGKKTLLFVSPLPIAPADLLQMVESTQRAMANPGFPALIGSTKDFMKNADILVNHEPKVSGKILDAKAFK
RLFETSATEVELRRSLSHHDISSALTPRTSARMFWDFTLVEPARSAGGTATAVKYQYSSYKGRTYKIHAK
RQLVNSASAQNTALPTSRPVELPITPVLVKVEYLKPNSTEIGRFTVVGADANGEPAFYGKNLSTDKTKIR
VLQLLSTVSRGKVDDIVSMKKLNAGESDYQDFMNKWNARNAQFKLKLSKITRLGMFATFIERLMEDCDVY
FSLGDVSWRQFMSRTDHNIDKVIAIMRALEFRKGAVERPDGLPYIPLTDIYNVIDVNFHRFQTRASALDM
LYELSMLELGNNKCATIAAPTEEDLLNKIANYCRSVQCEDVDDETYQQSADAGKTYNKNAVRMLRRSYGH
FIGHAARFSSKLNPAFHTAVVKQTMQKMEAQEARAVSQNDEILENLLEVLFPIYNSDADERKAYLYAGIK
LYLENQRLPLLARFLNEKVFYALDLMSLAADEWKVTISWSDEKIRFEHGPQGSVARTISFIGLMTTFSPT
DRRQFMRAFEECHALRGQKLFPEDKGDSEVTYTLIPAVSGQFNERFKISSTLKTTLDSLALKDSGLGYVL
PELDGSESTARNDLDYKDLRRKFNLDKLSTYGINRHLQTRKFNAKDTQAMAKTFSLWLPSSEIARHFYHT
GSEQAPSQAAPFTYLPADMQRDIGRMQAAVRSLPQALQGAEHAARAGMRYLMETYFTANTDAQRFSIIKH
LVQERQKNPLRYMLERTVAARFNVAEAVAARVGSARFPFDTSAAGLADGWSYIFLYLRALGHLPEERAFL
WNTLQEAFRLHHMELKTEIDYQQGSVSVTYDNKAYPDFADFKIADLEEFVDGLPPGTNFGIEYALQPVDP
DELPRSVFIVHGSRGPGDPHRKANDILLYLNHILPGPHIEILAHALIRQLDQKKGRQAPARAAT

>QHI97261.1 hypothetical protein GT347_04260 [Xylophilus rhododendri]
MTAEFNPSSFPKHGSTNSNITALTVLAKGLIHPAQRHSQNPLETNPQKTPIIAASNLPPQDINGEMTSLL
GLRHGASAHDSIYTPANVVANDESDHARRVRSRRDLRTISDQYGRYHYVSAQVEAQPHGQFAQWADGFYR
YYYKGGLQGQARYIVGVDGLIYQLFEPSHTGTSHAHAHAQDRTAVQPTQAPASPTTSRNPILPNWGPNQS
DGPPAGTVNPSKADYKNFGWDDPHPARPDQRLNHPTVRPGFQQPKPAYPASLGVNEQPAAPTRPAYVASA
STRDPSKAFYGSHQSDQPDPGTRKPLQTYKHPGFTQPSSRIAGRPFTKPPAPLYHSRYDENGQPVDDSED
YTAVTDYYGRTIKTHVVGYDMNSKKIFSHLGPRKLIGILPNKTAVYGDYERVLIGWDENRRPVYKGDNFK
QGTIDPQYLGRTASTPSYGKGFEFADTNWWEHNYTTTTRAPRIYISASRPSYLNKYDKYGHVLPDHENGR
NRNPAFNRNSQAGAGSPQRQANFIARERPKARYPLYFFEDITCQDEGWELPTVKSKRQLGVSLMPPAAPP
LPTGNKTIDGQHFAEYTAEMNIHRMQQKYFNRTLESEKPLMELLKEVEQEVLDAEQAQIENLQEKRESPE
QAVFAKLPIAHFTSDTLESFLEEVLPRYQDAYGQPLTIGQAGSSFHNYVAEEIKTTVLGYFKNPTPEPRK
QIIRLIKAFSALKTIFGNDASFMFTQSIQPGLMGENPRKISHSLHKSLADNLLSITYNPLFTMDAMRPLM
PLSHAGGIIYKVETELYKIHVITNIYFDPKNPDILGKISADLETVAKSALADEMLAKRTPGFPLTRTTKL

### Supplementary Data S2

HYRPRLEKYHPVSINTLDEYNRIQILYDAPQETSVRDEAESLEILARMPFGPAAQEIKKIITPIKMILDV
FFVPFAEEDSSGKPTGKIVRWTLESPSDFFSKYFIAPMGPLITGLNAISGSHSGEFHTFVDAAERYFSTI
PAINSLCAALEDCDNRPLESIYHSKLAFHLAGIADQCESEFTRIRAYRIWLLFKYASSAADDAKSTISLA
IARFMSEMANGDGMGEKVRIALKEINQQYLVNNPQMSQMAEQADAKLLELRNSVREIRLQDHIESKPQIA
DLLRRIGQMNFDSLVETDQIIKARQVTQQIGSAMVDIILTIASKGLLLPELIQNGVEILDDNTLGALNKI
ISGCVGEFDKMISPIAGSEKFLTTKAIEKVMVEIEQNFVKNDNALGKIARIFRGHLVSNLKLLFCVPGSQ
SISGLLSLIPQTIFSTINDFIFLDLIGHRLKEYQATTRQHVSDLQKKYYNGLLSKSRDELKSNFKWVYYS
LCTIGKIEDPVFLKESQPLFFGENYLSLAQEYVASITGEAPLPEFNPLTDGKPLPIDPNIPAQDEAEQED
ENDTVEKKRTFVNSADIVSPDAAKAAMGGKQIFNEKIVELSRIYGPKMNDIVDRMPKGIHFVDSDFVSVL
GPLDLEVAQLKATFNRRLDGTGGYPAYDEFIRTFTITTTNAVGLPEKISLEMVMKSGSIEYAQEFADEQI
RKNQQFMRFLAERSLMSANYCTIEDIQRSDGSRDQTVVRKTIEGDFRAFDIHVTYQEVHTESHGETEEII
GIIPTYTAVNPHDPHAVAFQNLMNGGQVLQEDRFKLITDVSDKIFYFKGQIESRLYELFYDYQLNFITAR
EAGGHLFLSEDNDFVKHIRTELTRKMAFSHLDTEPHVVPISTNFGKFKSFSKTPFIFSLPENAVSRTLFS
DDVATLRKAFDAAHWFVREANGTPFVGLGIANVKLVRTDFVSMQVPHVESAVVGSSIGETTLTVNFEIIH
RWRTQAADANTASAQSRDLRNQLAADPRAMTQKTLLSFFTSHFGNQFKILCTALGDASLSPPDGRFDKVL
DTILAPLTPYARVSRLKPLLNLNFRKKLRVDAAARAAAAPPNTVADALDAASPLRAISMADLNNALRNRP
AETIDAIARFDDMVAKVFSVYMHATLRVVIKHESDHASNVYSWMLGAMDAESIYQERIFYSLVRSVAIAS
KGIGMSAALSMDPREAVTTFYGNFDSGQYTPERMKYPDPELFIPEQTSVTGELWEELKILRQTNLADYNS
RIDQIAKSLIKNPSPKTFRELHAIFSVANRHITSGDCHQLADTSYNFLSEKTQAALQNLQDFAHAIYSQI
NSYGSDANRYIQLLRSQNFIQGNYQAEVEGITAFLDEFRTSIIARDGVDLTKCYRDIRMFFNPDDLPVKT
NSGPIRGRAFITIHSAETNQPLFTVAHKLSVITDQKTGEKTLIFVEPVSVSPANLQHMIASTQRSTGYAA
FPALIGTGKDFKKHAHTLVNFSPAIAGRTLDGQTFRNLFEATVSEIASRRPINQADRTTDLNPNGGIRIF
WDFTLLDKSSDAGSHPTAVKYQYASQGHHTFKFRPKRQNTNTANVQPRSPAIFQYRLLPSELTEPPVKPA
LIKAEYSIPDSDDLGQITTFAADVNNEPAFRGKNLSLKHTKTRILELLRATGRKVEDIVSMRQLSTSEID
YQKFMTEWSTRNALFKTKLSEINKVGMFETFMGRLMDDCDVYFSLGDISWRRYLSKTDDNIGKIVTIMHA
LEFRKGAVQRVDGKPYIGLVDIYNVIDINFYRFKTRASALDMLYELSSIELETNKCELISSASEDDLISK
IATYCRSVRCKEIDDEAFKKSEESGRVYNKNMVMTLRKYHGFFVGHARKFGSKINLPFHIHVVDQVILQM
EAESSKAVLQNDEIVESLLGVLFPMYQAHPEEIEPYFYASIKLCLDTKQFPILARFFNDKVLPVINLTKL
GTDESKTTLFWSEEEIRFEYGPQNSLIKHLSFKGLISGFSPDERARFLTALEDAFALRGQKLFLEEKKDA
SVIYTLIPAASTQFSERFKISSKLKKNLDRMALIESTKGYVLPELDHEKSNDQTAYEYRNLQGKYSLDKL
GTYAINRYLQTKAFAADDMDKLVKTFSTWLPSKEIARDYYENNAAFQSEPFSYSAADMQRDIAQMKAAVT
NLPQDIRRAGHAARAAMRYLIEKYAKADTDPQRFSIIKRLVQERGKDALLYFLERTVASNFDLLDQLAAH
PGGIKLQFSPFIGGASDNRHYIFAYLKKLAALPDECQFLFAALQDAFKLHNMKLQNNIDYAGNSVEVAID
NAVHQEFVDFKICDLEEFIAGLPNNTDFDIEYSSYQPSPDRQPVSVFTVGNSAANDARYKQANEILLYLN
NVIPGPSVENLARALKNQLESKKRKRN

>QHI97262.1 hypothetical protein GT347_04265 [Xylophilus rhododendri]
MAAEFNPSSSLQNASANSKISVLTVLTQGLIALKTQQSQNLPGGKSSKSEVIAEQKLPPNEVSEEMRNLL
GLKSGFKSHDNIDTSTETFPNDGHDHGRNGRSKRDLRAMQDRYGRYHYVTTLVEVQPGGQFLPWADGFYR
YYHKGVLQGQNIYIAGLDGRFYQVFQPNRADMAPGNVYVQGRTGIQATQAPPPASTSRNPSLPYWGPNQS
DQPPASTADPRKADYQNFGWFDANPARPDQRLNHAAAHPGFQAAQPVNPGAFGSHDRPNEPTRPPYIPSA
TGRDPSKAFYGSNQSDKPETGTRNPGQADNNKFHWTAATRSPMVHSPSTSTRRLSTKPPAPLYQSRYDEN
GQPVDESEDYTAVTDYYGRTIKTHVVGYDMNGKKIYSHVGPRKLIGMLPNKTAVYGDYERVLIGWDENRR
PVYLGDNFKQGTIDPQYLGRTASTPSYGKGFEFADTNWWEHNYTTTTRAPRIYISTSRPSNLNIYDKHGH
FIGDAETGRARGRPGQHTHDGPGNRLGNNGDIPETTTKDPRAAIYPVALDRSGKPTRINQAWRTPTDRLA
PDYGPNNDPAYQQRYSAAGRSSQRYGNSSNRASWQLARPKYPQYFIFEDITYRAMNQALATVRSKRQIGV
KLTEPAPPPFKPATVLSTVKTWPSTPHEKISTKCNRTISTPLRGSKSR

>QHI97263.1 hypothetical protein GT347_04270 [Xylophilus rhododendri]
MQQDHFNAITGFEKPLMDTIKELEREIRDAEQAQLTDSLDKRETTEDTLIAKLPVAHFTSPTMETILEEL
LPRYEDAHGTQLTEAQASNSYHNQVAETIKTQILQYFRNPTPAAEINIIRFVKASAGFKAAFGKDSSFIF
TPSGQAGLPGEPPRAISYSLRYSLSNTLTSILFNPIYDIVPGTSPQIVGQTGASVQKVETDLYSVQVVAS
MYYEISSPDALTAASELLEKCVKTALADEMIARRTPGIPLVRTSNFAYTAEADKYHPVTITTVEEYERLR
TAYGFAPRISTRDERAPLPFMARMPFSLAAQEQRKIFLPVKTMLDQFFVLFAEKDSSGQSTGKIVRWTIE
SAAEKVSKYFLDPINGLKFFLTSVSGPHTIQFAAFSDDAERYFSSIPGIFGLCEALEECQQTVLENVQDS
KIAWNAAGITSGSRTEFMRIRAYRIWLLAKYASLATEDSRAGFTLAMGKFLSDIADGTGRGELIRMALRE
VNRLSLRNQPEQWKIAEQADAKLVELRTSVREIKLQDHIESRPQIADLLGRIGQLNFESLTSADQLMKAR
QISQQVGSAFIDVLMTIVTKGALLPNLAQNTVEAIDNNTLGLMNYLISGCVGELDAVISAIPAGKKYLTT
EAIEKVMSEIETTFSQNDNKLGNISRIFRSHLIGNIKTMFCVPGSFSPAGLISAIPQTIFSTLNDMIFAD
MISANIQGYKIASREHVNALQRKYFNSLVSKGREELKENFKWVYYSLCKIGSIEDPAFVRENPILFFKEN
YQSLLQEYVALTTGEAPQPEFTGPLSSIRFAPTNLTQADGTPAARQNGAEIGKRVLVDAAEVQDVNDTWL
AKQGDEIFLQQMRQLDLIHGRKYEELCYLLPPGATLTDRDFFHVLGPLDPEVEQLKNTFNAKPDGSGGYP
AYHEFIRKFSITTYRGIVPTHVSFEITLPTNNIEIAQELTADELQKNDEFMQFISDKNTRAAKFVVIRDQ
ERPDGTREQMVVRNTPGNYRIFNVQVSYQAVYDERHAGIRDKIGILPHYSPLHAFDQHASTYCDLLNYGQ
VLQEDSFRRITEVTDTVFVINARFMHKRYELYYDNNRNLITTRPAGTRVFSLKDDAFTEQARKDHAARRG
LGFFATHRHVVPVPTAFGRFRSFSKTPFIFATKTPLAVSTPLQAQDIAEFEDLFRAAEYVASHARNIYMG
LGVVNVKFENSGIAGLSVPYIEKLAPGSSIGQTTLNVNHRFFAQLDHDLATANALRSEARTLKNTLDVYS
RHRPPQALLDIYRQHLDAQFHVLLSALPEWARTPADSRFEQVVDTVFAPLAPFVDSYRLKHRLNDLEISR
IRAAAPGNPAAASPVDLLIDKNILQVARSNPSYLMECLANFDARVAQIFEIKRNHQRAGTIRHEADHAAN
LDSAQVGAQDADSVQQERVFYSLVKNLISASTGVEMSEVLGLSRHQAIIRFYGTFVYDQAYPERFGYTAT

### Supplementary Data S2

```
DDIIAEQSSVTGEIWERLKVVSRDQPAEYLHVMDQIARSLLMDVSAGGFATMKAVFDVATRHVTSGDRHE
LADTSYDFVSRRTQRAAQNLQDFAHGLLMQIKALATTKKDYVRVLRSLNVIQNDYSTEWHQMKLFADAFH
MSPVNQIDLDLTSCYREVRMYFNPDEFPVTSVPTRINGRAFITLHRKDTAQPIYTVAFGLYVEKNTRTNE
NIMKLVDPLAVAPNALRAVIRETARYTGWPSFPALVGSVKHFPDLAGTVANFATAVAGKTMDERAFSGLF
ESSLVDMAHHRPMGPRDVAAALNPSDMRTFYDFTLLDAEGEGAPVQDVNRAGGVKYLWARETYGVPRTFK
ARPKRDSGGPTDRAITPLLVKLEYRIPNSTEVGNITFVAADIYNQPAFYGFHPSTHFTKIQALQLLSSLA
HVKADAFVGITQLKTGEKAYQDFMSAWNARQAAFKAKLSTISKIGMYATFMESLKADCDVYFPLGSVSWR
SFPSRTDGNIDKVLTVMRALEFKKGAAQRPDGAPYVVLTDLCNVIDVNFYRFRTDASALDMLYELSSIEL
SRNGCAVISGATELDLLEKIEIYCRGVQCNAVEEEKYAKPDDSRSSYNKHTVRELRKQYSYFLGYVGRFG
SDLDVGFHFNVVRPVMQAMERGRDTAVSLAEEITDSMLGLLQPQYQNNPADRRAFMYAGIKLCLENGQPA
SLAKFLNNAVFSDLGLAEIAGGEWNVLVSWIRQEIRFQYSGPGLFTRTISLAGLASGFSVDEKFHFLRAV
EESHALKGHKLILEDKTDEELTYTFIPANSAQYSERFKISSKLKASLDSMPLEAGSKGYVMPKLDHSRSS
AQNKNDYENLKIRFELDKLNSYGMNRYLQAREPSASDMEMNVKIFSQWLPSSELARLLSQASPQAAEEDG
IFQYYAADMHADILQMSEAVIALPQDLRGTVHAARVALRHLMDTYFTVATDAQRFSIIKRLIQGRQAKPL
VYILEQTVTSKFDEIEMLAAQLGGAKIGFEVAAGGSLDNWSYVFRYLQALLPLPEERRFLWDRLQDAFKL
HRLQLADEINYAEGSIRLTYDSTIYPELAQFRFADLEEFVSRLPQGTSYRIGFKARQTAPGQLPRSVFDV
DESKIDQGPYSKANKVLAYFNNVLSGPQVEYVAVYFTRLLEMKNHAQGSGQALN
```

```
QHI97260.1      -MHLYPISTPSNNSSAPSVVSTLALLAQGVIQRYLPSHVNPTPR------VVENDQLPSQ
QHI97263.1      ------------------------------------------------------------
QHI97411.1      MTAEFNPSSSANNSAAHSSISALTLLTQSLLRRNQKQPQDTLQTDS----IIKEPNLPPR
QHI97261.1      MTAEFNPSSFPKHGSTNSNITALTVLAKGLIHPAQRHSQNPLETNPQKTPIIAASNLPPQ
QHI97262.1      MAAEFNPSSSLQNASANSKISVLTVLTQGLIALKTQQSQNLPGGKSSKSEVIAEQKLPPN


QHI97260.1      GISHEMAGLLAL-------------------DETHAKTQSRSKRESSA--SDYRRKETTE
QHI97263.1      ------------------------------------------------------------
QHI97411.1      EISEEMNKLLNLKSTEASHESLHPPAEVPANSESDKPRHGRSRRELRAMQDRNGRHHYVT
QHI97261.1      DINGEMTSLLGLRHGASAHDSIYTPANVVANDESDHARRVRSRRDLRTISDQYGRYHYVS
QHI97262.1      EVSEEMRNLLGLKSGFKSHDNIDTSTETFPNDGHDHGRNGRSKRDLRAMQDRYGRYHYVT


QHI97260.1      PDYDANQGGR--------------------------------------------------
QHI97263.1      ------------------------------------------------------------
QHI97411.1      AQVEAQHGGQFVEWATGLYHYYHNGVLQGQTSYIVGLDGSIYQIFEPQHAGPAPASAYSQ
QHI97261.1      AQVEAQPHGQFAQWADGFYRYYYKGGLQGQARYIVGVDGLIYQLFEPSHTGTSHAHAHAQ
QHI97262.1      TLVEVQPGGQFLPWADGFYRYYHKGVLQGQNIYIAGLDGRFYQVFQPNRADMAPGNVYVQ


QHI97260.1      ------------------------------------------------------------
QHI97263.1      ------------------------------------------------------------
QHI97411.1      GQTGIRTTQAPPQITTSRNPKLPYWGPNQSDLPGVS------------------------
QHI97261.1      DRTAVQPTQAPASPTTSRNPILPNWGPNQSDGPPAGTVNPSKADYKNFGWDDPHPARPDQ
QHI97262.1      GRTGIQATQAPPPASTSRNPSLPYWGPNQSDQPPASTADPRKADYQNFGWFDANPARPDQ


QHI97260.1      ------------------------------------------------------------
QHI97263.1      ------------------------------------------------------------
QHI97411.1      ------------------------------------------------------------
QHI97261.1      RLNHPTVRPGFQQPKPAYPASLGVNEQPAAPTRPAYVASASTRDPSKAFYGSHQSDQPDP
QHI97262.1      RLNHAAAHPGFQAAQPVNPGAFGSHDRPNEPTRPPYIPSATGRDPSKAFYGSNQSDKPET


QHI97260.1      ----------------------AFDKPSSIPRTNALPSDPVPRHAYYQRALNKAPQKPEQ
QHI97263.1      -------------------------------------------------------MQQDH
QHI97411.1      -TRKPGQAGNNNFDGNAATRAGVFQKSSLA-ARHFSTKRPAPLYHSKYDENGRPVDASQE
QHI97261.1      GTRKPLQ----------TYKHPGFTQPSSRIAGRPFTKPPAPLYHSRYDENGQPVDDSED
QHI97262.1      GTRNPGQADNNKFHWTAATRSPMVHSPSTS-TRRLSTKPPAPLYQSRYDENGQPVDESED
                                                                          :
```

**Supplementary Data S2**

```
QHI97260.1        SDTVTDYYGRPVNPPPGAYGFNSETHPSGRGRKQFLGYDMTGKPIYRADSRYLVGWDEQR
QHI97263.1        FNAITGF-----------------------------------------------------
QHI97411.1        YTGVTDYYGRTVKTHVVGYDMNGKEIFAHLGRRKLIGMLPNKTAVYGDYERVLIGWDENR
QHI97261.1        YTAVTDYYGRTIKTHVVGYDMNSKKIFSHLGPRKLIGILPNKTAVYGDYERVLIGWDENR
QHI97262.1        YTAVTDYYGRTIKTHVVGYDMNGKKIYSHVGPRKLIGMLPNKTAVYGDYERVLIGWDENR
                     :*.:

QHI97260.1        KPIYSDDVRPQPSIDPSLIGNIPSTPAYGKGFEFADTNWWD--YTTTTRQPRKKAPAKKP
QHI97263.1        ------------------------------------------------------------
QHI97411.1        RPMYMGDNRPQGTIDPQYLGRTASTPAYGKGFEFADTNWWEHNYTTTTRGPRINIPTSRP
QHI97261.1        RPVYKGDNFKQGTIDPQYLGRTASTPSYGKGFEFADTNWWEHNYTTTTRAPRIYISASRP
QHI97262.1        RPVYLGDNFKQGTIDPQYLGRTASTPSYGKGFEFADTNWWEHNYTTTTRAPRIYISTSRP


QHI97260.1        SHLNQYDADGNILPSKAT------------------------------------------
QHI97263.1        ------------------------------------------------------------
QHI97411.1        SYLNQYDKYGHVTRDPETEKTRGRGRSRAGQHQPAGPGNPAPIQGQQPGHRPPDNRAPHY
QHI97261.1        SYLNKYDKYGHVLPDHEN------------------------------------------
QHI97262.1        SNLNIYDKHGHFIGDAET------------------------------------------


QHI97260.1        ------------------------------------------------------------
QHI97263.1        ---------------------------------EKP------------------------
QHI97411.1        GENSDPGHKYRNPAGGHLPQRHGSSNQLARWQVARP--KYPQYFVFEDIIYKGVDQMFAS
QHI97261.1        GRNRNPAFNRNSQAGAGSPQRQ--ANFIAR---ERPKARYPLYF-FEDITCQDEGWELPT
QHI97262.1        ------------------------------------------------------------


QHI97260.1        ------------------------------------------------------------
QHI97263.1        ---------------------------------------------------------LMD
QHI97411.1        VRSKRQIGVRFTELAAPPIPTGNQSIDIQAWTDYAAQNNIYQMQQRQFSSTIESEKPLMD
QHI97261.1        VKSKRQLGVSLMPPAAPPLPTGNKTIDGQHFAEYTAEMNIHRMQQKYFNRTLESEKPLME
QHI97262.1        ------------------------------------------------------------


QHI97260.1        ------------------------------------------------------------
QHI97263.1        TIKELEREIRDAEQAQLTDSLDKRETTEDTLIAKLPVAHFTSPTMETILEELLPRYEDAH
QHI97411.1        TIKTLEREMFEAEQARMENAQQKRESVDETAVAKLPVAYFTSETMDALLEEVLPRYRDAY
QHI97261.1        LLKEVEQEVLDAEQAQIENLQEKRESPEQAVFAKLPIAHFTSDTLESFLEEVLPRYQDAY
QHI97262.1        ------------------------------------------------------------


QHI97260.1        ------------------------------------------------------------
QHI97263.1        GTQLTEAQASNSYHNQVAETIKTQILQYFRNPTPAAEINIIRFVKASAGFKAAFGKDSSF
QHI97411.1        GNALTEVQAEESFHNKIAEDIKTRIFQYFRNPESELEKKIIRFIKACAGFKTIFGKNASF
QHI97261.1        GQPLTIGQAGSSFHNYVAEEIKTTVLGYFKNPTPEPRKQIIRLIKAFSALKTIFGNDASF
QHI97262.1        ------------------------------------------------------------


QHI97260.1        ------------------------------------------------------------
QHI97263.1        IFTPSGQAGLPGEPPRAISYSLRYSLSNTLTSILFNPIYDIVPGTSPQIVGQTGASVQKV
QHI97411.1        VFTPTLQKGLVGEYPRNISLELRYALADKLLNIQYNPIYIAAGPPDHPYIHLSNAFVQKI
QHI97261.1        MFTQSIQPGLMGENPRKISHSLHKSLADNLLSITYNPLFTMDAMRPLMPLSHAGGIIYKV
QHI97262.1        ----GRARGRPGQHTHD-------------------------------------------


QHI97260.1        ------------------------------------------------------------
QHI97263.1        ETDLYSVQVVASMYYEISSPDALTAASELLEKCVKTALADEMIARRTPGIPLVRTSNFAY
QHI97411.1        ETELYDTQTIVNIYYDPAHADALTRASEGLEKTAKTRIADEMIARRTPGFPLVRTSKAFF
QHI97261.1        ETELYKIHVITNIYFDPKNPDILGKISADLETVAKSALADEMLAKRTPGFPLTRTTKLHY
QHI97262.1        ----------------------------------------------GPGNRLGNNGDI--
```

### Supplementary Data S2

```
QHI97260.1      ------------------------------------------------------------
QHI97263.1      TAEADKYHPVTITTVEEYERLRTAYGFAPRISTRDERAPLPFMARMPFSLAAQEQRKIFL
QHI97411.1      TPESGKYSPVSINTLEEYEQVVIAYDLEPGVSIRDERAPLEVLARMPFQQAGQELRKIFL
QHI97261.1      RPRLEKYHPVSINTLDEYNRIQILYDAPQETSVRDEAESLEILARMPFGPAAQEIKKIIT
QHI97262.1      ----------------------------PETTTKDPRAAIYPVAL---------------

QHI97260.1      ----------------RREGKNQ-------------------------------------
QHI97263.1      PVKTMLDQFFVLFAEKDSSGQSTGKIVRWTIE-SAAEKVSKYFLDPINGLKFFLTSVSGP
QHI97411.1      PLKSRLDRVFVRFAGKDSAGNATGKIVRWKPERSVGEEVANYYIDPLYGFNKIVTAIMGK
QHI97261.1      PIKMILDVFFVPFAEEDSSGKPTGKIVRWTLE-SPSDFFSKYFIAPMGPLITGLNAISGS
QHI97262.1      ----------------DRSGKPTRINQAWR------------------------------
                                   *:

QHI97260.1      ------------------------------------------------------------
QHI97263.1      HTIQFAAFSDDAERYFSSIPGIFGLCEALEECQQTVLENVQDSKIAWNAAGITSGSRTEF
QHI97411.1      DRGDLHMLMQFAEREFNSISGIHALCEALEDCETRMLETIYHSKIAWHVVGVAAESEEEF
QHI97261.1      HSGEFHTFVDAAERYFSTIPAINSLCAALEDCDNRPLESIYHSKLAFHLAGIADQCESEF
QHI97262.1      ------------------------------------------------------------

QHI97260.1      ------------------------------------------------------------
QHI97263.1      MRIRAYRIWLLAKYASLATEDSRAGFTLAMGKFLSDIADGTGRGELIRMALREVNRLSLR
QHI97411.1      RRMPAYRIWVLCKYAISALDDGKGGVTLAMARFLTDIAEGNERGDLVRIALKEINRFCLA
QHI97261.1      TRIRAYRIWLLFKYASSAADDAKSTISLAIARFMSEMANGDGMGEKVRIALKEINQQYLV
QHI97262.1      ------------------------------------------------------------

QHI97260.1      ------------------------------------------------------------
QHI97263.1      NQPEQWKIAEQADAKLVELRTSVREIKLQDHIESRPQIADLLGRIGQLNFESLTSADQLM
QHI97411.1      QDPEQWRIAEQADEKLVELRNSVREIKLQDHIETRPQIAELLSRIGQLNFESLVETDDIV
QHI97261.1      NNPQMSQMAEQADAKLLELRNSVREIRLQDHIESKPQIADLLRRIGQMNFDSLVETDQII
QHI97262.1      ------------------------------------------------------------

QHI97260.1      ------------------------------------------------------------
QHI97263.1      KARQISQQVGSAFIDVLMTIVTKGALLPNLAQNTVEAIDNNTLGLMNYLISGCVGELDAV
QHI97411.1      KARQISQEVGSALIDLIITVVSRGSLFQSSAQNAVEIIDNHTLGVMNYVIAGCVEHFDKV
QHI97261.1      KARQVTQQIGSAMVDIILTIASKGLLLPELIQNGVEILDDNTLGALNKIISGCVGEFDKM
QHI97262.1      ------------------------------------------------------------

QHI97260.1      ------------------------------------------------------------
QHI97263.1      ISAIPAGKKYLTTEAIEKVMSEIETTFSQNDNKLGNISRIFRSHLIGNIKTMFCVPGSFS
QHI97411.1      LGAIPGADKYITTDAIEKVMAEIEANFDANDNLLGKTARIFRGHFVQNIKQMFCIPGSFS
QHI97261.1      ISPIAGSEKFLTTKAIEKVMVEIEQNFVKNDNALGKIARIFRGHLVSNLKLLFCVPGSQS
QHI97262.1      ------------------------------------------------------------

QHI97260.1      ------------------------------------------------------------
QHI97263.1      PAGLISAIPQTIFSTLNDMIFADMISANIQGYKIASREHVNALQRKYFNSLVSKGREELK
QHI97411.1      PAGLISLLPQTIFSTLNDLVYLDMITHNIKGYRNISREHVNALQRKYFNSLADKGREELK
QHI97261.1      ISGLLSLIPQTIFSTINDFIFLDLIGHRLKEYQATTRQHVSDLQKKYYNGLLSKSRDELK
QHI97262.1      ------------------------------------------------------------

QHI97260.1      ----------------------------------------YIDGGTG-------------
QHI97263.1      ENFKWVYYSLCKIGSIEDPAFVRENPILFFKENYQSLLQEYVALTTGEAPQPEFTGPLSS
QHI97411.1      ANFKWVYYSLCKIGAIEDSEFLTKSQPLFFGENFFSLLQEYVASTTGEAPLPEHH-PLEA
QHI97261.1      SNFKWVYYSLCTIGKIEDPVFLKESQPLFFGENYLSLAQEYVASITGEAPLPEFN-PLTD
QHI97262.1      ------------------------------------------------------------
```

**Supplementary Data S2**

```
QHI97260.1      -----------------------------TVLASSE------------------------
QHI97263.1      IRFAP--TNLTQADGTPAARQNGAEIGKRVLVDAAEVQDVNDTWLAKQGDEIFLQQMRQL
QHI97411.1      ITAMA-ESGAGAEPPAADADPNAVDDKKRTLVDPADIVPIQSSNVMLQGQEAYEKKHFLV
QHI97261.1      GKPLPIDPNIPAQDEAEQEDENDTVEKKRTFVNSADIVSPDAAKAAMGGKQIFNEKIVEL
QHI97262.1      --------------------------------TPTDRLAPD-------------------
                                                 .::

QHI97260.1      ------------------------------------------------------PMLHNF
QHI97263.1      DLIHGRKYEELCYLLPPGATLTDRDFFHVLGPLDPEVEQLKNTFNAKPDGSGGYPAYHEF
QHI97411.1      AQIYIRQIAEIKHLRPTGMEFPDADYINMLGPLDPEVAQLKATFNMR-GGTGGYPAYDEY
QHI97261.1      SRIYGPKMNDIVDRMPKGIHFVDSDFVSVLGPLDLEVAQLKATFNRRLDGTGGYPAYDEF
QHI97262.1      ---YGPNND---------------------------------------------PAYQQ-
                                                                      *   :

QHI97260.1      IHD---------------------------------------------------------
QHI97263.1      IRKFSITTYRGI-VPTHVSFEITLPTNNIEIAQELTADELQKNDEFMQFISDKNTRAAKF
QHI97411.1      IRKFTITTTNAAGQPREVSFEVTLKSNNVESAQEVIAEQMQKNQEFMDFLADKNRKAAEF
QHI97261.1      IRTFTITTTNAVGLPEKISLEMVMKSGSIEYAQEFADEQIRKNQQFMRFLAERSLMSANY
QHI97262.1      ------------------------------------------------------------


QHI97260.1      ------------------------------------------------------------
QHI97263.1      VVIRDQERPDGTREQMVVRNT-PGNYRIFNVQVSYQAVYDERHAGIRDKIGILPHYSPLH
QHI97411.1      CVIADVRKSGGGRTQTVARQTVNHGIRFFDVQVDYQAVHKVKQDGYRAVIGVLPRYSPVN
QHI97261.1      CTIEDIQRSDGSRDQTVVRKTIEGDFRAFDIHVTYQEVHTESHGETEEIIGIIPTYTAVN
QHI97262.1      ------------------------------------------------------------


QHI97260.1      ------------------------------------------------------------
QHI97263.1      AFDQHASTYCDLLNYGQVLQEDSFRRITEVTDTVFVINARFMHKRYELYYDNNRNLITTR
QHI97411.1      PTNPEALDYCRLMNDGQVLQEDRFKVITEVPDRVVYIKGRMKGRQRELFYDEKNNLITTR
QHI97261.1      PHDPHAVAFQNLMNGGQVLQEDRFKLITDVSDKIFYFKGQIESRLYELFYDYQLNFITAR
QHI97262.1      ------------------------------------------------------------


QHI97260.1      ------------------------------------------------------------
QHI97263.1      PAGTRVFSLKDDAFTEQARKDHAARRGLGFFATHRHVVPVPTAFGRFRSFSKTPFIFATK
QHI97411.1      EGGTQQFSSVDSDFTREARIDHARKRGFFTVDTETHVVPVATEFGKFKVFSKTPFIFRAL
QHI97261.1      EAGGHLFLSEDNDFVKHIRTELTRKMAFSHLDTEPHVVPISTNFGKFKSFSKTPFIFSLP
QHI97262.1      ------------------------------------------------------------


QHI97260.1      ------------------------------------------------------------
QHI97263.1      TPLAVSTPLQAQDIAEFEDLFRAAEYVASHARNI-YMGLGVVNVKFENSGIAGLSVPYIE
QHI97411.1      AP-IVSPLMTHTDVETLKRAIWAGQWVADRARTAPFMGLGVVHVDMTKTASMGLGWPYVE
QHI97261.1      EN-AVSRTLFSDDVATLRKAFDAAHWFVREANGTPFVGLGIANVKLVRTDFVSMQVPHVE
QHI97262.1      ------------------------------------------------------------


QHI97260.1      ------------------------------------------------------------
QHI97263.1      KLAPGSSIGQTTLNVNHRFFAQLDHDLATANALRSEARTLKNTLDVYSRHRPPQALLDIY
QHI97411.1      AGVVGSAVGQTTLYVDFETISLMRTSMDEAAAASAASRALRNRVAANPFSERPTAIMQFF
QHI97261.1      SAVVGSSIGETTLTVNFEIIHRWRTQAADANTASAQSRDLRNQLAADPRAMTQKTLLSFF
QHI97262.1      ------------------------------------------------------------


QHI97260.1      ------------------------------------------------------------
QHI97263.1      RQHLDAQFHVLLSALPEWAR-TPADSRFEQVVDTVFAPLAPFVDSYRLKHRLNDLEISRI
QHI97411.1      WDHLHAQFNILATSLTDAVRFATPDIRFNEIIKAVFAPLTPYADVGQLKSLLLPSTQQQL
QHI97261.1      TSHFGNQFKILCTALGDASL-SPPDGRFDKVLDTILAPLTPYARVSRLKPLLNLNFRKKL
QHI97262.1      ------------------------------------------------------------
```

### Supplementary Data S2

```
QHI97260.1      ------------------------------------------------------------
QHI97263.1      RAAAPGNPAAASPVDL------------LIDKNILQVARSNPSYLMECLANFDARVAQIF
QHI97411.1      R--AAAQAASAQPNNAAAQAAAVSATRTVLRQNLYAAALNSPVAVMDAIARFDEMVSGIS
QHI97261.1      RVDAAARAAAAPPNTVADALDAASPLRAISMADLNNALRNRPAETIDAIARFDDMVAKVF
QHI97262.1      RYSAAGRSS---------------------------------------------------

QHI97260.1      ------------------------------------------------------------
QHI97263.1      EIKRNHQRAGTIRHEADHAANLDSAQVGAQDADSVQQERVFYSLVKNLISASTGVEMSEV
QHI97411.1      AAFAHDTLDLAVKHEADHASNNHAFTMSASDKESFYQEKIFYSFVKSLQIAAYGIEMSTA
QHI97261.1      SVYMHATLRVVIKHESDHASNVYSWMLGAMDAESIYQERIFYSLVRSVAIASKGIGMSAA
QHI97262.1      -------------QRYGNSSNRASWQLARPKYPQYF------------------------

QHI97260.1      ------------------------------------------------------------
QHI97263.1      LGLSRHQAIIRFYGTFVYDQAYPERFGYTATDDIIAEQSSVTGEIWERLKVVSRDQPAEY
QHI97411.1      VGLNQQDAVATFYGHFDSAQHTAQRWGYDEFNEVTAEQTSMVGEVWDKLKVLSRANPQDY
QHI97261.1      LSMDPREAVTTFYGNFDSGQYTPERMKYPDPELFIPEQTSVTGELWEELKILRQTNLADY
QHI97262.1      ------------------------------------------------------------

QHI97260.1      ------------------------------------------------------------
QHI97263.1      LHVMDQIARSLLMDVSAGGFATMKAVFDVATRHVTSGDRHELADTSYDFVSRRTQRAAQN
QHI97411.1      ERRIDQITRSLIKNPTPQTLKELGGIFAVANRHLTTGDRHQLADTAYDFLSQKTQRALQN
QHI97261.1      NSRIDQIAKSLIKNPSPKTFRELHAIFSVANRHITSGDCHQLADTSYNFLSEKTQAALQN
QHI97262.1      ------------------------------------------------------------

QHI97260.1      ------------------------------------------------------------
QHI97263.1      LQDFAHGLLMQIKALATTKKDYVRVLRSLNVIQNDYSTEWHQMKLFADAFHMSPVNQIDL
QHI97411.1      LQDFTHAVYTQIKMYGQDQNRYIQLLRSNNIIKQDYDVEWRSVTTFIKEFDSSLTAIRNL
QHI97261.1      LQDFAHAIYSQINSYGSDANRYIQLLRSQNFIQGNYQAEVEGITAFLDEFRTSIIARDGV
QHI97262.1      ------------------------------------------------------------

QHI97260.1      ------------------------------------------------------------
QHI97263.1      DLTSCYREVRMYFNPDEFPVTSVPTRINGRAFITLHRKDTAQPIYTVAFGLYVEKNTRTN
QHI97411.1      DLTRCFRDIRMVFDPEDLQVKKQLTRIGGRAFATIYSRETQQPLYTVAHQLAISMDRQSG
QHI97261.1      DLTKCYRDIRMFFNPDDLPVKTNSGPIRGRAFITIHSAETNQPLFTVAHKLSVITDQKTG
QHI97262.1      ------------------------------IFEDITYRAMNQALATVRS-----------

QHI97260.1      ------------------------------------------------------------
QHI97263.1      ENIMKLVDPLAVAPNALRAVIRETARYTGWPSFPALVGSVKHFPDLAGTVANFATAVAGK
QHI97411.1      KKTLLFVSPLPIAPADLLQMVESTQRAMANPGFPALIGSTKDFMKNADILVNHEPKVSGK
QHI97261.1      EKTLIFVEPVSVSPANLQHMIASTQRSTGYAAFPALIGTGKDFKKHAHTLVNFSPAIAGR
QHI97262.1      ------------------------------------------------------------

QHI97260.1      ------------------------------------------------------------
QHI97263.1      TMDERAFSGLFESSLVDMAHHRPMGPRDVAAALNPS-DMRTFYDFTLLDAEGEGAPVQDV
QHI97411.1      ILDAKAFKRLFETSATEVELRRSLSHHDISSALTPRTSARMFWDFTLVEPARSAG-----
QHI97261.1      TLDGQTFRNLFEATVSEIASRRPINQADRTTDLNPNGGIRIFWDFTLLDKSSDAG-----
QHI97262.1      ------------------------------------------------------------

QHI97260.1      -------------------------------------------------PAPSDSP----
QHI97263.1      NRAGGVKYLWARETYGVPRTFKARPKRDS--------------------GGPTDRAITPL
QHI97411.1      GTATAVKYQYSSYKG---RTYKIHAKRQLVNSASAQNTA-------LPTSRPVELPITPV
QHI97261.1      SHPTAVKYQYASQGH---HTFKFRPKRQNTNTANVQPRSPAIFQYRLLPSELTEPPVKPA
QHI97262.1      KRQIGVKL-----------------------------------------TEPAPPPFKPA
                                                               .
```

**Supplementary Data S2**

```
QHI97260.1      ------------------------------------------------------------
QHI97263.1      LVKLEYRIPNSTEVGNITFVAADIYNQPAFYGFHPSTHFTKIQALQLLSSLAHVKADAFV
QHI97411.1      LVKVEYLKPNSTEIGRFTVVGADANGEPAFYGKNLSTDKTKIRVLQLLSTVSRGKVDDIV
QHI97261.1      LIKAEYSIPDSDDLGQITTFAADVNNEPAFRGKNLSLKHTKTRILELLRATGR-KVEDIV
QHI97262.1      TV----------------------------------------------------------

QHI97260.1      ------------------------------------------------------------
QHI97263.1      GITQLKTGEKAYQDFMSAWNARQAAFKAKLSTISKIGMYATFMESLKADCDVYFPLGSVS
QHI97411.1      SMKKLNAGESDYQDFMNKWNARNAQFKLKLSKITRLGMFATFIERLMEDCDVYFSLGDVS
QHI97261.1      SMRQLSTSEIDYQKFMTEWSTRNALFKTKLSEINKVGMFETFMGRLMDDCDVYFSLGDIS
QHI97262.1      -----------------------------LSTVK-------------------------T

QHI97260.1      ------------------------------------------------------------
QHI97263.1      WRSFPSRTDGNIDKVLTVMRALEFKKGAAQRPDGAPYVVLTDLCNVIDVNFYRFRTDASA
QHI97411.1      WRQFMSRTDHNIDKVIAIMRALEFRKGAVERPDGLPYIPLTDIYNVIDVNFHRFQTRASA
QHI97261.1      WRRYLSKTDDNIGKIVTIMHALEFRKGAVQRVDGKPYIGLVDIYNVIDINFYRFKTRASA
QHI97262.1      WPSTPH------------------------------------------------------

QHI97260.1      ------------------------------------------------------------
QHI97263.1      LDMLYELSSIELSRNGCAVISGATELDLLEKIEIYCRGVQCNAVEEEKYAKPDDSRSSYN
QHI97411.1      LDMLYELSMLELGNNKCATIAAPTEEDLLNKIANYCRSVQCEDVDDETYQQSADAGKTYN
QHI97261.1      LDMLYELSSIELETNKCELISSASEDDLISKIATYCRSVRCKEIDDEAFKKSEESGRVYN
QHI97262.1      -----------------------------EKISTKCN-----------------------

QHI97260.1      -------KKQHRFFI---------------------------------------------
QHI97263.1      KHTVRELRKQYSYFLGYVGRFGSDLDVGFHFNVVRPVMQAMERGRDTAVSLAEEITDSML
QHI97411.1      KNAVRMLRRSYGHFIGHAARFSSKLNPAFHTAVVKQTMQKMEAQEARAVSQNDEILENLL
QHI97261.1      KNMVMTLRKYHGFFVGHARKFGSKINLPFHIHVVDQVILQMEAESSKAVLQNDEIVESLL
QHI97262.1      ------------------------------------------------------------

QHI97260.1      ------------------------------------------------------------
QHI97263.1      GLLQPQYQNNPADRRAFMYAGIKLCLENGQPASLAKFLNNAVFSDLGLAEIAGGEWNVLV
QHI97411.1      EVLFPIYNSDADERKAYLYAGIKLYLENQRLPLLARFLNEKVFYALDLMSLAADEWKVTI
QHI97261.1      GVLFPMYQAHPEEIEPYFYASIKLCLDTKQFPILARFFNDKVLPVINLTKLGTDESKTTL
QHI97262.1      ------------------------------------------------------------

QHI97260.1      ------------------------------------------------------------
QHI97263.1      SWIRQEIRFQYSGPGLFTRTISLAGLASGFSVDEKFHFLRAVEESHALKGHKLILEDKTD
QHI97411.1      SWSDEKIRFEHGPQGSVARTISFIGLMTTFSPTDRRQFMRAFEECHALRGQKLFPEDKGD
QHI97261.1      FWSEEEIRFEYGPQNSLIKHLSFKGLISGFSPDERARFLTALEDAFALRGQKLFLEEKKD
QHI97262.1      ------------------------------------------------------------

QHI97260.1      ------------------------------------------------------------
QHI97263.1      EELTYTFIPANSAQYSERFKISSKLKASLDSMPLEAGSKGYVMPKLDHSRSSAQNKNDYE
QHI97411.1      SEVTYTLIPAVSGQFNERFKISSTLKTTLDSLALKDSGLGYVLPELDGSESTARNDLDYK
QHI97261.1      ASVIYTLIPAASTQFSERFKISSKLKKNLDRMALIESTKGYVLPELDHEKSNDQTAYEYR
QHI97262.1      ------------------------------------------------------------

QHI97260.1      ------------------------------------------------------------
QHI97263.1      NLKIRFELDKLNSYGMNRYLQAREPSASDMEMNVKIFSQWLPSSELARLLSQA-SPQAAE
QHI97411.1      DLRRKFNLDKLSTYGINRHLQTRKFNAKDTQAMAKTFSLWLPSSEIARHFYHTGSEQAPS
QHI97261.1      NLQGKYSLDKLGTYAINRYLQTKAFAADDMDKLVKTFSTWLPSKEIARDYYEN---NAAF
QHI97262.1      ------------------------------------------------------------
```

### Supplementary Data S2

```
QHI97260.1      ------------------------------------------------------------
QHI97263.1      EDGIFQYYAADMHADILQMSEAVIALPQDLRGTVHAARVALRHLMDTYFTVATDAQRFSI
QHI97411.1      QAAPFTYLPADMQRDIGRMQAAVRSLPQALQGAEHAARAGMRYLMETYFTANTDAQRFSI
QHI97261.1      QSEPFSYSAADMQRDIAQMKAAVTNLPQDIRRAGHAARAAMRYLIEKYAKADTDPQRFSI
QHI97262.1      -----------------------RTISTPLRGSKSR------------------------

QHI97260.1      ------------------------------------------------------------
QHI97263.1      IKRLIQGRQAKPLVYILEQTVTSKFDEIEMLAAQLGGAKIGFEVAAGGSLDNWSYVFRYL
QHI97411.1      IKHLVQERQKNPLRYMLERTVAARFNVAEAVAARVGSARFPFDTSAAGLADGWSYIFLYL
QHI97261.1      IKRLVQERGKDALLYFLERTVASNFDLLDQLAAHPGGIKLQFSPFIGGASDNRHYIFAYL
QHI97262.1      ------------------------------------------------------------

QHI97260.1      ------------------------------------------------------------
QHI97263.1      QALLPLPEERRFLWDRLQDAFKLHRLQLADEINYAEGSIRLTYDSTIYPELAQFRFADLE
QHI97411.1      RALGHLPEERAFLWNTLQEAFRLHHMELKTEIDYQQGSVSVTYDNKAYPDFADFKIADLE
QHI97261.1      KKLAALPDECQFLFAALQDAFKLHNMKLQNNIDYAGNSVEVAIDNAVHQEFVDFKICDLE
QHI97262.1      ------------------------------------------------------------

QHI97260.1      ------------------------------------------------------------
QHI97263.1      EFVSRLPQGTSYRIGFKARQTAPGQLPRSVFDVDESKIDQGPYSKANKVLAYFNNVLSGP
QHI97411.1      EFVDGLPPGTNFGIEYALQPVDPDELPRSVFIVHGSRGPGDPHRKANDILLYLNHILPGP
QHI97261.1      EFIAGLPNNTDFDIEYSSYQPSPDRQPVSVFTVGNSAANDARYKQANEILLYLNNVIPGP
QHI97262.1      ------------------------------------------------------------

QHI97260.1      ---------------------------
QHI97263.1      QVEYVAVYFTRLLEMKNHAQGSGQALN
QHI97411.1      HIEILAHALIRQLDQKKGRQAPARAAT
QHI97261.1      SVENLARALKNQLESKKRKRN------
QHI97262.1      ---------------------------
```

### Supplementary Data S2

**HrpX-regulated protein family 4**

```
>QHI99309.1 hypothetical protein GT347_15795 [Xylophilus rhododendri]
MMPALANLAAPLPAAHPPPVEAQGCAVAEVAIEFEAYPPEIRLLVSEEWLRWPLADNPLLSQDLWNSRLQ
QTFHEPIAASMLMLALSLARDPTELRQALDALADAPHRYLGECWDQALAWFCDWIGRNHDPRPLVLLGDL
LDRLPPVQPLRLQLLHRLADLARRPRFGDRAGVADRLLKSCAELPDIPASLWRMLMQLQRRALKDPLRDR
HPIEQIPGVDRLPAARREALALLQRCGGRARHAQSAQALQLLIRQVEAVEDPAIRFELLELLRARPERGE
EPCAEFDLASVTQALLRVPGAAWHAKALRMLSLDELELTPRQLMQELGELAPVHAIRVLAMHIDTPVIDA
DDRQAMADGLIALLQRSREEPCGHLQFLHGMSYFIHRLGQPAVMQRLQELAMEDCTRVDPRYRLALLAVL
EPGARCGGDVRQLWKRHWLTALAQVSGTLRQARNAAQAWPAIQGLLPALRQEENRDAVLRLQIDALPLLP
AQDQARVLEQMLTHCLRSRFDTDEAHRVLLIEACRRLPFHLRAGPLRCLRRFCATAPQASVALLTALQRD
TTQAQAAWAENSLAVLASREPA

>QHI99316.1 hypothetical protein GT347_15830 [Xylophilus rhododendri]
MQVDDTRPQPARTVQLPGNPPLVAQGQDEADFEPTRFEELPPELALMLSDIWLKLHPAENPIGPIASGNA
HLHDVYRQPLRASDLIRALYCAQDDTQLRAAIDALADAPVQYQEDCWAHVWKILPRWVTPCSTWQLQALL
DHLLSRFPTAPALRLQQMQRAVRFVDACHSDAWSREFHDRLLRACLELPEIPRALWRLLLQLRHTSVADP
EAGPERIEDIDGAERLPAARRAELALLLRCMDQRYRQRTLAEMLDEIAEVETVADPAVRLDLLQWLRIRR
ATARTEDADTVAAARCRALLNIEGAAQREEVLRMLPLKEGELSVDTLRKELPGLPPVAAMRVLGAHFYRL
LEAAGGRRLLADCLVPLLLRSRETPSGHLAFLHLLALRVHLVCAVGLAPQLPDLLLEECARLEPCLRLAL
LDKLSTANWRSSETRQVWAREWRACLRQSIETLQHARTAAQAWPALRCLLVTLRRPELRDTVLQTVLSKL
PLLAAGDLALALKEVVLHCLFSPFLTSREQVTQLIAACAELPFHLRPGLLMQIRKLIAPHHEGEDEGLLA
LERQTAEAVERWRALPADRTDI

>QHI99310.1 hypothetical protein GT347_15800 [Xylophilus rhododendri]
MDNITLRNTTATTAGQPPDMTQPPPATRYEQMPDELMLLVVECSLRGAVQNPAQSGLPAFDCNQRQHRLL
TRPMAAYEVIKALWVAPNLPRFCQAADALAAVPPPYLDLCWQAAWPALAAMTRRLKARQAEAPLTHLLDR
LPDAAEMRLLQLRRAATFLLDHGHGAWRSDCADRILRLTAQGPGIPAPLWRQLLQLQHRAPHDASRRALA
DTPGLSAQQRGQWLVLQDCLDSRHHTPTLQHTQALLARIEAAGSRAIRFDLLCLLRQRLGVFEPAELKQA
NAALSRALLRLADSPLRPQVLQKVCLRDDPAIGQALPVQLAALAPRVGLQVLIRHIKAFHDSPAHLALLA
GCIGSLLARSRQAAQDHAAFLCTLAGLAEGVVDPELWQRLDRLLLDECAALTPAKRLRVLAALQVGIEGE
PLLRAEWQTAWDTALQDTVALLSQAGDGTQAGPLIQALLPALRRHESRDTVLGSQLAALRLLDAHAQAPL
LRQIVSMLLNGCYVVDDRHKAWLVQACSQLPFYLRATLLERLESLRSWPPHQSLEALAGLQGRTARQRRE
WAHGQPPG

>QHI99312.1 hypothetical protein GT347_15810 [Xylophilus rhododendri]
MKLNRMTPAGHSIKAFCRCNQVLRQTFAKPLAAHRAIKALWVAGGTEDFSKAIEALKEVPQQYEPECWDA
AWINLERLGRRKPTRDFERLFIHMLEQLPDTSGLRLQQLQRAAACLLKHSRDHRRYGKTQWPANQLMCLC
AAEPDLPDRLWRKVLHLQGKNLAHGGLFRPSAALVARLPQNRQQEFAILQRCREFFLETVSLQDALELLQ
RIEGVADPAIRFDLLQVATMCMMRREPDVWSVVSKKQREMLLTLASTHLRQQVLLSMWVDQNPALRRTLL
EQLKPLQPLAAAHVLDWQHFAFQNQPEAIAQLEQRLLELLVLSREQHADDHPKFLLALANCAGLIEDQTS
RQRLRTLVEGEIEQVGLRWRLPILAALETGTSCVIGVPPSWTERWNRALADTFAALRQAGSAEAAWPLVQ
ALLPALGKSGQRDAALDGLLAALPLLALDDQARVLCQIIERLEREPFCWLTPEHRCRLIEVCGGLPFYLR
SAPLEALAHDSDLPPQPGDALLAELQRETDEAMQAWAADESP

>QHI99313.1 hypothetical protein GT347_15815 [Xylophilus rhododendri]
MTLPSATQPPALPPGASFASTVAAAGQQIPPWFERLPPEIKLGTLEPFLLERPQASPVHAFSGCNREMAR
QLAPLRRADATVKALWTAPDFPEFRQAVETLGEIPLRYQSDCWHAAWSALARLSRMQNRPEVEAAFFHLF
DRLPADPAVLVQQLHRAVSWALHQGPGIWSSAGADAILRRCAALMPDMPDELWRRLLRLQRATRFTREAP
DYHELLATPGLSPAQRGQLELLRGCTKVGLEAMTEAQMLALIGRIEDVEDQAVRVELFRHLLPLLLAANP
PYRDVAKAAIDRALLRITDPAQRRQVLTRLRFRPGEAEPATLPAELAAMTPLQALRVLAWQVETLLSQTE
GSQLLDVCIDRLLERSCQDPQEHPLFLGRLAKLCVQFTDRVQRQRIETLVLHQCAQLPPWQRLPILEKLE
KEVWSCAEVEQAWSAAWEAAVAPATQALRQATTAQAAWPLLLGLLPALPKREGRDAVLRAQLDALPLLDP
RDQAQMLERMLRCCLRYGCQLEEGHTSLLIEACVRLPFYLRPPALDKLQNLCRRTLPGEASGPIQAQLAA
LLEQTAREQQAWMASAAAGATQA

>QHI99319.1 hypothetical protein GT347_15845 [Xylophilus rhododendri]
MHLVPVQQSKPLVAQNHEPGRFETLPNEIKYEVFRAMLQWIVPLPQCGGADLFGDLNQALAQTFAEPSAT
CRVIESLWAAGTGPGLRAALGAIEQAPAQYRCDCWRAAWAVLAAADRDENVDEPQVDAAFLHILDHLPAD
PGQRLRQLQHAARFMVDRIFDPRGTTSATRLLQHCAALPRIPDSLWRDLLLFWMHFSTAGDFQPGSSLEG
LTPVQQEELALMVECNRDVTDGPLAGRFPGWLQGAERVRDAAARLNLLLTLRARVRRVSRTQEECERNEK
AADEALLRVREPRLRPQVLRCIEAHADHPLRETLAQELAAFDPSEAIQVLLQQARALSLDTQGRALLTRC
LDELMQRSRLAPDRHADFLRNMAMASRNASGARMNEDIARRVMQECATLEPASRLTVLAALQITYWFDMD
MRQRWQADWRTALDETIAALPQADTSAAVLRLAQGLLPALRRPDSRDAALQAMLHGLPAMEPHLQARLLQ
KIVGCCIQQDPMLLEQHALQLIQACARLPFHLRPARLDLLPCQPEAPSCRQLADLVRQTRQEHLNWAIQS
RTRQATGAGAR

>QHI99317.1 hypothetical protein GT347_15835 [Xylophilus rhododendri]
MHITSASQGSGSIHPPQPLSFEALPNELKQTIADDILNRLLWQPEAGGPEVFVGNAALAAQLARQVAASA
AIKALWVAADVPSFRRAVDAIDSAPQEYKSQCWQAAWHALMAFASANAKAPVVEAMLAHMLEHLPENPAL
```

RLRQLRRATAFLQRKAGVGWSLSSATTILRRCAEYPDLSPGLWHRLLELQLLHMSYHPSAYRHWRIAMTA
GLAAAQFSQLTLLGDCVRRDPVEPTPRALRGLLARIERIADPAIRLTLFGVLESRAMWQRRLRPEFADVQ
REVHQAMLKLTDPALRPRVLEIVPFQEGHAERDTLLQELAAMPPLAAMAVLDRQFLNFAPDAEGMDLLRQ
CVDALLERSRNAAETHPALLHKLAIASRQLGDEALRLHIAARVLQESAALGAADRLTVLTIRHTASWPGD
ALRPDWESAQRAALAATSDLLRQAKAACAARPLMLAMTNALGLQDSRDAVLQAQRAALSLFEPERQAELL
DDIAFQCVVDPLAMEEEHTLQLIEDCGRLPFYLRPRLLDTLGYLCSDPESPGATQLAALQQRTRQERQAW
IDLPPPGGIARPSSHGEPPCR

>QHI99314.1 hypothetical protein GT347_15820 [Xylophilus rhododendri]
MRLDLLGLLYMVVLMGNPHSEERESAETQVHQARLRITDPALRKEVLLEIPFEEEIAPRVDLLRELLAFP
PVEAIRLLEPQVMTLGVDTHGLLLTGQSMELLLARSRQVPDGHPAFLHTLGRASKSIPDVVLRTRIGGLV
LQDSLRLEVGHRLNILAALDGPEAGFRERWRAEWAAALAGTLQALGQATGSASAWHLIHALVPASKKRQS
CDAVLQAQLAALPLLDPHDQVRALHTICHEYLVDRFDKEDGHLLQFIEAARQLPCYLRPRPGKLLHLLSK
EAAAAVKARLGALDEETQAQRQAGLHDRGATAAPAPKPVG

>QHI99318.1 hypothetical protein GT347_15840 [Xylophilus rhododendri]
MHVTATTTTTTTTVASQTASTSASAPPLRYEELPEEMLLEVAHWSLRWMLDHPQADGVTPLADCNQRLGR
ALAEPAAAAAAISALWASKSVAQARMAIDALEHVSPHHTAECWEAAWRALKTRALRPPPRPDVEILFFHM
LDRLPEDPQQRLRQMRRAVAFSYKQSRFTSEWRLAPIAKLLRHCAQLPVLPDSMWRCLLKMLADKLCAEN
LETGRQLAWTAGLSPQRQQQLTLLLDCLDGLPGPMTPQRMRALLGPVEQIADPAIRLTLLWCLQRRAVAC
AEPSEALNAAEAELHRALLRIRDAALRPKVLENLPFLQYLVPQNTLLAELAALTPLAALEVIASQTHAFG
RDSDGIALLGRCLDETMQRSRELPDGHPAFLCAMGKDLRHLRDQALHARIAGLVLEDSRALGAAQRLDVL
AALDRLEWRDIDLQDRWQAEWTAAVAAAHHALRQATTGAAAWPLLLALVPALTKPDQHDAALQAQLEALP
LLHPHDAARHLHTICIECMLSPVNKAGMQEDQLIQLIEAGRCLPSYLRRDSIPLLNFMLLPGDEAAVSQA
RMKALAEETEQEHRAWVALHAPGLALTARATDDTGD

>QHI99311.1 hypothetical protein GT347_15805 [Xylophilus rhododendri]
MIPPPGAASQLPASGAGLPVPPPPDFEGLVPELPGMVAREIQHGNPDTPPLDTVPVIAGLNSRLDAIFSG
PAAASKVIGRLWLASDGDAFKAAVDTIAEVPEACIGPCWDAVWTALQRLDPAETERSVEHLLEHLPSHPQ
LRALQWERAIVYLTAHGCSMRNAVADRLLQIRAAMGAIPDDTWHKLLRLYGRMRLHRLDPDPPALVLPSL
TPGQQADLPLLAAMRRTSTLGRRQHFLELLERARRLADPRIRLLMLQLLLQRRCHLNRGEQDLTTTQLPE
AFLEFGNTPFARQALKVMPSPEGDPQRRLQLQQQLLQQLQAFSPLAAMDILAHHAWSFHQGGEDLRLLEK
TWLGLLDRSLQYPHEHLHLLRRLVNNRFFLTAAICQRFLAECAAVPLQQRLALLAPLDWQAERRPDFQPA
WKALWDSTRQDIAAALDRSTRAPDIRIWLKALLPALRRHPASLLRRLPAALAVFEPPEQAQILASFCQGL
RPARVLKKVPETGLLLLIELCARLPFYLRREALKGLGALPRSTAPVAARLAQLRRRTEAEIQAWAADSAA
A

>QHI99315.1 hypothetical protein GT347_15825 [Xylophilus rhododendri]
MHFPPTAPGPITKPRTLPPPRYEQQAPEMKSAIARCNLQWQLFEPESGGIACFAQCNSTLARELALEVAA
SAVIHALGTANDAAQFQAALDTVASAPACYLPECWKAAWNALEALSAPAKAAELQAMFTAVLDRLPAHPG
LRLQQLRRAARFGSQCSRGTSAWSVESTARLLQLWAEQADMPPRVWRQLLQMQQRTQYRQVATDFERILR
TPGLTAEQQDQLQLQMHCMCRRLPPRSCRRCSPRWSVSPTRACAWTCWDCCTWSC

>QHI99324.1 hypothetical protein GT347_15875 [Xylophilus rhododendri]
MNTASVQDTRHTPPASRTAWQTTAGTEPGWLERLPVELQYEILDKLLKLGNEEAGINPFKGLNSRLDKSL
ASPFGASELIRALWQAPDAAGFLAEVDAIDEMPQQYHEACWNAVWNALARFDLRDADADLAGRLLDRLPA
HPVLRRQQWRCALDLVTRGRQPCTLRFVERLMQLGAAGGEAAPGTWRRLLQLYHLAGVRSSVEARGPVAG
LPPAWSAQLALLQECMRLGDVDHDATLPEFHALLGRILAVADPDIRLLLLANAQSCLFGLPEQDQEIACL
ALRDAAAPFIGTPLEARLLQTVFRAFWNDDQGVVLKAVQKRTPEDALRLIESLEVGHGPLLDPMLRQALP
ALLRQCVDRVREHPQLLLRLAGICEWCGQDPGRDIGRALLAHCAQLDLHLRLAFLEKLAIHSQQDPDGQA
RWQAQWQQALQAICTALAAAETPAARRRLAKALLPALRFTHSLETALPPLLAAAPKLLPDERLDVLTAAS
MDVLAHTDIDEDTWWTRLAVLRRAAYGDLPLQLQSGLLKTQNTMLALFLEQKNADL

>QHI98760.1 hypothetical protein GT347_12630 [Xylophilus rhododendri]
MLSRVPPPSSPPRAPLPKVGTALIVAPQRRLEYLPNELLQELFLTGMEQNAVWGCLGIDATRVVAGLNAR
MAAALAQAAPASEAMRQLTTARDTPAFAAALALLPRVAPQYRQRCREAAWWAVVHRAPAPVQRSQEDWLH
CLLDAEASSGKLHPPLWQRAVRLLARPEMDAPGSALVGRLLACAAAARTMAPADWRRLLRLVARSMEGRG
PDASCFDAKGLNTPQQAELDTVLACCALSWQRCDRTTVRAMIARIEASVSPDSRCEVLVALSRSWPWNLS
LDAVALEPLRDALLRVPDGDLERALEELPPIGDPASRQRMFLDQCARLPPDSALRVLDRQSYDFDRDTQG
RDLLQAGSIALLNAALAQPAIRCQALQAFARLSARLKPGADTSGLSTLALRACGNLPAAERIAVLARLKP
DAPNAQPWAAAWQAALAELADDETPGKPWRDVRALTSGLWSRCGRAAVLPRLEAALQGLTVQDRSRALAE
LAALWNRCPGGCTEAEFDGLLRWCEALPFYLRQPPLQVLRQTAATLPPYCAHELEALFTASDEAHAAWAA
GLPPLS

### Supplementary Data S2

```
QHI98760.1      ------MLSRVPPPSSPPRAPLPKVGTALIVAP----QRRLEYLPNELLQELFLTGMEQN
QHI99324.1      --------MNTASVQDTRHTPPASRTAWQTTAGTE--PGWLERLPVELQYEIL----DKL
QHI99311.1      MIPPPGAASQLPASGAGLPVPP---------------PPDFEGLVPELPGMVA-REIQHG
QHI99316.1      ----MQVDDTRPQPARTVQLPGNPPLVAQGQDEADFEPTRFEELPPELALMLS----DIW
QHI99309.1      ------MMPALANLAAPLPAAHPPPVEAQGCAVAEV-AIEFEAYPPEIRLLVS----EEW
QHI99315.1      --------MHFPPTAPGPITKPRTLP-----------PPRYEQQAPEMKSAIA-RCNLQW
QHI99312.1      --------MKLNRMT---------------------------------------------
QHI99319.1      --------MHLVPVQQSKPLVAQNHE-----------PGRFETLPNEIKYEVF-RAMLQW
QHI99310.1      MDNITLRNTTATTAGQPPDMTQPPP------------ATRYEQMPDELMLLVV-ECSLRG
QHI99317.1      --------MHITSASQGSGSIHPPQ------------PLSFEALPNELKQTIA-DDILNR
QHI99313.1      --------MTLPSATQPPALPPGASFASTVAAAGQQIPPWFERLPPEIKLGTL----EPF
QHI99314.1      ------------------------------------------------------------
QHI99318.1      --------MHVTATTTTTTTTVASQTASTSASAP---PLRYEELPEEMLLEVA-HWSLRW


QHI98760.1      AVWGCLGIDATRVVAGLNARMAAALAQAAPASEAMRQLTTARDTPAFAAALALLPRVAPQ
QHI99324.1      LKLG-NEEAGINPFKGLNSRLDKSLASPFGASELIRALWQAPDAAGFLAEVDAIDEMPQQ
QHI99311.1      NPDT-PPLDTVPVIAGLNSRLDAIFSGPAAASKVIGRLWLASDGDAFKAAVDTIAEVPEA
QHI99316.1      LKLH-PAENPIGPIASGNAHLHDVYRQPLRASDLIRALYCAQDDTQLRAAIDALADAPVQ
QHI99309.1      LRWP-LADNPLLSQDLWNSRLQQTFHEPIAASMLMLALSLARDPTELRQALDALADAPHR
QHI99315.1      QLFE-PESGGIACFAQCNSTLARELALEVAASAVIHALGTANDAAQFQAALDTVASAPAC
QHI99312.1      -----PAGHSIKAFCRCNQVLRQTFAKPLAAHRAIKALWVAGGTEDFSKAIEALKEVPQQ
QHI99319.1      IVPL-PQCGGADLFGDLNQALAQTFAEPSATCRVIESLWAAGTGPGLRAALGAIEQAPAQ
QHI99310.1      AVQN-PAQSGLPAF-DCNQRQHRLLTRPMAAYEVIKALWVAPNLPRFCQAADALAAVPPP
QHI99317.1      LLWQ-PEAGGPEVFVG-NAALAAQLARQVAASAAIKALWVAADVPSFRRAVDAIDSAPQE
QHI99313.1      LLER-PQASPVHAFSGCNREMARQLAPLRRADATVKALWTAPDFPEFRQAVETLGEIPLR
QHI99314.1      ------------------------------------------------------------
QHI99318.1      MLDH-PQADGVTPLADCNQRLGRALAEPAAAAAAISALWASKSVAQARMAIDALEHVSPH


QHI98760.1      YRQRCREAAWWAVVH--RAPAPVQRSQEDWLHCLLDAEASSGKLHPPLWQRAVRLLARPE
QHI99324.1      YHEACWNAVWNALAR--FDLRDADADLA---GRLLDRLPAHPVLRRQQWRCALDLVTRGR
QHI99311.1      CIGPCWDAVWTALQR--LDPAETERSVE----HLLEHLPSHPQLRALQWERAIVYLTAHG
QHI99316.1      YQEDCWAHVWKILPR--WVTPCSTWQLQALLDHLLSRFPTAPALRLQQMQRAVRFVDACH
QHI99309.1      YLGECWDQALAWFCD--WIGRNHDPRPLVLLGDLLDRLPPVQPLRLQLLHRLADLARRPR
QHI99315.1      YLPECWKAAWNALEA--LSAPAKAAELQAMFTAVLDRLPAHPGLRLQQLRRAARFGSQCS
QHI99312.1      YEPECWDAAWINLER--LGRRKPTRDFERLFIHMLEQLPDTSGLRLQQLQRAAACLLKHS
QHI99319.1      YRCDCWRAAWAVLAAADRDENVDEPQVDAAFLHILDHLPADPGQRLRQLQHAARFMVDRI
QHI99310.1      YLDLCWQAAWPALAA--MTRRLKARQAEAPLTHLLDRLPDAAEMRLLQLRRAATFLLDHG
QHI99317.1      YKSQCWQAAWHALMAF-ASANAKAPVVEAMLAHMLEHLPENPALRLRQLRRATAFLQRKA
QHI99313.1      YQSDCWHAAWSALAR--LSRMQNRPEVEAAFFHLFDRLPADPAVLVQQLHRAVSWALHQG
QHI99314.1      ------------------------------------------------------------
QHI99318.1      HTAECWEAAWRALKTR-ALRPPPRPDVEILFFHMLDRLPEDPQQRLRQMRRAVAFSYKQS


QHI98760.1      MDA--PGSA--LVGRLLACAAAA-RTMAPADWRRLLRLVARSM----EGRGPDASCFDAK
QHI99324.1      QP---CTLR--FVERLMQLGAAG-GEAAPGTWRRLLQLY-----HLAGVRSSVEARGPVA
QHI99311.1      -----CSMRNAVADRLLQIRAAM-GAIPDDTWHKLLRLYGR---MRLHRLDPDPPALVLP
QHI99316.1      SDA--WSRE--FHDRLLRACLEL-PEIPRALWRLLLQLRHTSVADPEAGPERIEDIDGAE
QHI99309.1      -----FGDRAGVADRLLKSCAEL-PDIPASLWRMLMQLQRRALKDPLRDRHPIEQIPGVD
QHI99315.1      RGTSAWSVE--STARLLQLWAEQ-ADMPPRVWRQLLQMQQRTQ-YRQVATDFERILRT-P
QHI99312.1      RDHRRYGKTQWPANQLMCLCAAE-PDLPDRLWRKVLHLQGK---NLAHGGLFRPSAALVA
QHI99319.1      FDP--RGTT--SATRLLQHCAAL-PRIPDSLWRDLLLFWM----HFSTAGDFQPGSSL-E
QHI99310.1      HGA--WRSD--CADRILRLTAQG-PGIPAPLWRQLLQLQ-----HRAPHDASRRALADTP
QHI99317.1      GVG--WSLS--SATTILRRCAEY-PDLSPGLWHRLLELQLLHM-SYHPSAYRHWRIAMTA
QHI99313.1      PGI--WSSA--GADAILRRCAALMPDMPDELWRRLLRLQRATR-FTREAPDYHELLAT-P
QHI99314.1      ------------------------------------------------------------
QHI99318.1      RFTSEWRLA--PIAKLLRHCAQL-PVLPDSMWRCLLKMLADK--LCAENLETGRQLAWTA
```

**Supplementary Data S2**

```
QHI98760.1      GLNTPQQAELDTVLACCALS--WQRCDRTTVRAMIARIEASVSPDSRCEVLVALSRSWPW
QHI99324.1      GLPPAWSAQLALLQECMRLGDVDHDATLPEFHALLGRILAVADPDIRLLLLA--NAQSCL
QHI99311.1      SLTPGQQADLPLLAAMRRTS---TLGRRQHFLELLERARRLADPRIRLLMLQLLLQRRCH
QHI99316.1      RLPAARRAELALLLRCMDQR--YRQRTLAEMLDEIAEVETVADPAVRLDLLQWLRIRRAT
QHI99309.1      RLPAARREALALLQRCGGRA--RHAQSAQALQLLIRQVEAVEDPAIRFELLELLRARPER
QHI99315.1      GLTAEQQDQLQLQMHCM-------------------------------------------
QHI99312.1      RLPQNRQQEFAILQRCREFF--LETVSLQDALELLQRIEGVADPAIRFDLLQV--ATMCM
QHI99319.1      GLTPVQQEELALMVECNRDV--TDGPLAGRFPGWLQGAERVRDAAARLNLLLTLRARVRR
QHI99310.1      GLSAQQRGQWLVLQDCLDSR--HHTPTLQHTQALLARIEAAGSRAIRFDLLCLLRQRLGV
QHI99317.1      GLAAAQFSQLTLLGDCVRRD--PVEPTPRALRGLLARIERIADPAIRLTLFGVLESRAMW
QHI99313.1      GLSPAQRGQLELLRGCTKVG--LEAMTEAQMLALIGRIEDVEDQAVRVELFRHLLPLLLA
QHI99314.1      ------------------------------------------------MRLDLLGLLYMVVLM
QHI99318.1      GLSPQRQQQLTLLLDCLDGL--PGPMTPQRMRALLGPVEQIADPAIRLTLLWCLQRRAVA

QHI98760.1      NLSLD---AVALEPLRDALLRVPDGDLERALEELPPIGDPASRQR-----MFLDQCARLP
QHI99324.1      FGLPEQDQEIACLALRDAAAPFIGTPLEARLLQTVFRAFWNDDQG-----VVLKAVQKRT
QHI99311.1      LNR--GEQDLTTTQLPEAFLEFGNTPFARQALKVMPSPEGDPQRRLQLQQQLLQQLQAFS
QHI99316.1      ART--EDADTVAAARCRALLNIEGAAQREEVLRMLPLKEGELSVD-----TLRKELPGLP
QHI99309.1      GEEPCAEFDLAS--VTQALLRVPGAAWHAKALRMLSLDELELTPR-----QLMQELGELA
QHI99315.1      ------------------------------------------------------------
QHI99312.1      MRREPDVWSVVSKKQREMLLTLASTHLRQQVLLSMWVDQNPALRR-----TLLEQLKPLQ
QHI99319.1      VSRTQEECERNEKAADEALLRVREPRLRPQVLRCIEAHADHPLRE-----TLAQELAAFD
QHI99310.1      FEP--AELKQANAALSRALLRLADSPLRPQVLQKVCLRDDPAIGQ-----ALPVQLAALA
QHI99317.1      QRRLRPEFADVQREVHQAMLKLTDPALRPRVLEIVPFQEGHAERD-----TLLQELAAMP
QHI99313.1      ANPPY--RDVAKAAIDRALLRITDPAQRRQVLTRLRFRPGEAEPA-----TLPAELAAMT
QHI99314.1      GNPHSEERESAETQVHQARLRITDPALRKEVLLEIPFEEEIAPRV-----DLLRELLAFP
QHI99318.1      CAEPSEALNAAEAELHRALLRIRDAALRPKVLENLPFLQYLVPQN-----TLLAELAALT

QHI98760.1      PDSALRVLDRQSYDFDRDTQGRDLLQAGSIALLNAALAQ-PAIRCQALQAFARLSARLKP
QHI99324.1      PEDALRLI--ESLEVGHGPLLDPMLRQALPALLRQCVDR-VREHPQLLLRLAGICE-WCG
QHI99311.1      PLAAMDILAHHAWSFHQGGEDLRLLEKTWLGLLDRSLQY-PHEHLHLLRRLVNNRFFLTA
QHI99316.1      PVAAMRVLGAHFYRLLEAAGGRRLLADCLVPLLLRSRET-PSGHLAFLHLLALRVHLVCA
QHI99309.1      PVHAIRVLAMHIDTPVIDADDRQAMADGLIALLQRSREE-PCGHLQFLHGMSYFIHRLGQ
QHI99315.1      ------------------------------------------------------------
QHI99312.1      PLAAAHVLDWQHFAFQNQPEAIAQLEQRLLELLVLSREQHADDHPKFLLALANCAGLIED
QHI99319.1      PSEAIQVLLQQARALSLDTQGRALLTRCLDELMQRSRLA-PDRHADFLRNMAMASRNASG
QHI99310.1      PRVGLQVLIRHIKAFHDSPAHLALLAGCIGSLLARSRQA-AQDHAAFLCTLAGLAEGVVD
QHI99317.1      PLAAMAVLDRQFLNFAPDAEGMDLLRQCVDALLERSRNA-AETHPALLHKLAIASRQLGD
QHI99313.1      PLQALRVLAWQVETLLSQTEGSQLLDVCIDRLLERSCQD-PQEHPLFLGRLAKLCVQFTD
QHI99314.1      PVEAIRLLEPQVMTLGVDTHGLLLTGQSMELLLARSRQV-PDGHPAFLHTLGRASKSIPD
QHI99318.1      PLAALEVIASQTHAFGRDSDGIALLGRCLDETMQRSREL-PDGHPAFLCAMGKDLRHLRD

QHI98760.1      GADTSGLSTLALRACGNLPAAERIAVLARLK----PDAPNAQPWAAAWQAALAE----LA
QHI99324.1      QDPGRDIGRALLAHCAQLDLHLRLAFLEKLAIHSQQDPDGQARWQAQWQQALQAICTALA
QHI99311.1      A-----ICQRFLAECAAVPLQQRLALLAPLDWQAERRPDFQPAWKALWDSTRQDIAAALD
QHI99316.1      VGLAPQLPDLLLEECARLEPCLRLALLDKLSTANWRSSETRQVWAREWRACLRQSIETLQ
QHI99309.1      PAVMQRLQELAMEDCTRVDPRYRLALLAVLEPGARCGGDVRQLWKRHWLTALAQVSGTLR
QHI99315.1      --------------CRRLPPR--------------SCRRCSPRWSVSPTRACA-------
QHI99312.1      QTSRQRLRTLVEGEIEQVGLRWRLPILAALETGTSCVIGVPPSWTERWNRALADTFAALR
QHI99319.1      ARMNEDIARRVMQECATLEPASRLTVLAALQITYWFDMDMRQRWQADWRTALDETIAALP
QHI99310.1      PELWQRLDRLLLDECAALTPAKRLRVLAALQVGIEGEPLLRAEWQTAWDTALQDTVALLS
QHI99317.1      EALRLHIAARVLQESAALGAADRLTVLTIRHTASWPGDALRPDWESAQRAALAATSDLLR
QHI99313.1      RVQRQRIETLVLHQCAQLPPWQRLPILEKLEKEVWSCAEVEQAWSAAWEAAVAPATQALR
QHI99314.1      VVLRTRIGGLVLQDSLRLEVGHRLNILAALD---GPEAGFRERWRAEWAAALAGTLQALG
QHI99318.1      QALHARIAGLVLEDSRALGAAQRLDVLAALDRLEWRDIDLQDRWQAEWTAAVAAAHHALR
                                 :                         *
```

### Supplementary Data S2

```
QHI98760.1      DDETPGKPWRDVRALTSGLWSRCGRAAVLPRLEAALQGLTVQDRSRALA----ELAALWN
QHI99324.1      AAETPAARRRLAKALLPALRFTHSLETALPPLLAAAPKLLPDERLDVLTAASMDVLAHTD
QHI99311.1      RSTRAPDIRIWLKALLPALRRHPA--SLLRRLPAALAVFEPPEQAQILA----SFCQGLR
QHI99316.1      HARTAAQAWPALRCLLVTLRRPELRDTVLQTVLSKLPLLAAGDLALALK----EVVLHCL
QHI99309.1      QARNAAQAWPAIQGLLPALRQEENRDAVLRLQIDALPLLPAQDQARVLE----QMLTHCL
QHI99315.1      --------W---------------------------------------------TCWDC-
QHI99312.1      QAGSAEAAWPLVQALLPALGKSGQRDAALDGLLAALPLLALDDQARVLC----QIIERLE
QHI99319.1      QADTSAAVLRLAQGLLPALRRPDSRDAALQAMLHGLPAMEPHLQARLLQ----KIVGCCI
QHI99310.1      QAGDGTQAGPLIQALLPALRRHESRDTVLGSQLAALRLLDAHAQAPLLR----QIVSMLL
QHI99317.1      QAKAACAARPLMLAMTNALGLQDSRDAVLQAQRAALSLFEPERQAELLD----DIAFQCV
QHI99313.1      QATTAQAAWPLLLGLLPALPKREGRDAVLRAQLDALPLLDPRDQAQMLE----RMLRCCL
QHI99314.1      QATGSASAWHLIHALVPASKKRQSCDAVLQAQLAALPLLDPHDQVRALH----TICHEYL
QHI99318.1      QATTGAAAWPLLLALVPALTKPDQHDAALQAQLEALPLLHPHDAARHLH----TICIECM

QHI98760.1      RCPGGC---TEAEFDGLLRWCEALPFYLRQPPLQVLR-----QTAATLPPYCAHELEALF
QHI99324.1      IDEDTW---WTRLAVLRRAAYGDLPLQLQSGLLKTQN----------------TMLALFL
QHI99311.1      PARVLKK-VPETGLLLLIELCARLPFYLRREALKGLG-----ALPR-STAPVAARLAQLR
QHI99316.1      FSPFLT---SREQVTQLIAACAELPFHLRPGLLMQIR-----KLIAPHHEGEDEGLLALE
QHI99309.1      RSRFDT---DEAHRVLLIEACRRLPFHLRAGPLRCLR-----RFCATAPQASVALLTALQ
QHI99315.1      ---CTW-------------SC---------------------------------------
QHI99312.1      REPFCW--LTPEHRCRLIEVCGGLPFYLRSAPLEALA-----HDSDLPPQPGDALLAELQ
QHI99319.1      QQDPML---LEQHALQLIQACARLPFHLRPARLD--------LLPCQPEAPSCRQLADLV
QHI99310.1      NGCYVV---DDRHKAWLVQACSQLPFYLRATLLERLE-----SLRSWPPHQSLEALAGLQ
QHI99317.1      VDPLAM---EEEHTLQLIEDCGRLPFYLRPRLLDTLG-----YLCSDPESPGATQLAALQ
QHI99313.1      RYGCQL---EEGHTSLLIEACVRLPFYLRPPALDKLQNLCRRTLPGEASGPIQAQLAALL
QHI99314.1      VDRFDK---EDGHLLQFIEAARQLPCYLRPRPGKLLH-----LLSKEAAAAVKARLGALD
QHI99318.1      LSPVNKAGMQEDQLIQLIEAGRCLPSYLRRDSIPLLNFM---LLPGDEAAVSQARMKALA

QHI98760.1      TASDEAHAAWAAGLPPLS-------------
QHI99324.1      EQKNADL------------------------
QHI99311.1      RRTEAEIQAWAADSAAA--------------
QHI99316.1      RQTAEAVERWRALPADRTDI-----------
QHI99309.1      RDTTQAQAAWAENSLAVLASREPA-------
QHI99315.1      -------------------------------
QHI99312.1      RETDEAMQAWAADESP---------------
QHI99319.1      RQTRQEHLNWAIQSRTRQATGAGAR------
QHI99310.1      GRTARQRREWAHGQPPG--------------
QHI99317.1      QRTRQERQAWIDLPPPGGIARPSSHGEPPCR
QHI99313.1      EQTAREQQAWMASAAAGATQA----------
QHI99314.1      EETQAQRQAGLHDRGATAAPAPKPVG-----
QHI99318.1      EETEQEHRAWVALHAPGLALTARATDDTGD-
```

### Supplementary Data S2

**HrpX-regulated protein family 5**

```
>QHI99321.1 hypothetical protein GT347_15855 [Xylophilus rhododendri]
MDISRTPPAQPSWGALSAGAVDPSKPPSGGEPTRLEHLSNELLVEIFDHLDPGFRYPGRRRGEEARGIAV
FTGLNRRFAEVGRHHAVAEDLARRARCPGTARQCADVVKEALALPVSLRLRVLHTLVEAVQFADGQAHGA
LAREAARCIVEEHPGLLRQDLFLKLQIHAFARKDICCPYLDRLHETAPLSPRQEQMLQTRFHSARVRTLS
EHLAHVEQKKGQDLLMHSLAGVCTQPAADVVATIHRRRIDAWFAGLEPARQATMLRALAAAIRSSSVRHR
PAHFEALLLLYGHLAQADRGRHTGLMEATARHVPGGFAQLAALTAAGMVLPPPSASSRVPPDILATPPVV
PPDSEPEDGVQVVTRTRLEQLPNEVLQWLFDYVDPCLADVAPDDKGARGVAAFTSLNRRLAHAGRDHAAA
ARILAAARRLATPQHCQALVSPLLQLPAALQSRVLHALMELMRQHNGPVHWAMRECVVLLLLEHLGAAVR
GSEFIRVQLHELVLNGASARLEKAVRAEVPLTSREEHLVRTRFFRSRLGMLGGYLSLTRHREPDAWASAI
AGACSQPDTSEHELLARGRIGELFEQMDDTQRRAMSRELLRAIRTSLQINRELPFLMLLRLHEKMPDAER
RWHVHRLAAAALHVPEGPELLAAARPG

>QHI99322.1 hypothetical protein GT347_15860 [Xylophilus rhododendri]
MRRDPSTHRVSFAQPVVRKGGRAGRLDFSTMREERSFLSVRCFTGKFEIVAALLPRIKRLEPCAWILTIA
GTCARCDADMQALSSGLRLDALFSGMDAAEQWAMSRALRRSIRNSLQRQRAASFEALLLLHEKMPAADRP
RHEQRLSATACHVNGGAALLAWHRARRMADTVAAAASPPTETETPPWPIPSWPHRQPSPSPRSPPPTA
```

```
QHI99321.1      MDISRTPPAQPSWGALSAGAVDPSKPPSGGEPTRLEHLSNELLVEIFDHLDPGFRYPGRR
QHI99322.1      ----------------------------------------------------------MR
                                                           *

QHI99321.1      RGEEARGIAVFTGLNRRFAEVGRHHAVAEDLARRARCPGTARQCADVVKEALALPVSLRL
QHI99322.1      RDPSTHRVSFAQPVVRKGGRAGRLD-----------------------------------
                *. .:. ::.   : *. . .**

QHI99321.1      RVLHTLVEAVQFADGQAHGALAREAARCIVEEHPGLLRQDLFLKLQIHAFARK-DICCPY
QHI99322.1      --FSTMREERSF--------------------------------LSVRCFTGKFEIVAAL
                  : *: *  .*                                *.:..*: * :* ..

QHI99321.1      LDRLHETAPLSPRQEQMLQTRFHSARVRTLSEHLAHVEQKKGQDLLMHSLAGVCTQPAAD
QHI99322.1      LPRIKRLEPCA----------------------------------WILTIAGTCARCDAD
                * *::   * :                                   : ::**.*:.  **

QHI99321.1      VVATIHRRRIDAWFAGLEPARQATMLRALAAAIRSSSVRHRPAHFEALLLLYGHLAQADR
QHI99322.1      MQALSSGLRLDALFSGMDAAEQWAMSRALRRSIRNSLQRQRAASFEALLLLHEKMPAADR
                : *     *:** *:*::.* * :* ***  :**.*  *:*.* *******: ::. ***

QHI99321.1      GRHTGLMEATARHVPGGFAQLAALTAAGMVLPPPSASSRVPPDILATPPVVPPDSEPEDG
QHI99322.1      PRHEQRLSATACHVNGGAALLAWHRARRMADTVAAAAS--PPTETETPP-----------
                 **   :.*** ** ** * **   *  *. . .:*:*  **    ***

QHI99321.1      VQVVTRTRLEQLPNEVLQWLFDYVDPCLADVAPDDKGARGVAAFTSLNRRLAHAGRDHAA
QHI99322.1      ------------------W-----------------------------------------
                                  *

QHI99321.1      AARILAAARRLATPQHCQALVSPLLQLPAALQSRVLHALMELMRQHNGPVHWAMRECVVL
QHI99322.1      ----------------------PIPSWP--------------------------------
                                      *: . *

QHI99321.1      LLLEHLGAAVRGSEFIRVQLHELVLNGASARLEKAVRAEVPLTSREEHLVRTRFFRSRLG
QHI99322.1      ------------------------------------------------------------


QHI99321.1      MLGGYLSLTRHREPDAWASAIAGACSQPDTSEHELLARGRIGELFEQMDDTQRRAMSREL
QHI99322.1      ----------HRQPS------PSPRSPPPTA-----------------------------
                          **:*.      ... * * *:

QHI99321.1      LRAIRTSLQINRELPFLMLLRLHEKMPDAERRWHVHRLAAAALHVPEGPELLAAARPG
QHI99322.1      ----------------------------------------------------------
```

### Supplementary Data S2

**HrpX-regulated protein family 6**

```
>QHI99589.1 hypothetical protein GT347_17385 [Xylophilus rhododendri]
MPFDIPAPDCTDACFSSPAASAATRFADLVQALLEAAPHETSDLSGLHAALAAVTGDTDPASRPDRTEET
RYHLDKRNPAGKYYWCDPGIGPFRVAPLLDVLKAIALARRRGPLSNEDRELLARLAEMLPEALRVECGRD
SRLDAIFHPPEPVAVPAVLDSSAREQLRADLASGRLASPQALADALVGAAHRYVIAELTKAHPDKAVPIG
NFSLRDINRRQIVPVAQTDEFFTQQQPFIVDSLHAALMRIADELGHAGPQATEASASIDSTISTGLILFN
QLMELQAAPGWRMHAETLRKLATRQPSTVDEMRDILSRSMAARMLKEAKFGYARGTQWEEEKTLHFFWGA
ERLLLRPESRMVLGRVLQDQLIAAKEQHARAVARRQKLDPALTQALVDSAASHVEHYFGFMSSEARPFVR
PPEPAAPPAAARPDPLQRYRRAIDELDEGTDIDDPQDPRFVSGGDFLARSNALLAVSRAEQQADAAGEPF
DHQAFATASIDPDVIASAGYLKTLFDRWAA
```

```
>QHI99590.1 hypothetical protein GT347_17390 [Xylophilus rhododendri]
MFSSHFHQFLSNLSQCLGRKAPQPQAAPGPQPPGLPAATGGIGKTLQEAVPKIAADAASFSQPDPQQAPL
PASSAADRLHDLAGALVSAEHASTACLQQLFAKSDLSTLLGILPPAALLDVLTFAFDPAFECPEGRAFDS
LKNWMLETSGHDGRLVGQVLKQAVWRKALQHLPVSDAAMAQRVDAIQSQLDASGDYQAYLAANLEIVREL
REKSGAAGSAAEPQALFLNNDIEKWRAALLDGSIDSPEKLTAVLLDAVDRNYAVRLARLYPGAAMKIGKY
GLGGVKHGYVSSTELSEFTQENLQATLAAAHTAMLEMLNDPAFAPSAGGPEIGGILSDGLLLLQNILEIP
QRHQANDYVLREIAQTQPSTPEHIGEILEQSMMQRQVTLKKFGHTSVEERTSVDFFWTALKLLRKPASKM
ALGKAIQDLIIQFKQVQAQAIGKRQQLPPDGIEALLDNTVSHVRHYYDFMAQAADPFATPAGMAPFSETD
APYRQAVEDLENGTRIADPKDPRFVSASDFLLRSHALLEVNRAERAARIAGTPFDRAAFIDSSTNADVAA
AAPYLQASFSRWPA
```

```
QHI99589.1      ----------------------MPFDIPAPD-----------------------CTDACF
QHI99590.1      MFSSHFHQFLSNLSQCLGRKAPQPQAAPGPQPPGLPAATGGIGKTLQEAVPKIAADAASF
                                       *   *.*:                       .  *.*

QHI99589.1      SS--------PAASAATRFADLVQALLEAAPHETSDLSGLHA--ALAAVTGDTDPASRPD
QHI99590.1      SQPDPQQAPLPASSAADRLHDLAGALVSAEHASTACLQQLFAKSDLSTLLGILPPAALLD
                *.        **:*** *: **. **:.*   .*: *. *.*   *::: *   **:  *

QHI99589.1      RTEETRYHLDKRNPAGKYY-----WCDPGIGPFRVAPLL-DVLKAIALARRRGPLSNEDR
QHI99590.1      -VLTFAFDPAFECPEGRAFDSLKNWMLETSG--HDGRLVGQVLKQAVWRKALQHLPVSDA
                 .    :      * *. :     *     *  . . *: :***  .  .    *. .*

QHI99589.1      ELLARLAEMLPE-----------ALRVECGRDSRLDAIFHPPEPVAVP-AVLDSSAREQL
QHI99590.1      AMAQRVDAIQSQLDASGDYQAYLAANLEIVRELREKS--GAAGSAAEPQALFLNNDIEKW
                 :  *:  : .:           * .:*  *: * .:   .. ..* * *:: ..  *:

QHI99589.1      RADLASGRLASPQALADALVGAAHRYVIAELTKAHPDKAVPIGNFSLRDINRRQIVPVAQ
QHI99590.1      RAALLDGSIDSPEKLTAVLLDAVDRNYAVRLARLYPGAAMKIGKYGLGGV-KHGYVSSTE
                ** * .* : **: *: .*:.*. *   . *:. :*. *: **::.* .: ..  *. ::

QHI99589.1      TDEFFTQQQPFIVDSLHAALMRIADELGHAGPQATEASASIDSTISTGLILFNQLMELQA
QHI99590.1      LSEFTQENLQATLAAAHTAMLEMLNDPAFA---PSAGGPEIGGILSDGLLLLQNILEI--
                 .**  ::    : : *:*:: : :: ..*   .: ....*.. :* **:*:::::*:

QHI99589.1      APGWRMHAETLRKLATRQPSTVDEMRDILSRSMAARMLKEAKFGYARGTQWEEEKTLHFF
QHI99590.1      PQRHQANDYVLREIAQTQPSTPEHIGEILEQSMMQRQVTLKKFGH---TSVEERTSVDFF
                .  .. :  .**::*  **** : : :**..**  * :.  ***:   *. ** .:: **

QHI99589.1      WGAERLLLRPESRMVLGRVLQDQLIAAKEQHARAVARRQKLDPALTQALVDSAASHVEHY
QHI99590.1      WTALKLLRKPASKMALGKAIQDLIIQFKQVQAQAIGKRQQLPPDGIEALLDNTVSHVRHY
                * * .** .* *.*.**..:** :*  *: :*.*:..**:* *   :**:*.:.*** **

QHI99589.1      FGFMSSEARPFVRPPEPAAPPAAARPDPLQRYRRAIDELDEGTDIDDPQDPRFVSGGDFL
QHI99590.1      YDFMAQAADPFATPAGMAPFSETDAP-----YRQAVEDLENGTRIADPKDPRFVSASDFL
                :.**:. * **. *.  *. . :  *     **.*:::*::** * **:******..***

QHI99589.1      ARSNALLAVSRAEQQADAAGEPFDHQAFATASIDPDVIASAGYLKTLFDRWAA
QHI99590.1      LRSHALLEVNRAERAARIAGTPFDRAAFIDSSTNADVAAAAPYLQASFSRWPA
                 **:*** *.***. *  ** ***. **  :* :.** *:* **:: *.**.*
```

### Supplementary Data S2

**HrpX-regulated protein family 7**

>QHI99757.1 hypothetical protein GT347_18300 [Xylophilus rhododendri]
MAWALAELSATELTARNAMRKVREVFARRQLNVYCRHLPGWLALARPDLYEDRWPGLGRMYQELRHACST
GTRLLDAGGTWRQHITWYARHYGLPLTHFLALHETALVHQLVWELRGSLPSRDTIATLMQTCGGGGMHTV
QGQARIVDHTNSIVTCVNAMAEHGFPTAGLEICMLLLARGVRLTKEQMRDLPLAVPGLAYAGGWYFPEAE
QPVRFLGCRCDDPELQLFRGFAPMGLDSTGQASFAKHLVHSHPEGCSLQQCQALLRAWNTGLDERAACDY
TAAFRRNAWLLPISETRFTVDPDLVCTSLRQFILANCTQPGSWLAHRHMPDAASQTGETPADGHGVRESR
LVFGLLAQAVQRHALDSYLQTDCRGRQVFHNKKIRQQLLASHPGLCDTAIRDCLAHELPAHIDRTLAARQ
WLKAPASREQLAASLATLQARLPLLAFTDRSVLAAWVNRIDPEHYCTAKMLSQYIRRLPNSEARAVKKLD
ELIGGASLSCYLVPECRLLDLYALRNQLGLQGISITRKRLMELVLERRDAIAAAQGDPVDARLWDTSSRR
QARVREIMALRKLEPGSPMPSWPLFEEALEQVLQVHAPGGARRHFSGSCAMEIYGHLELELAQRPSPRPT
ADPLPEPAPLPQQPDGAADLQRRKRTAGEAQDMPDEPVARRTRSAAARVPRAGAERRGRPGIRS

>QHI99758.1 hypothetical protein GT347_18305 [Xylophilus rhododendri]
MPAPPPWPARDAAGAPAAAFQPVPKPRKSDRLMPRPADVAWALAKLPEELNSVPAVLAAVGQALSARGLA
TSCRHLRAWIALARPDLFAEQWPGLRELFDQALATCRREPWRLEDSAGFRQLIAQCAQAHELAPNHFLAT
HESALAHRLIFEVAQRAPSRKEMCMLMGTQFSLRLARGREHITPHAQAISECVAAMAREGFATTDAEITV
LLLRRQIRLNSRQLKDLPLAVPGIAHSDRWFHPPEDAPDKYFVFRMDDPGAWLFRGVAPIGIEGTGQPSF
ARHLVTTEPAGCSTARCQALLAAWAPLLERTVVALQIERFLRSGWLLDRGADRLVVNPELVCGWLRRRIL
AGWPPHQPPAVAPAPPPPADREPDAFDDEADIDTVPIPRYGIAHTRAVFEMLASCDAAGLAALVTDDYLG
REIFDFRKISQALASHRPAVGRAAIVFILQGHTPGSIGRTLAAAQWRLENSSSEQLAINLAALRSRMPLL
AFAKRCTIARWINLIDPQHWCSTGFLQHYIRTLPIDEQRAARTLDAMIRDERILLFANRDCSGVHTSTLR
THLGLMGIGVDANRLKLLLKERREQIRVVRQEFIPPAGWRTRLQQQLRVQELMCTLGLQRGGRMPSWKTY
MKAFQATFCARRGQPPLRFSGICAMLGYSELALELEPRMLRGQSPKRPAEDPESPPSARRARHDGPPAAG
PPDPGPRLG

>QHI99760.1 hypothetical protein GT347_18315 [Xylophilus rhododendri]
MGWPPLRTPDAACAPAGSPWPCSVAAPQAIDPPPGRLDENTEALLHLDGGTPPPQKRIARDTRPMASARD
VALALQHLPPGPVGILQVEAVVEQRLAAGGLGVVGEHLRPWIALARPELFAELWPGLAGMSAFVLKACRQ
DRGLLGNSERFRALIAWCATHSGAALTHFLARHEAALIHAVMQLLNLGAPNLRQLSCIMGTAGAAKDDAL
RQRLMPHARAIQDCVGGMAREGFATAVVEIEMLLLGCGLRLSRLQMDALPNLLEGLAYTQNWFYPVGQSP
QRLRAFQAGDPVQRLFRGHPAVGGEPARGSFARHLLETHPAGFGIEQCRDLLLGWDRSLDADAARRMTET
FCDEGWLTGVQNGLLSVNPERVCGTMRESILANIPALDPRPSIRRQAMAPPDLAPPTRGPEPAPQQGMQD
DTQPGAEEARVVFELLQSACPGQLADYVRRDPLGREHFDLEKIHQAIRPQRPDIDVRRIDICLQGETPAR
IAATLAARPWLREPAQHRRLRADLTQLWQQLPLLAFVQPGMTAAWINLKDPDHFCTAGILQDLLARQRTW
NTRAAAALDALIDDGHIHYFMRPDLRTLDQAALHTHMGLLGLSIRADRLAMLVACRQGRIDLQPQAFIDP
ELWNSAGRLQQRARELMAVLGLRRGDNMPPWYNFTRALEQTLARHHPGQPARRYTGACAMQIHASLRLEL
ELQLLPDAASKRPAEDAEPLPAAKRASGPGGLPDALDASELAGRMQACRAMPGFGDACEAIGEATGGVPP
DDAFARWLARRLPWPPGASPAPPGTQPEHDLSLGLVARFLARTQGRRWLPVDKLGQPQARDGGRAFVNKI
AGHVCAPGAVGLLRLPRAGQIVAVRSVDGMASACVGGGAVRLAELLAWDLREPPRRQAEFYLPCSS

>QHI99761.1 hypothetical protein GT347_18320 [Xylophilus rhododendri]
MDAHAGDGALDGSIALVLAGLPLPNIDTAIATGSQARPFQPMLTTPRDIAAIVERLAPGPDNSKTLLRIV
RCELNCQGLAPSDIHLKGWTALLRPELFAVQWPGLQAFRDDMLAACRGDPALLDDRAAFRRLIAEQAATV
PRLLRSFLAVYEAPLIHYLIQDQGEATAERQRISRLMGTDQADSAAKLRSYIAPHAQDILDCVNTMAAQG
FATSSVEIDMLLMLRQLRLNLRQMHKLGRLLPGLECRHGWLFPKGQSPRKFMHYRLDDENMRIFRGEASA
GIVPDGRAGFAHCLQQAYPAGGSLNQCLELLEVWAPTLDASARLTHLHNFRSRGLLVSARAGRLAVNPDL
VCRRTRELILRGTPGLGPAAGQGCAAAAEPASPADEQPAESAAPEPAPYGLTEAVMTFECLLQARDEGRL
GDYLGRDLFGREIFQLEAIRQALRPARAHPRLTAIKEALKGLTPARLDDTLAARQWQRPRVAMARLPARL
VSLARELPLLAFAGHSQIAAHLNRHDPTHFCTSAMVTDLVKRQKHDGKATARAIADLAARKLLHLFVSAD
GRRPNLTALRMQLILSNAGLDRPGLQTLLRPYRHLMILRRSRLIDAGLWNGSAGLRETARRLKHSLGLRQ
GDPLPDWKRYMEHFHQLLCQRGPGEQPLLYTGLGAMRIYAELEQELGAADEPAPTTKRAAEDGEDAAPAK
RARIETPDAAPAAPAPPAPAPWNAFEQRPQLRLKTLFPGRGGLYELVCAIGNALQDPDAVESLEGLADCE
ATLDLALHRLSHAPGESKGVRSTPDGQGWALLQLSARQWQAGAADLLNSLDMAVVQRGAGEFAALRRGAD
SLWLLHAQQAEPQQAPPPGELALADGRFAAALPAAACRGAIGRQLSYQAIAGFAAACDAIAELTGDAPPL
DDFARWLDDPFEPTEGMPHEPLDALRRRLHPFIGLEQVTQYLQERHGQAWVPGEALPSLLPWDDAQSLMP
LLELFTRGMDPAQAVLLALPGTPHVLVVRQQAGMPMAQGDAGPQPLQELLAWIAQGLQDLQEVECYWPCD
TDLSSGPAAAW

>QHI99762.1 hypothetical protein GT347_18325 [Xylophilus rhododendri]
MAFLIRAAGNSDAARHLPPSPAPGADSTGTGAAPSPAASGTGDSPESSGAQAQDFAAARRLPLTPTDVID
VLASLTPDRQDSAEAVLREVKPLLLARGRTLSAQHLKPWVALERPELFAATWPGLAQLRAGLLDACRRDS
MLLANSKAFRRMMAECAASHGAELQQFITSHESALIHHLQHHLNGGTLVRTQMSSLMGTRQPCTPRTLAH
LIAEHRQAIMACVSYMAGQGFPTMMLEIRALLGSRELRLNPAQLRDLTRVMPELERKGFWLFPKGQAPWR
FTRFRANDAVLALFRGQPATGSTTPLRGGFALYLLQQHPAGCSAAECDAALAHFAPELSAPLRRARFESW
RKRRLLQTLDGLWFTINPALVCKDMRRRICSRLPRQRGRASQA

>QHI99763.1 hypothetical protein GT347_18330 [Xylophilus rhododendri]
MAFLPSVRSSTDSTPFARTLSSSDECSSSPGLIHEPLPPAALTQTARPGDFTDPRRATNVSDVMALLERL

### Supplementary Data S2

```
PRSTRDSSVSVIRAVKRALQERGFSLPRMAVRPWVALARPRLFSAELAGLKAVRDSLMEACRDDVMALCD
HASFRPLLASCTACADIPTRDFLHTHESALVHYLFHHLRGRPATRQEVSRLMGTDQQPGNQNLARALEPH
KQAVYDQVRHLAAQGFATPLTEIDALLSARGRRLTNSQLPQLAELLPELERSRGWVYPRGQAPKSFPGCR
PDNPAFVLFRGQPAIGDRPASAGFAFYLHQQHPGGCPSQECEKALARFDPQLDRATRHEYIDTLLLRRLL
QPLDMHLVIVNPDMVCARLRESILASCPRRGCSPDDNGAPDEKPESDGPDEPSGRVAAPSPERDGQTIGH
GVADARRVFEQLRSLSIDGRLGDCLARDELGRPILDPSALQRMLQQQHPGLRIRLAYEVLKGLTPGGIAA
TLAQRQWLKPPPTSRQLCACLKALQKELPLIAFAPHNQIVACINRRWPQFFCTTQMLDDLMQRSRSAKAS
VAKVIDELIASQRIFLFANARATALNPHTLRSHLTFSGNCTTPALVRLLLAERWDRLRFPPTPFIDSMLW
GSPRGRQQNARLLMQRLEHTPGGPMPDWSTFMQAFEQEFCPPNEQGVRPLYQGGAPINLYLALELEYALE
SEARSDGAASAPAVPPARPARPAPPQRQRRRPVDLGEDDPAPKKMATQAPAPGWLRFDRSPRILTRSADA
RPGGLYELACAIGNALQDFAQVGLPGHRPDHPASLQTALLDFGPDAPGTVPQARRTPDGAAWELLQLDAD
HWALVGPLLLADLDIAVVQHQAPAGSFAALRREGRALMLLESSGAEARPLAAAEFQALFRRPDGGFAAAV
PGAACRGAASRQLSYASFPGYVQACEAIGEAMWCDPPLDSFARWLDDRLDAPPGTPHAQMHRLRREAWMG
FSHGQRVAGFLLERHRSRWWPVPGMPPLTHQKAFLGALRAQLSQHRQVGLLSLPGAFPVVAAGIVNGELA
ARFGRVSRPLAPLVEAAARQAVQWPPCTLYLQSS

>QHI99764.1 hypothetical protein GT347_18335 [Xylophilus rhododendri]
MKDTSGGVTEAGLVFELLEEAVRDGHLARYTRGRGFDVHAILCQLQHRTPGLPLAAVTDTLSGQTPAGLA
ALLQARQWLRPRPSAEQLRASLDALGRRLPLLACVPRHRLAAWVNQYDPAHYCTTAMVVEFVERQAMNAQ
ALDEALDRLSASGHLALFANISACGLRLSHLRNELALAGIGVRSDALQSLLRKRPIARLLADADLAPADY
CSGPAALQQSLYWLTHEMGLTPGDDMPPWDRFLAQFQRLFCKRRRGRPAQNFHGGGAMSLYMELELTLAP
RSGTTAARAGQEDRDAASAQPGAGTPPPQPSPPPGRWNDYDRDPEFLLSPSQQVLQGGLFALACAIGNAL
QDPPQVDPLGGVADRPAALETALDWFVHAPGEYFGPRSTPDGQPWTLLQLPARHWRAAAPCLLDDLDMAV
VQRGEGQFVALRRSWRGLWLLLASHPLPMALPKPRELALADGRFTAALPAAACHGALGRQVSYQAIAGFA
AACDAIADLDGGEPPLDDFAAWLDGPLGPVTALPHATLDALRGQYHRFIAPTQVAAYLLERRRQMWLQGL
PLPPLLPQDEGASLLAALDARTSELHGGNPLLLGLPGTPHLLLVRRAEGMLLCHEDPTLVALQERLRRIA
ATLAAEQPVLWCWLQPPVGPQPAGELAATQEQQGD

>QHI99765.1 hypothetical protein GT347_18340 [Xylophilus rhododendri]
MAYFPSVTSLPVHWPDGNDIANEPDGIDDSLMVVSFPETEAGADHDRFRPVSPRQRSHRLQPTPQDIADA
LHASDEEPASREAALRLVMQQLRARGLRLRGGHVDAWLALVRPELHADRWPGLAQLRDELLDACRRRPAL
LRDSTVFRGLIATRMRGASPALRRFVAAYESAVIHYLAQQVKGDIADRPLLSRLMGTNLHDSQQKLQAGV
ALHAQAIQDCVQAMAGYGFPTAMVELDMLLQARQLRLNLRQMLQLPQLIPALDYAAGWFFPRDQAPRRFL
GTRMVDKSLALFRGEAAAGTQAGVRGGFAYELLCRYPGGCTIGECELLLAQYATGLHAAERRAHMRNFRR
RFLQSLDGQRLRIHPDIVCEPIRQLILENRPPRAPPTGHRRGLAASRRRRKPRPLM


QHI99762.1      MAFLIRAAGNSDAARHLPPSP----APGADSTGTGAAPSPAASGTGDSPESSGAQAQDFA
QHI99763.1      MAFLPSVRSSTDSTPFARTLS----SSDECSSSPGLIHE----PLPPAALTQTARPGDFT
QHI99765.1      ------------------------------------------------------------
QHI99761.1      -------------------MD----AHAGDGALDGSIALVLAGLPLPNIDTAIATGSQAR
QHI99764.1      ------------------------------------------------------------
QHI99757.1      ------------------------------------------------------------
QHI99758.1      -------------MPAPPPWP-----------------ARDAAGAPAAAFQPVPKP---R
QHI99760.1      MGWPPLRTPDAACAPAGSPWPCSVAAPQAIDPPPGRLDENTEALLHLDGGTPPPQKRIAR


QHI99762.1      AARRLPLTPTDVIDVLASLTPDRQDSAEAVLREVKPLLLARGRTLSAQHLKPWVALERPE
QHI99763.1      DPRRAT-NVSDVMALLERLPRSTRDSSVSVIRAVKRALQERGFSLPRMAVRPWVALARPR
QHI99765.1      ----------------------------------------------------MAYFPSVT
QHI99761.1      PFQPMLTTPRDIAAIVERLAPG-PDNSKTLLRIVRCELNCQGLAPSDIHLKGWTALLRPE
QHI99764.1      ------------------------------------------------------------
QHI99757.1      -----------MAWALAELSAT-ELTARNAMRKVREVFARRQLNVYCRHLPGWLALARPD
QHI99758.1      KSDRLMPRPADVAWALAKLPEE-LNSVPAVLAAVGQALSARGLATSCRHLRAWIALARPD
QHI99760.1      DTRPMA-SARDVALALQHLPPG-PVGILQVEAVVEQRLAAGGLGVVGEHLRPWIALARPE


QHI99762.1      LFAATWPGLAQLRAGLLDACRRDSMLLANSKAFRRMMAECAASHGAELQQFITSHESALI
QHI99763.1      LFSAELAGLKAVRDSLMEACRDDVMALCDHASFRPLLASCTACADIPTRDFLHTHESALV
QHI99765.1      SLPVHWPD----------------------------------------------------
QHI99761.1      LFAVQWPGLQAFRDDMLAACRGDPALLDDRAAFRRLIAEQAATVPRLLRSFLAVYEAPLI
QHI99764.1      ------------------------------------------------------------
QHI99757.1      LYEDRWPGLGRMYQELRHACSTGTRLLDAGGTWRQHITWYARHYGLPLTHFLALHETALV
QHI99758.1      LFAEQWPGLRELFDQALATCRREPWRLEDSAGFRQLIAQCAQAHELAPNHFLATHESALA
QHI99760.1      LFAELWPGLAGMSAFVLKACRQDRGLLGNSERFRALIAWCATHSGAALTHFLARHEAALI
```

**Supplementary Data S2**

```
QHI99762.1      HHLQHHLNGGTLVRTQMSSLMGTRQPCTPRTL--AHLIAEHRQAIMACVSYMAGQGFPTM
QHI99763.1      HYLFHHLRGRPATRQEVSRLMGTDQQPGNQNL--ARALEPHKQAVYDQVRHLAAQGFATP
QHI99765.1      ---------------------GND-------------IANEPDGIDDSLMVVS-------
QHI99761.1      HYLIQDQGEATAERQRISRLMGTDQADSAAKL--RSYIAPHAQDILDCVNTMAAQGFATS
QHI99764.1      ------------------------------------------------------------
QHI99757.1      HQLVWELRGSLPSRDTIATLMQTCGGGGMHTVQGQARIVDHTNSIVTCVNAMAEHGFPTA
QHI99758.1      HRLIFEVAQRAPSRKEMCMLMGTQFSLRLARG--REHITPHAQAISECVAAMAREGFATT
QHI99760.1      HAVMQLLNLGAPNLRQLSCIMGTAGAAKDDAL--RQRLMPHARAIQDCVGGMAREGFATA

QHI99762.1      MLEIRALLGSRELRLNPAQLRDLTRVMPELERKGFWLFPKGQAPWRFTRFRANDAVLALF
QHI99763.1      LTEIDALLSARGRRLTNSQLPQLAELLPELERSRGWVYPRGQAPKSFPGCRPDNPAFVLF
QHI99765.1      -------------------------------------FPETEAGADHDRFRPVSPRQRSH
QHI99761.1      SVEIDMLLMLRQLRLNLRQMHKLGRLLPGLECRHGWLFPKGQSPRKFMHYRLDDENMRIF
QHI99764.1      ------------------------------------------------------------
QHI99757.1      GLEICMLLLARGVRLTKEQMRDLPLAVPGLAYAGGWYFPEAEQPVRFLGCRCDDPELQLF
QHI99758.1      DAEITVLLLRRQIRLNSRQLKDLPLAVPGIAHSDRWFHPPEDAPDKYFVFRMDDPGAWLF
QHI99760.1      VVEIEMLLLGCGLRLSRLQMDALPNLLEGLAYTQNWFYPVGQSPQRLRAFQAGDPVQRLF

QHI99762.1      RGQPATGSTTPLRGGFALYLLQQHPAGCSAAECDAALAHFAPELSAPLRRARFESWRKRR
QHI99763.1      RGQPAIG-DRPASAGFAFYLHQQHPGGCPSQECEKALARFDPQLDRATRHEYIDTLLLRR
QHI99765.1      RLQP--------------------------------------------------------
QHI99761.1      RGEASAGIVPDGRAGFAHCLQQAYPAGGSLNQCLELLEVWAPTLDASARLTHLHNFRSRG
QHI99764.1      -MKDTSG-----------------------------------------------------
QHI99757.1      RGFAPMGLDSTGQASFAKHLVHSHPEGCSLQQCQALLRAWNTGLDERAACDYTAAFRRNA
QHI99758.1      RGVAPIGIEGTGQPSFARHLVTTEPAGCSTARCQALLAAWAPLLERTVVALQIERFLRSG
QHI99760.1      RGHPAVG-GEPARGSFARHLLETHPAGFGIEQCRDLLLGWDRSLDADAARRMTETFCDEG

QHI99762.1      LLQTLDGLWFTINPALVCKDMR--------------------------------------
QHI99763.1      LLQPLDMHLVIVNPDMVCARLRESILASCPRRGCSPDDNGAPDEKPESDGPDEPSGRVAA
QHI99765.1      ------------------------------------------------------------
QHI99761.1      LLVSARAGRLAVNPDLVCRRTRELILRGTPGLGPAAGQGCAAAAEPASPADEQPAESAAP
QHI99764.1      ------------------------------------------------------------
QHI99757.1      WLLPISETRFTVDPDLVCTSLRQFILANCTQPGSWLAHRHMPDAASQTGET---------
QHI99758.1      WLLDRGADRLVVNPELVCGWLRRRILAGWPPHQP-------PAVAPAPPPPADREPDAFD
QHI99760.1      WLTGVQNGLLSVNPERVCGTMRESILANIPALDPRPSIRRQAMAPPDLAPPTRGPEPAPQ

QHI99762.1      ------------------------------------------------------------
QHI99763.1      PSPERDGQT-IGHGVADARRVFEQLRSLSIDGRLGDCLARDELGRPILDPSALQRMLQQQ
QHI99765.1      ------------------------------------------------------------
QHI99761.1      EP--------APYGLTEAVMTFECLLQARDEGRLGDYLGRDLFGREIFQLEAIRQALRPA
QHI99764.1      -------------GVTEAGLVFELLEEAVRDGHLARYTR----GRG-FDVHAILCQLQHR
QHI99757.1      --------PADGHGVRESRLVFGLLAQAVQRHALDSYLQTDCRGRQVFHNKKIRQQLLAS
QHI99758.1      DEADIDTVPIPRYGIAHTRAVFEMLAS-CDAAGLAALVTDDYLGREIFDFRKISQALASH
QHI99760.1      QGMQDDTQP----GAEEARVVFELLQSAC-PGQLADYVRRDPLGREHFDLEKIHQAIRPQ

QHI99762.1      -------------------------------------RRICSRL--------PR------
QHI99763.1      HPGLRIRLAYEVLKGLTPGGIAATLAQRQWLKPPPTSRQLCACLKALQKE-LPLIAFAPH
QHI99765.1      ----------------TPQDIADALHASD-EEPASREAALRLVMQQLRARGLRL----RG
QHI99761.1      RAHPRLTAIKEALKGLTPARLDDTLAARQWQRPRVAMARLPARLVSLARE-LPLLAFAGH
QHI99764.1      TPGLPLAAVTDTLSGQTPAGLAALLQARQWLRPRPSAEQLRASLDALGRR-LPLLACVPR
QHI99757.1      HPGLCDTAIRDCLAHELPAHIDRTLAARQWLKAPASREQLAASLATLQAR-LPLLAFTDR
QHI99758.1      RPAVGRAAIVFILQGHTPGSIGRTLAAAQWRLENSSSEQLAINLAALRSR-MPLLAFAKR
QHI99760.1      RPDIDVRRIDICLQGETPARIAATLAARPWLREPAQHRRLRADLTQLWQQ-LPLLAFVQP
                                                       :   :
```

**Supplementary Data S2**

```
QHI99762.1      ------------------------------------------------------------
QHI99763.1      NQIVACINRRWPQFFCTTQMLDDLMQRSRSAKASVAKVIDELIASQRIFLFANARATALN
QHI99765.1      GHVDAWLALVRPELHADRW-------------PGLAQLRDELL-----------DACRRR
QHI99761.1      SQIAAHLNRHDPTHFCTSAMVTDLVKRQKHDGKATARAIADLAARKLLHLFVSADGRRPN
QHI99764.1      HRLAAWVNQYDPAHYCTTAMVVEFVERQAMNAQALDEALDRLSASGHLALFANISACGLR
QHI99757.1      SVLAAWVNRIDPEHYCTAKMLSQYIRRLPNSEARAVKKLDELIGGASLSCYLVPECRLLD
QHI99758.1      CTIARWINLIDPQHWCSTGFLQHYIRTLPIDEQRAARTLDAMIRDERILLFANRDCSGVH
QHI99760.1      GMTAAWINLKDPDHFCTAGILQDLLARQRTWNTRAAAALDALIDDGHIHYFMRPDLRTLD


QHI99762.1      ------------------------------------------------------------
QHI99763.1      PHTLRSHLTFSG-NCTTPALVRLLLAERWDRLRFPPTPFIDSMLWGSPRGRQQNARLLMQ
QHI99765.1      PALLRDSTVFRGLIATRMRGASPALRRFVAAYESAVIHYLAQQVKGDIADRPLLSRLMGT
QHI99761.1      LTALRMQLILSN-AGLDRPGLQTLLRPYRHLMILRRSRLIDAGLWNGSAGLRETARRLKH
QHI99764.1      LSHLRNELALAG-IGVRSDALQSLLRKRPIARLLADADLAPADYCSGPAALQQSLYWLTH
QHI99757.1      LYALRNQLGLQG-ISITRKRLMELVLERRDAIAAAQGDPVDARLWDTSSRRQARVREIMA
QHI99758.1      TSTLRTHLGLMG-IGVDANRLKLLLKERREQIRVVRQEFIPPAGWRTRLQQQLRVQELMC
QHI99760.1      QAALHTHMGLLG-LSIRADRLAMLVACRQGRIDLQPQAFIDPELWNSAGRLQQRARELMA


QHI99762.1      ------------------------------------------------------------
QHI99763.1      RLEHTPGGPMPDWSTFMQAFEQEFCPPNEQGVRPLYQGGAPINLYLALELEYALESEARS
QHI99765.1      NLHDSQQKLQAGVALHAQAIQDCVQAMAGYGFPTAMVELDMLLQARQLRLNL--------
QHI99761.1      SLGLRQGDPLPDWKRYMEHFHQLLCQRGPGEQPLLYTGLGAMRIYAELEQELGAADEPAP
QHI99764.1      EMGLTPGDDMPPWDRFLAQFQRLFCKRRRGRPAQNFHGGGAMSLYMELELTL-----APR
QHI99757.1      LRKLEPGSPMPSWPLFEEALEQVLQVHAPGGARRHFSGSCAMEIYGHLELEL--------
QHI99758.1      TLGLQRGGRMPSWKTYMKAFQATFCARR-GQPPLRFSGICAMLGYSELALEL--------
QHI99760.1      VLGLRRGDNMPPWYNFTRALEQTLARHHPGQPARRYTGACAMQIHASLRLEL--------


QHI99762.1      ------------------------------------------------------------
QHI99763.1      DGAASAPAVPPARPARPAPPQRQRRRPVDLGEDDPAPKKMATQAPAPGWLRFDRSPR-IL
QHI99765.1      -------------------------------------------------RQMLQLPQ--L
QHI99761.1      TTKRAAEDGEDAAPAKRARIETPDAAPAAPAPPAPAP-----------WNAFEQRPQ-LR
QHI99764.1      SGTTAARAGQEDRDAASAQPGAGTPPPQPSPPPGR-------------WNDYDRDPEFLL
QHI99757.1      --------------------------------------------------AQRPSPR---
QHI99758.1      ------------------------------------------------------EPR-ML
QHI99760.1      --------------------------------------------------ELQLLPDAAS


QHI99762.1      ------------------------------------------------------------
QHI99763.1      TRSADARPGGLYELACAIGNALQDFAQVGLPGHRPDHPASLQTALLDFGPDAPGTVPQAR
QHI99765.1      IPALDYAAG---------------------------------------------------
QHI99761.1      LKTLFPGRGGLYELVCAIGNALQDPDAVESLEGLADCEATLDLALHRLS-HAPGESKGVR
QHI99764.1      SPSQQVLQGGLFALACAIGNALQDPPQVDPLGGVADRPAALETALDWFV-HAPGEYFGPR
QHI99757.1      ------------------------------------------------------------
QHI99758.1      RG------------------------------------------------QSPKR-----
QHI99760.1      KR----------------------------------------------------------


QHI99762.1      ------------------------------------------------------------
QHI99763.1      RTPDGAAWELLQLDADHWALVGPLLLADLDIAVVQHQAPAGSFAALRREGRALMLLESSG
QHI99765.1      -----------------------------------------------------WFFPRDQ
QHI99761.1      STPDGQGWALLQLSARQWQAGAADLLNSLDMAVVQRG--AGEFAALRRGADSLWLLHAQQ
QHI99764.1      STPDGQPWTLLQLPARHWRAAAPCLLDDLDMAVVQRG--EGQFVALRRSWRGLWLLLASH
QHI99757.1      --------------------------------------------------------PTAD
QHI99758.1      --------------------------------------------------------PAED
QHI99760.1      --------------------------------------------------------PAED
```

### Supplementary Data S2

```
QHI99762.1      -----------------QRGRASQA-----------------------------------
QHI99763.1      AEARPLAAAEFQALFRRPDGGFAAAVPGAACRGAASRQLSYASFPGYVQACEAIGEAMWC
QHI99765.1      APRRFLGTRMVDKSLALFRGEAAAGTQAGVRGG-----FAYELLCRYPGGCT----IGEC
QHI99761.1      AEPQQAPPPGE---LALADGRFAAALPAAACRGAIGRQLSYQAIAGFAAACDAIAELTGD
QHI99764.1      PLPMALPKPRE---LALADGRFTAALPAAACHGALGRQVSYQAIAGFAAACDAIADLDGG
QHI99757.1      PLPEPAPLPQQ------PDGAADLQRRKRTAGE---------------------------
QHI99758.1      PESPPSARRAR------HDG----------------------------------------
QHI99760.1      AEPLPAAKRAS------GPGGLPDALDASELAG---RMQACRAMPGFGDACEAIGEATGG
                                   *

QHI99762.1      ------------------------------------------------------------
QHI99763.1      DPPLDSFARWLDDRLDAPPGTPHAQMHRLRREAWMGFSHGQRVAGFLLERHRSRWWPVPG
QHI99765.1      ELLLAQYATGL------HAAERRAHMRNFRRR-------------FLQSLDGQRLRIHPD
QHI99761.1      APPLDDFARWLDDPFEPTEGMPHEPLDALRRR-LHPFIGLEQVTQYLQERHGQAWVPGEA
QHI99764.1      EPPLDDFAAWLDGPLGPVTALPHATLDALRGQ-YHRFIAPTQVAAYLLERRRQMWLQGLP
QHI99757.1      -----------------AQDMPDEPVARRTRS----------------------------
QHI99758.1      -----------------PPAAGPPDPG---------------------------------
QHI99760.1      VPPDDAFARWLARRLPWPPGASPAPPGTQPEH----DLSLGLVARFLARTQGRRWLPVDK

QHI99762.1      ------------------------------------------------------------
QHI99763.1      MPPLTHQ---KAFLGALRAQLSQ--HRQVGLLSLPGAFPVVAAGIVNGELAARFGRVSRP
QHI99765.1      I-------------------VCEPIRQLILENRPPRAPP---TGHRRGLAASRRRRKPRP
QHI99761.1      LPSLLPWDDAQSLMPLLELFTRGMDPAQAVLLALPGTPHVLVVRQQAGMPMAQGDAGPQP
QHI99764.1      LPPLLPQDEGASLLAALDARTSELHGGNPLLLGLPGTPHLLLVRRAEGMLLCHEDPTLVA
QHI99757.1      -----------------------------AAARVPRAG-----------AERRGRPGIRS
QHI99758.1      ----------------------------------PRLG----------------------
QHI99760.1      LGQPQARDGGRAFVNKIAGHVCA--PGAVGLLRLPRAGQIVAVRSVDGMASACVGGGAVR

QHI99762.1      --------------------------------------------
QHI99763.1      LAPLVEAAARQAVQWPPCTLYLQSS-------------------
QHI99765.1      LM------------------------------------------
QHI99761.1      LQELLAWIAQGLQDLQEVECYWPCDTDLSSGPAAAW--------
QHI99764.1      LQERLRRIAATLAAEQPVLWCWLQPPVGPQPAGELAATQEQQGD
QHI99757.1      --------------------------------------------
QHI99758.1      --------------------------------------------
QHI99760.1      LAELLAWDLREPPR-RQAEFYLPCSS------------------
```

### Supplementary Data S2

**HrpX-regulated protein family 8**

```
>QHJ00715.1 hypothetical protein GT347_23660 [Xylophilus rhododendri]
MFPSSHQSSAASGQTQAQDTQSTASAATTTTPTTTTDRSADPGGTSSALAPLPRQDTAGPADSHASQLSR
SLFTQPGPTAPSTAEIEELDRLIDRALAAKWLQEGLSKCEALDCPPDRNPSPGELARSLASSHDTLVLFL
AMSREMELALMPRFVETLLRLGMIDGRHAAAAREAAVRVAQYAGLRGPGYGGGSGRSGTTWRRHYMIMPT
LRLAIDPPALERALTDPGQWPAMLAISGTQGVLNDGAIAMLGRLLQDLCSDMAASGFLPAGLCTRLHEQV
PAILQAARQFDRRPGT

>QHJ00716.1 hypothetical protein GT347_23665 [Xylophilus rhododendri]
MDSRFPGLPPRLPDTGTSGCTAAELIQKIRSCEKKEAPALVAALRAMMECQPPVNAAHRVDAGKVPAGHE
AELQALVDRAFRVEHHQACIAAIARLFNELEADARRDPVALALAIQHDELTLWGNHPAEVLSAHMPHLIE
TMLAWGLIDARHAPLARTAARRVAHYEALYQGRLGMTPEPELALTGARRTRLFTARLLRAAMDPASLATT
LADPDRWASLIGLPQLQGPMHLSAYRALIKDLFGLVDDMQASAFLEVQHCELLRGLIPDIVRARHRQDRA
KHRIPDDWNAHDQNTSAFDADALVSALRSLPTFDDARLRELKSAVKDFYRHGRMTHPEFDFAARFGPMTQ
CLQEDAGKARAFLICLKKRGF
```

```
QHJ00715.1      ------MFPSSHQSSAASGQTQAQDTQSTASAATTTTPTTTTDRSADPGGTSSALAPLPR
QHJ00716.1      MDSRFPGLPPRLPDTGTSGCTAAELIQKI--------------RSCEKKEAPALVAALRA
                       :*.   .:.:** * *:  *.               **.:   :.: :*.*

QHJ00715.1      QDTAGPADSHASQLSRSLFTQPGPTAPSTAEIEELDRLIDRALAAKWLQEGLSKCEALDC
QHJ00716.1      MMECQPPVNAAHRVDAG-------KVPAGHE-AELQALVDRAFRVEHHQACIAAIARLFN
                   . *. . * .:. .       ..*:  *  **: *:***: .:. *  ::    *

QHJ00715.1      PPDRNPSPGELARSLASSHDTLVLFLAMSREMELALMPRFVETLLRLGMIDGRHAAAARE
QHJ00716.1      ELEADARRDPVALALAIQHDELTLWGNHPAEVLSAHMPHLIETMLAWGLIDARHAPLART
                  : :.  . :* :** .** *.*:   . *:  * **.::**:*  *:**.***. **

QHJ00715.1      AAVRVAQYAGLRGPGYGGGSGRSGT-------TWRRHYMIMPTLRLAIDPPALERALTDP
QHJ00716.1      AARRVAHYEAL----YQGRLGMTPEPELALTGARRTRLFTARLLRAAMDPASLATTLADP
                ** ***:* .*    * *  * :         :.* . :    ** *:**.:*  :*:**

QHJ00715.1      GQWPAMLAISGTQGVLNDGAIAMLGRLLQDLCSDMAASGFLPAGLCTRLHEQVPAILQAA
QHJ00716.1      DRWASLIGLPQLQGPMHLSAYRALIKDLFGLVDDMQASAFLEVQHCELLRGLIPDIVRAR
                ..*.:::.:.  ** :: .*   * . * .* .** **.** .  *  *.  :* *:.*

QHJ00715.1      RQFDR-------------------------------------------------------
QHJ00716.1      HRQDRAKHRIPDDWNAHDQNTSAFDADALVSALRSLPTFDDARLRELKSAVKDFYRHGRM
                .. **

QHJ00715.1      ---------------------------------RPGT
QHJ00716.1      THPEFDFAARFGPMTQCLQEDAGKARAFLICLKKRGF
                                                 . *
```
