## Supplementary Data S3 for "Identification of protein secretion systems and type III effectors in wood-associated bacteria of the genus *Xylophilus*"

**Multiple alignment of members of OG_1116 (in green), OG_1115 (in blue) and OG_3308 (in red).**

```
DFQ15_RS17525/WP_233504294      ------------------------------------------------------------
NBM50_RS05530/WP_233504294      ------------------------------------------------------------
NBM50_05530/MCS4509724          ------------------------------------------------------------
DFQ15_105107/PYE78748           ------------------------------------------------------------
DFQ15_105106/PYE78747           MNVDATLEANAQLAESCQPGVRALNHPAMAANRDFHSVRPVATRIQARRMLRFTEEPTPT
DFQ15_RS17380/WP_199399817      ------------------------------------------------------------
NBM50_RS05540/WP_259292613      ------------------------------------------------------------
NBM50_05540/MCS4509725          ------------------------------------------------------------
DFQ15_RS17300/WP_199399909      ------------------------------------------------------------
DFQ15_1541/PYE72703             ------------------------------------------------------------
NBM50_RS17175/WP_259292686      ------------------------------------------------------------
NBM50_17140/MCS4511918          ------------------------------------------------------------


DFQ15_RS17525/WP_233504294      ------------------------------------------------------------
NBM50_RS05530/WP_233504294      ------------------------------------------------------------
NBM50_05530/MCS4509724          ------------------------------------------------------------
DFQ15_105107/PYE78748           ------------------------------------------------------------
DFQ15_105106/PYE78747           ASNRHNSPLSGSVARRVTHRCSLKFTVPANRNNGLAYEIKRYGPPPQVDFSFKAWKDRSW
DFQ15_RS17380/WP_199399817      ------------------------------------------------------------
NBM50_RS05540/WP_259292613      ------------------------------------------------------------
NBM50_05540/MCS4509725          ------------------------------------------------------------
DFQ15_RS17300/WP_199399909      ------------------------------------------------------------
DFQ15_1541/PYE72703             ------------------------------------------------------------
NBM50_RS17175/WP_259292686      ------------------------------------------------------------
NBM50_17140/MCS4511918          ------------------------------------------------------------


DFQ15_RS17525/WP_233504294      ---------------------------MNESDRLQEAQNAGVSQPSGTAPAAGPLAGLPT
NBM50_RS05530/WP_233504294      ---------------------------MNESDRLQEAQNAGVSQPSGTAPAAGPLAGLPT
NBM50_05530/MCS4509724          ---------------------------MNESDRLQEAQNAGVSQPSGTAPAAGPLAGLPT
DFQ15_105107/PYE78748           -----------MKDLYPFATNTAWTHDLNESDRLQEAQNAGVSQPSGTAPAAGPLAGLPT
DFQ15_105106/PYE78747           KFTVRIPSRHEMKDLYPFATGTAWTHDLSDSDRLQEAQNAVLSQQSGTAPAAGPLAGLPT
DFQ15_RS17380/WP_199399817      ---------------------------MSDSDRLQEAQNAVLSQQSGTAPAAGPLAGLPT
NBM50_RS05540/WP_259292613      ---------------------------MSDSDRLQEAQNAVLSQQSGTAPAAGPLAGLPT
NBM50_05540/MCS4509725          ---------------------------MSDSDRLQEAQNAVLSQQSGTAPAAGPLAGLPT
DFQ15_RS17300/WP_199399909      ---------------------------MYEADRLQEMQNAGVSQQSGTALAAGPLAGLPA
DFQ15_1541/PYE72703             -----------MKDLYPFGTGTSWTHDLYEADRLQEMQNAGVSQQSGTALAAGPLAGLPA
NBM50_RS17175/WP_259292686      -----------MKDLYPFGTGTSWTHDLYEADRLQEMQNAGVSQQSGTALAAGPLAGLPA
NBM50_17140/MCS4511918          -----------MKDLYPFGTGTSWTHDLYEADRLQEMQNAGVSQQSGTALAAGPLAGLPA
                                                           : ::***** *** :** **** *********:

DFQ15_RS17525/WP_233504294      RPRGDARPEETRLSLGTTMPTMPTLPILESMNLPTPSV-------PRCDAPRPSRRRRQV
NBM50_RS05530/WP_233504294      RPRGDARPEETRLSLGTTMPTMPTLPILESMNLPTPSV-------PRCDAPRPSRRRRQV
NBM50_05530/MCS4509724          RPRGDARPEETRLSLGTTMPTMPTLPILESMNLPTPSV-------PRCDAPRPSRRRRQV
DFQ15_105107/PYE78748           RPRGDARPEETRLSLGTTMPTMPTLPILESMNLPTPSV-------PRCDAPRPSRRRRQV
DFQ15_105106/PYE78747           RPQGDGRPEETRLALGTTIPT-PT--VIESVYLPSPSIHRSSANFPRFDAPRPSRRRRQV
DFQ15_RS17380/WP_199399817      RPQGDGRPEETRLALGTTIPT-PT--VIESVYLPSPSIHRSSANFPRFDAPRPSRRRRQV
NBM50_RS05540/WP_259292613      RPQGDGRPEETRLALGTTIPT-PT--VIESVYLPSPSIHRSSANFPRFDAPRPSRRRRQV
NBM50_05540/MCS4509725          RPQGDGRPEETRLALGTTIPT-PT--VIESVYLPSPSIHRSSANFPRFDAPRPSRRRRQV
DFQ15_RS17300/WP_199399909      RPQGDGRPEETRRALETAIPT-PT--VIESVYLPSPSIHRSSANFPRFDAPRPSRRGRQV
DFQ15_1541/PYE72703             RPQGDGRPEETRRALETAIPT-PT--VIESVYLPSPSIHRSSANFPRFDAPRPSRRGRQV
NBM50_RS17175/WP_259292686      RPQGDGRPEETRRALETAIPT-PT--VIESVYLPSPSIHRSSANFPRFDAPRPSRRGRQV
NBM50_17140/MCS4511918          RPQGDGRPEETRRALETAIPT-PT--VIESVYLPSPSIHRSSANFPRFDAPRPSRRGRQV
                                **.**.****** :* *::** **  ::**: **:**:       ** ******** ***

DFQ15_RS17525/WP_233504294      VSEQTE-----------------------------QALSKMAAEPYPEDAGIIERFSANA
NBM50_RS05530/WP_233504294      VSEQTE-----------------------------QALSKMAAEPYPEDAGIIERFSANA
NBM50_05530/MCS4509724          VSEQTE-----------------------------QALSKMAAEPYPEDAGIIERFSANA
DFQ15_105107/PYE78748           VSEQTE-----------------------------QALSKMAAEPYPEDAGIIERFSANA
DFQ15_105106/PYE78747           VSEQAPIEVRSRDHSSSELQARVQEAPSAQHALPARARRAIGEIPYPEDAWLITRFQLNA
DFQ15_RS17380/WP_199399817      VSEQAPIEVRSRDHSSSELQARVQEAPSAQHALPARARRAIGEIPYPEDAWLITRFQLNA
NBM50_RS05540/WP_259292613      VSEQAPIEVRSRDHSSSELQARVQEAPSAQHALPARARRAIGEIPYPEDAWLITRFQLNA
NBM50_05540/MCS4509725          VSEQAPIEVRSRDHSSSELQARVQEAPSAQHALPARARRAIGEIPYPEDAWLITRFQLNA
DFQ15_RS17300/WP_199399909      VSEQASVEVNSPDHSSAELHARVQDAPRVQCALPAKAKRAISGIPYPDDAWLITMFRLNA
DFQ15_1541/PYE72703             VSEQASVEVNSPDHSSAELHARVQDAPRVQCALPAKAKRAISGIPYPDDAWLITMFRLNA
NBM50_RS17175/WP_259292686      VSEQASVEVNSPDHSSAELHARVQDAPRVQCALPAKAKRAISGIPYPDDAWLITMFRLNA
NBM50_17140/MCS4511918          VSEQASVEVNSPDHSSAELHARVQDAPRVQCALPAKAKRAISGIPYPDDAWLITMFRLNA
                                ****:                              .*   :.  ***:** :*  *  **
```

**Supplementary Data S3**

```
DFQ15_RS17525/WP_233504294      HASGSNKNTILIYSKNLTRFSAWLRKNDMPGINGRLFDEELTNDLREFTRRTEKWRFTTS
NBM50_RS05530/WP_233504294      HASGSNKNTILIYSKNLTRFSAWLRKNDMPGINGRLFDEELTNDLREFTRRTEKWRFTTS
NBM50_05530/MCS4509724          HASGSNKNTILIYSKNLTRFSAWLRKNDMPGINGRLFDEELTNDLREFTRRTEKWRFTTS
DFQ15_105107/PYE78748           HASGSNKNTILIYSKNLTRFSAWLRKNDMPGINGRLFDEELTNDLREFTRRTEKWRFTTS
DFQ15_105106/PYE78747           EAGGSNLSTASIYSRALTKFSAWLRQNDKPGLHARLFEDELANDLKQFVMQTQKWYIPLA
DFQ15_RS17380/WP_199399817      EAGGSNLSTASIYSRALTKFSAWLRQNDKPGLHARLFEDELANDLKQFVMQTQKWYIPLA
NBM50_RS05540/WP_259292613      EAGGSNLSTASIYSRALTKFSAWLRQNDKPGLHARLFEDELANDLKQFVMQTQKWYIPLA
NBM50_05540/MCS4509725          EAGGSNLSTASIYSRALTKFSAWLRQNDKPGLHARLFEDELANDLKQFVMQTQKWYIPLA
DFQ15_RS17300/WP_199399909      EAGGSNKNTALIYSKILTKLSIWLRENDKPGLHARLFEEELAQDLRKFAMQTQKWYVPMS
DFQ15_1541/PYE72703             EAGGSNKNTALIYSKILTKLSIWLRENDKPGLHARLFEEELAQDLRKFAMQTQKWYVPMS
NBM50_RS17175/WP_259292686      EAGGSNKNTALIYSKILTKLSIWLRENDKPGLHARLFEEELAQDLRKFAMQTQKWYVPMS
NBM50_17140/MCS4511918          EAGGSNKNTALIYSKILTKLSIWLRENDKPGLHARLFEEELAQDLRKFAMQTQKWYVPMS
                                 *.*** .*  ***. **.:* ***:** **::.***::**::**.:*. .*:** .. :

DFQ15_RS17525/WP_233504294      LLCHLRQSETSGMVKIPNIEQKHLQIPNEDNWLLCEAFPKNDQTSKNYKILLKQFSTWLQ
NBM50_RS05530/WP_233504294      LLCHLRQSETSGMVKIPNIEQKHLQIPNEDNWLLCEAFPKNDQTSKNYKILLKQFSTWLQ
NBM50_05530/MCS4509724          LLCHLRQSETSGMVKIPNIEQKHLQIPNEDNWLLCEAFPKNDQTSKNYKILLKQFSTWLQ
DFQ15_105107/PYE78748           LLCHLRQSETSGMVKIPNIEQKHLQIPNEDNWLLCEAFPKNDQTSKNYKILLKQFSTWLQ
DFQ15_105106/PYE78747           ILDHIRQSVTNDSVQIPRLDRKPLQIPDEDDWLICEAFSETDRTSITYRNKLKLFSSWLH
DFQ15_RS17380/WP_199399817      ILDHIRQSVTNDSVQIPRLDRKPLQIPDEDDWLICEAFSETDRTSITYRNKLKLFSSWLH
NBM50_RS05540/WP_259292613      ILDHIRQSVTNDSVQIPRLDRKPLQIPDEDDWLICEAFSETDRTSITYRNKLKLFSSWLH
NBM50_05540/MCS4509725          ILDHIRQSVTNDSVQIPRLDRKPLQIPDEDDWLICEAFSETDRTSITYRNKLKLFSSWLH
DFQ15_RS17300/WP_199399909      LLDHLRESMTNGPLKIPRSDRNLIQIPEEDDWLICEVFSQNDKTSTGYNNQLKRFSSWLH
DFQ15_1541/PYE72703             LLDHLRESMTNGPLKIPRSDRNLIQIPEEDDWLICEVFSQNDKTSTGYNNQLKRFSSWLH
NBM50_RS17175/WP_259292686      LLDHLRESMTNGPLKIPRSDRNLIQIPEEDDWLICEVFSQNDKTSTGYNNQLKRFSSWLH
NBM50_17140/MCS4511918          LLDHLRESMTNGPLKIPRSDRNLIQIPEEDDWLICEVFSQNDKTSTGYNNQLKRFSSWLH
                                :* *:*:* *.. ::**. :.: :***:**:**:**.*.:.*.**  *.  ** **:**:

DFQ15_RS17525/WP_233504294      EKGEPGLCEKDNLHSDRLTAQARTYIQEVPNRECLISALKHLRRYDCGENPVYATKRDTR
NBM50_RS05530/WP_233504294      EKGEPGLCEKDNLHSDRLTAQARTYIQEVPNRECLISALKHLRRYDCGENPVYATKRDTR
NBM50_05530/MCS4509724          EKGEPGLCEKDNLHSDRLTAQARTYIQEVPNRECLISALKHLRRYDCGENPVYATKRDTR
DFQ15_105107/PYE78748           EKGEPGLCEKDNLHSDRLTAQARTYIQEVPNRECLISALKHLRRYDCGENPVYATKRDTR
DFQ15_105106/PYE78747           EKGEPGLCDEGKLHSKYLTEQAKIFSSGKNGRNSLIPALKHLRNFDRTGTKVYQHKNDTR
DFQ15_RS17380/WP_199399817      EKGEPGLCDEGKLHSKYLTEQAKIFSSGKNGRNSLIPALKHLRNFDRTGTKVYQHKNDTR
NBM50_RS05540/WP_259292613      EKGEPGLCDEGKLHSKYLTEQAKIFSSGKNGRNSLIPALKHLRNFDRTGTKVYQHKNDTR
NBM50_05540/MCS4509725          EKGEPGLCDEGKLHSKYLTEQAKIFSSGKNGRNSLIPALKHLRNFDRTGTKVYQHKNDTR
DFQ15_RS17300/WP_199399909      ENGERGLCDEGSLHSDHLTAQARIYMTEGTGRNGLTPALKQLRNFDRTGTKVYQHKNDTR
DFQ15_1541/PYE72703             ENGERGLCDEGSLHSDHLTAQARIYMTEGTGRNGLTPALKQLRNFDRTGTKVYQHKNDTR
NBM50_RS17175/WP_259292686      ENGERGLCDEGSLHSDHLTAQARIYMTEGTGRNGLTPALKQLRNFDRTGTKVYQHKNDTR
NBM50_17140/MCS4511918          ENGERGLCDEGSLHSDHLTAQARIYMTEGTGRNGLTPALKQLRNFDRTGTKVYQHKNDTR
                                *:** ***::..***. ** **. :     .*: * .***:**.:*   . **  *.***

DFQ15_RS17525/WP_233504294      AIPADDRLFSEKYEVWLREQGYGGKPCRR---DSAASRASSFRSFSAWLQHESPSSASLA
NBM50_RS05530/WP_233504294      AIPADDRLFSEKYEVWLREQGYGGKPCRR---DSAASRASSFRSFSAWLQHESPSSASLA
NBM50_05530/MCS4509724          AIPADDRLFSEKYEVWLREQGYGGKPCRR---DSAASRASSFRSFSAWLQHESPSSASLA
DFQ15_105107/PYE78748           AIPADDRLFSEKYEVWLREQGYGGKPCRR---DSAASRASSFRSFSAWLQHESPSSASLA
DFQ15_105106/PYE78747           TIPVEDILLSEKFAAWLREQGYGGKVNRRGKDDGAAVRASALRSFCAWMQRTGRASTSLA
DFQ15_RS17380/WP_199399817      TIPVEDILLSEKFAAWLREQGYGGKVNRRGKDDGAAVRASALRSFCAWMQRTGRASTSLA
NBM50_RS05540/WP_259292613      TIPVEDILLSEKFAAWLREQGYGGKVNRRGKDDGAAVRASALRSFCAWMQRTGRASTSLA
NBM50_05540/MCS4509725          TIPVEDILLSEKFAAWLREQGYGGKVNRRGKDDGAAVRASALRSFCAWMQRTGRASTSLA
DFQ15_RS17300/WP_199399909      TIPVEDILLSEKFAAWLREQGYGGKVNRRGKDDGAAIQASSLRSFCAWMQRTGRASTSLA
DFQ15_1541/PYE72703             TIPVEDILLSEKFAAWLREQGYGGKVNRRGKDDGAAIQASSLRSFCAWMQRTGRASTSLA
NBM50_RS17175/WP_259292686      TIPVEDILLSEKFAAWLREQGYGGKVNRRGKDDGAA------------------------
NBM50_17140/MCS4511918          TIPVEDILLSEKFAAWLREQGYGGKVNRRGKDDGAA------------------------
                                :**.:* *:***: .**********  **   *.**

DFQ15_RS17525/WP_233504294      SRIQSGDRSLEVDLDLYTHSMTAGFARTMKVVLRRARQMVADSHADSANYLCSFTGLGQM
NBM50_RS05530/WP_233504294      SRIQSGDRSLEVDLDLYTHSMTAGFARTMKVVLRRARQMVADSHADSANYLCSFTGLGQM
NBM50_05530/MCS4509724          SRIQSGDRSLEVDLDLYTHSMTAGFARTMKVVLRRARQMVADSHADSANYLCSFTGLGQM
DFQ15_105107/PYE78748           SRIQSGDRSLEVDLDLYTHSMTAGFARTMKVVLRRARQMVADSHADSANYLCSFTGLGQM
DFQ15_105106/PYE78747           SRLQSGDRSLDTELDFYTYGSNANFVETMKTTLRLARHMVGDSHGDSVNNPSRVTGLGHT
DFQ15_RS17380/WP_199399817      SRLQSGDRSLDTELDFYTYGSNANFVETMKTTLRLARHMVGDSHGDSVNNPSRVTGLGHT
NBM50_RS05540/WP_259292613      SRLQSGDRSLDTELDFYTYGSNANFVETMKTTLRLARHMVGDSHGDSVNNPSRVTGLGHT
NBM50_05540/MCS4509725          SRLQSGDRSLDTELDFYTYGSNANFVETMKTTLRLARHMVGDSHGDSVNNPSRVTGLGHT
DFQ15_RS17300/WP_199399909      SRLRSGDRSLDTELDCYTYGKSVSFAKTMKTTLRRARQMVGDSHGDSVNNSSRVTGLGHT
DFQ15_1541/PYE72703             SRLRSGDRSLDTELDCYTYGKSVSFAKTMKTTLRRARQMVGDSHGDSVNNSSRVTGLGHT
NBM50_RS17175/WP_259292686      ------------------------------------------------------------
NBM50_17140/MCS4511918          ------------------------------------------------------------
```

### Supplementary Data S3

```
DFQ15_RS17525/WP_233504294      GLHLSRPESRGAAEPSA---SVNRYPATPAGSSYSSMFPPTPRGGWPDAPQGSWSELPEV
NBM50_RS05530/WP_233504294      GLHLSRPESRGAAEPSA---SVNRYPATPAGSSYSSMFPPTPRGGWPDAPQGSWSELPEV
NBM50_05530/MCS4509724          GLHLSRPESRGAAEPSA---SVNRYPATPAGSSYSSMFPPTPRGGWPDAPQGSWSELPEV
DFQ15_105107/PYE78748           GLHLSRPESRGAAEPSA---SVNRYPATPAGSSYSSMFPPTPRGGWPDAPQGSWSELPEV
DFQ15_105106/PYE78747           APQMLRLGPRSAAELSASRGSVN-------------------------------------
DFQ15_RS17380/WP_199399817      APQMLRLGPRSAAELSASRGSVN-------------------------------------
NBM50_RS05540/WP_259292613      APQMLRLGPRSAAELSASRGSVNQYPGTSVASSYSSTFPPAPPEVLTQPPQDSPSALPET
NBM50_05540/MCS4509725          APQMLRLGPRSAAELSASRGSVNQYPGTSVASSYSSTFPPAPPEVLTQPPQDSPSALPET
DFQ15_RS17300/WP_199399909      APQMLRLGPRSAAELSASRGSV--------------------------------------
DFQ15_1541/PYE72703             APQMLRLGPRSAAELSASRGSV--------------------------------------
NBM50_RS17175/WP_259292686      ------------------------------------------------------------
NBM50_17140/MCS4511918          ------------------------------------------------------------

DFQ15_RS17525/WP_233504294      SSSTFGDLESLDSGHPDGGRESYWDALNQEATGPWHMASSRASVPTFDEGDTGSGWQHGA
NBM50_RS05530/WP_233504294      SSSTFGDLESLDSGHPDGGRESYWDALNQEATGPWHMASSRASVPTFDEGDTGSGWQHGA
NBM50_05530/MCS4509724          SSSTFGDLESLDSGHPDGGRESYWDALNQEATGPWHMASSRASVPTFDEGDTGSGWQHGA
DFQ15_105107/PYE78748           SSSTFGDLESLDSGHPDGGRESYWDALNQEATGPWHMASSRASVPTFDEGDTGSGWQHGA
DFQ15_105106/PYE78747           ------------------------------------------------------------
DFQ15_RS17380/WP_199399817      ------------------------------------------------------------
NBM50_RS05540/WP_259292613      SSSTFGELEPLESGNHHGGRGLDWDALNQEVTGPWHMADPRPSIPAFNEDDTGTQWQHGT
NBM50_05540/MCS4509725          SSSTFGELEPLESGNHHGGRGLDWDALNQEVTGPWHMADPRPSIPAFNEDDTGTQWQHGT
DFQ15_RS17300/WP_199399909      ------------------------------------------------------------
DFQ15_1541/PYE72703             ------------------------------------------------------------
NBM50_RS17175/WP_259292686      ------------------------------------------------------------
NBM50_17140/MCS4511918          ------------------------------------------------------------

DFQ15_RS17525/WP_233504294      QPAPPWLVQRGVHDQQIVRIRDLEYRVVANPGTSNAWGESQLFLYPHIRGG
NBM50_RS05530/WP_233504294      QPAPPWLVQRGVHDQQIVRIRDLEYRVVANPGTSNAWGESQLFLYPHIRGG
NBM50_05530/MCS4509724          QPAPPWLVQRGVHDQQIVRIRDLEYRVVANPGTSNAWGESQLFLYPHIRGG
DFQ15_105107/PYE78748           QPAPPWLVQRGVHDQQIVRIRDLEYRVVANPGTSNAWGESQLFLYPHIRGG
DFQ15_105106/PYE78747           ---------------------------------------------------
DFQ15_RS17380/WP_199399817      ---------------------------------------------------
NBM50_RS05540/WP_259292613      QAAPPWLVQRGVYEGQIIRIRDLEYRVAAAPGTSNAWGEPQLYVYPRLRGG
NBM50_05540/MCS4509725          QAAPPWLVQRGVYEGQIIRIRDLEYRVAAAPGTSNAWGEPQLYVYPRLRGG
DFQ15_RS17300/WP_199399909      ---------------------------------------------------
DFQ15_1541/PYE72703             ---------------------------------------------------
NBM50_RS17175/WP_259292686      ---------------------------------------------------
NBM50_17140/MCS4511918          ---------------------------------------------------
```

### Supplementary Data S3

**Multiple alignment of 500-bp upstream regions of the most downstream annotated translational start codon of OG_1116 (in green), OG_1115 (in blue) and OG_3308 (in red). Annotated start codons are highlighted in green. Putative Shine-Dalgarno boxes are underlined.**

```
OG_3308        CTTCGATCAAGGCATGGCCGGTCAGGCGCTGGCCCGAT-ACGCCGTAGGGGTGACCCGAG
OG_1115        -------------ATGGACCGCAGGCAGATAACGCAAC-AGGCCGCAACAGGCGCCAG--
OG_1116        --------------GGGGAAGGAAGGTGTTGCCGCAGTCGCGCACCCCCTGCATCCTGCC
                              **   *   *  * *  * *      **       *   ** *

OG_3308        GCATGCCAAGCGCAAGGAATAAAGCGACCATAACCGACACTTGTACAAGACGCGCCATCT
OG_1115        ------CGAGAGCGAGGCGACATGAATGTCGATGCGACGCTCGAAGCGAATGCGCAG-CT
OG_1116        GCCGATCGCGGGCACCTCGTCATGA---TAGAAGCACCGCCCG--GCTGGCCCGCCGCTT
                     *  * **        * *       *  *  * *  *         ***    *
OG_1115                             MetAsnValAspAlaThrLeuGluAlaAsnAlaGln-Le

OG_3308        CATCGAAAACATCTTCGCCCGCCTGAAGCG---GTACTGCGC-----TAGTTCCATATGC
OG_1115        CGCCGAAAGC-----TGCCAGCCAGGCGTGCGTGCGCTCAAC-CACCCAGCGATGGCGGC
OG_1116        TCGCGC---------CGCCGACGCAGCGCCCG-GTGCTCCACAAAAACAGCGCCCTTAGT
                  **           ***  *     *     *  **   *      **        *
               uAlaGluSer-----CysGlnProGlyValArgAlaLeuAsn-HisProAlaMetAlaAl

OG_3308        CACGACAAG-----ACGGCGCGCAACCTCCTCGGGGCCATTCATTTCGCGGCGACGATTG
OG_1115        CAATAGAGACTTCCACAGCGTGCGCCC-CGTCGCAACGAGGATTCAGGCGAGGAGGATGT
OG_1116        AAGTAGAGACTTCCACAGCGTGCGCC--CGTTGCAGCGAGGATTCGGGCGAGGAGGATGT
                *  * *       ** *** **  *  * * *   * *    *   ***  ** ***
               aAsnArgAspPheHisSerValArgPr-oValAlaThrArgIleGlnAlaArgArgMetL

OG_3308        TGCGGTTTGCCCGTCCCTATGACCAACGCTTTCATA-----------GATTATTTATTGA
OG_1115        TACGGTTTACCGA-----AGAACCGACGCCTACA-GCCTCAAACAGGCATAACTCACCAT
OG_1116        CACGGTTCACCGA-----AGGACCAATGCCTACGCACCCTGAACGGGCATAACTCACTAT
                 *****  **       *  *** * ** * *               ** * * *
               euArgPheThrGl-----uGluProThrProThr-AlaSerAsnArgHisAsnSerProL

OG_3308        TGACACGC-----------CCAGGTCAC-CACTGTTGCAGTTTAAAATTTTCCGTTCCAG
OG_1115        TATCAGGCTCGGTGGCGCGCCGGGTCACTCACCGTTGCAGTTTGAAATTTACCGTTCCAG
OG_1116        TATCAGGCTCGGTGGCGTGCCGGGTCACTCACGGTTGCAGTTTAAAATTAACCGTTCCAG
               *  ** **           ** ****** *** ********** *****  *********
               euSerGlySerValAlaArgArgValThrHisArgCysSerLeuLysPheThrValProA

OG_3308        CAAACCGAAATAATGGACTAGCATACGAAATAAAATGATTCGGCCCGCCCCCCGGAAGTA
OG_1115        CAAACCGAAATAATGGGCTGGCATACGAAATAAAAAGATACGGCCCG-CCCCCGCAGGTA
OG_1116        CAAACCGAAATAATGGGCCGGCATCCGAAATAAAAAGATCCGGCCCG-CCCCCGCAGGTA
               **************** *  **** ********** *** ******* ****** * ***
               laAsnArgAsnAsnGlyLeuAlaTyrGluIleLysArgTyrGlyPro-ProProGlnVal

OG_3308        GACTTTTTGTTCAAAGCATGCAAGGATCAGATTTAGAAATTCACATTAATAATACCATCG
OG_1115        GATTTTTCCTTCAAAGCATGGAAGGATCGGAGTTGGAAATTCACAGTCAGAATACCATCG
OG_1116        GATTTTTACTTCAAAGCATGAAAGGATCGGAGTTGGAAATTCACATTCAGAATACTATCG
               ** ****  *********** ******* ** ** ********** * * ***** ****
               AspPheSerPheLysAlaTrpLysAspArgSerTrpLysPheThrValArgIleProSer

OG_3308        AGGCATAAGATGAAAGATCTTTATCCATTCGGTACTGGTACTTCGTGGACACACGACTTG
OG_1115        AGGCATGAGATGAAAGATCTTTATCCATTCGCTACTGGTACCGCGTGGACGCACGACTTG
OG_1116        AGGCATGAGATGAAAGATCTTTATCCATTCGCTACTAATACCGCGTGGACGCACGACTTG
               ****** ************************ ****  ***  ******* *********
               ArgHisGluMetLysAspLeuTyrProPheAlaThrGlyThrAlaTrpThrHisAspLeu
                        MetLysAspLeuTyrProPheAlaThrAsnThrAlaTrpThrHisAspLeu
                        MetLysAspLeuTyrProPheGlyThrGlyThrSerTrpThrHisAspLeu
```
